# Harmonized sample metadata and processed cortical and hippocampal transcriptomes from wild-type and Alzheimer’s disease model mice

**DOI:** 10.64898/2026.11.13.766595

**Authors:** Haruto Ijiri, Hirofumi Kariyayama, Haruka Ozaki

## Abstract

Public mouse brain single-cell and single-nucleus RNA sequencing (sc/snRNA-seq) datasets are difficult to reuse across studies because sample annotations are heterogeneous and incomplete. We harmonized sample-level metadata for 2,005 cortex and hippocampus samples from 389 studies, covering wild-type (WT) controls and Alzheimer’s disease (AD)-related mouse models. Usable count data from 1,863 samples in 360 studies were uniformly processed into 12,628,942 cell and nuclear profiles with cell-type annotations. Expression-based indicators were consistent with the annotated sequencing protocol and sex, and integrated microglial profiles recovered established AD-associated signatures. To illustrate cross-study use, we asked whether microglial AD–WT expression differences varied with age and between cortex and hippocampus, using covariate-adjusted expression reconstructed with Kanade. Genes separated into groups with predominantly negative or positive AD–WT contrasts, including a group whose age-associated contrasts differed between cortex and hippocampus. These findings demonstrate the utility of harmonized metadata for investigating age- and region-dependent AD-associated transcriptional differences across studies. The resource supports explicit, reproducible cohort definitions across public mouse brain datasets.

## 1 Introduction

Single-cell and single-nucleus RNA sequencing (sc/snRNA-seq) has resolved extensive cellular diversity in the mouse brain, including transcriptomic cell types distributed across distinct anatomical regions [1, 2]. Public studies span development, aging, genetic perturbations and disease models, creating opportunities to examine biological variation beyond the design of an individual experiment. In Alzheimer’s disease (AD) research, these approaches have identified disease-associated microglial states and age-, sex- and genotype-related differences in cellular responses [3–5]. Combining such experiments requires expression data together with sufficiently detailed descriptions of what each sample represents. AD-related mouse datasets often encompass diverse ages, genotypes, and experimental conditions [6, 7], highlighting the importance of preserving detailed sample-level metadata.

Sample metadata are a central obstacle to this reuse. Repositories such as the Gene Expression Omnibus (GEO) link expression data to study and sample records, but descriptions of tissue, age, sex, genotype and experimental manipulation vary in vocabulary and detail [8]. Reporting guidance for single-cell experiments emphasizes biological material, experimental design and processing information, while incomplete annotations remain a barrier to data reuse [9, 10]. For example, AD-related samples can carry different disease-associated alleles and additional interventions, and an age mentioned in a protocol can refer to treatment initiation rather than tissue collection. Sample-level interpretation is therefore necessary to avoid combining biologically distinct conditions under a common study label.

Existing resources address complementary aspects of this problem. scREAD, ssREAD and SCAD-Brain organize AD-focused datasets [11–13], while CZ CELLxGENE Discover provides ontology-based metadata standardization and scBaseCount combines uniform read processing with AI-assisted annotation [14, 15]. In scBaseCount, disease labels are extracted at the study level and can propagate to control samples. Its current documentation also describes a per-sample confidence audit to flag potentially heterogeneous disease assignments [16]. These resources motivate complementary curation of collection age, genotype, treatment and model at the individual sample level.

In this study, we developed a harmonized sample-metadata resource linked to processed sc/snRNA-seq data from publicly available mouse cortex and hippocampus datasets, covering WT controls and AD-related models. We manually curated 2,005 GEO samples (GSMs) from 389 studies and processed usable counts from 1,863 samples, yielding 12.6 million retained cell and nuclear profiles. We assessed the consistency of selected annotations with expression-derived indicators and examined whether integrated microglial profiles recovered established AD-associated signatures. Finally, we investigated how microglial AD–WT expression differences vary with age and brain region while controlling the technical heterogeneity that complicates cross-study transcriptomic analyses [17–20].

## 2 Materials and Methods

We distinguish GEO series (GSEs), curated sample records (GSMs, including explicitly marked placeholders) and sequencing libraries used for processing. A library usually corresponds to one GSM, but GSMs containing identifiable independent libraries were split into GSM_k entries. Library counts therefore differ from GSM counts. Supplementary Methods describe additional technical details.

### Mouse sc/snRNA-seq data curation

Candidate studies were assembled through successive searches of GEO DataSets using NCBI E-utilities on 30 September 2026, together with studies identified through ssREAD [12]. The search query was (sequencing OR transcriptome OR RNA-seq) AND (hippocampus OR cortex) AND “Mus musculus”[Organism] AND 2020[PDAT]:3000[PDAT]. Study and sample descriptions were retrieved with GEOquery [21]. GSMs from GSE93421, GSE95315, GSE99888 and GSE104323, which deposited individual cells as separate GSMs, were regrouped into 14 sample-level units (local identifiers beginning GSM000). The selection flow is reported after this regrouping, whereas Gemma screening was performed on the original GSM inventory.

### Metadata annotation

#### Annotation fields and eligibility criteria

Each GSM was annotated for brain region, sex, age, strain, sc/sn protocol, 10x platform, model, genotype and treatment. The metadata schema is provided in Supplementary Table S4. Samples were retained when they were derived from mouse, profiled by a 10x Genomics sc/snRNA-seq assay, sampled from cortex or hippocampus, and assigned to a WT or AD-related model group. Count data availability was recorded separately from biological eligibility.

#### Primary annotation by GPT-4o

OpenAI GPT-4o (gpt-4o, temperature 0) was used to pre-fill annotation fields using GSM characteristics, title, source name, and growth, treatment, and extraction protocols (Supplementary Fig. S1). Six separate requests extracted brain region, sex, age, strain, sequencing strategy/sc-sn protocol/platform, and genotype/treatment, and a seventh request inferred the experimental-model label from the extracted strain, genotype, and treatment. The prompts are shown in the Supplementary Methods (Prompt S5).

The pre-filling procedure produced usable preliminary annotations for 94,060 GSMs, designated cohort 1. The remaining GSMs lacked usable preliminary annotations and were assigned to cohort 2 for Gemma screening. Cohort 2 contained 66,165 unique sample identifiers after re-grouping individual-cell deposits, using the same counting unit as the selection flow. The reasons for missing preliminary annotations could not be established for all GSMs. GPT-4o outputs were treated as suggestions, and inclusion decisions and retained metadata were established by manual review.

#### Screening of cohort 2 with Gemma

To reduce the burden of manual annotation of cohort 2, we used Gemma 4 31B-it (run locally with vLLM, greedy decoding) to classify each GEO sample (GSM) as potentially eligible or ineligible from its study-level and sample-level GEO metadata. The prompt specified three sample-level criteria: (i) 10x Genomics single-cell or single-nucleus RNA-sequencing, (ii) cortex or hippocampus tissue (including subregions), and (iii) a wild-type/control or Alzheimer’s disease (AD)-related condition. Gemma returned a binary label together with criterion-level judgments, a confidence value and failure-reason labels.

The prompt was optimized automatically against a class-balanced reference set of 3,402 manually annotated cohort-1 GSMs, split by GSE into training, validation and held-out test sets. Over 15 rounds, a separate LLM (Claude Opus 5.5, Anthropic) revised Gemma’s instructions in response to errors on the training set only, and the final prompt was chosen by validation *F*_1_, preferring simpler prompts among near-optimal candidates (Supplementary Fig. S2a). On the held-out test set (*n* = 676), the frozen prompt reached precision 0.976, recall 0.959 and *F*_1_ = 0.967 (Supplementary Fig. S2b). The split sizes, optimization procedure, selection rule and all prompts are given in the Supplementary Methods (Prompts S1–S4).

Gemma predicted 478 of the 66,165 unique cohort-2 GSMs (after regrouping individual-cell deposits; Fig. 1a) to be eligible. All 2,883 GSMs from the 117 GSEs containing at least one positive prediction, including 2,405 individually negative predictions, were manually reviewed. The remaining 307 GSEs had no positive prediction and were not reviewed, so their false-negative count is unknown. Original-accession counts are given in the Supplementary Methods.

**Figure 1:**
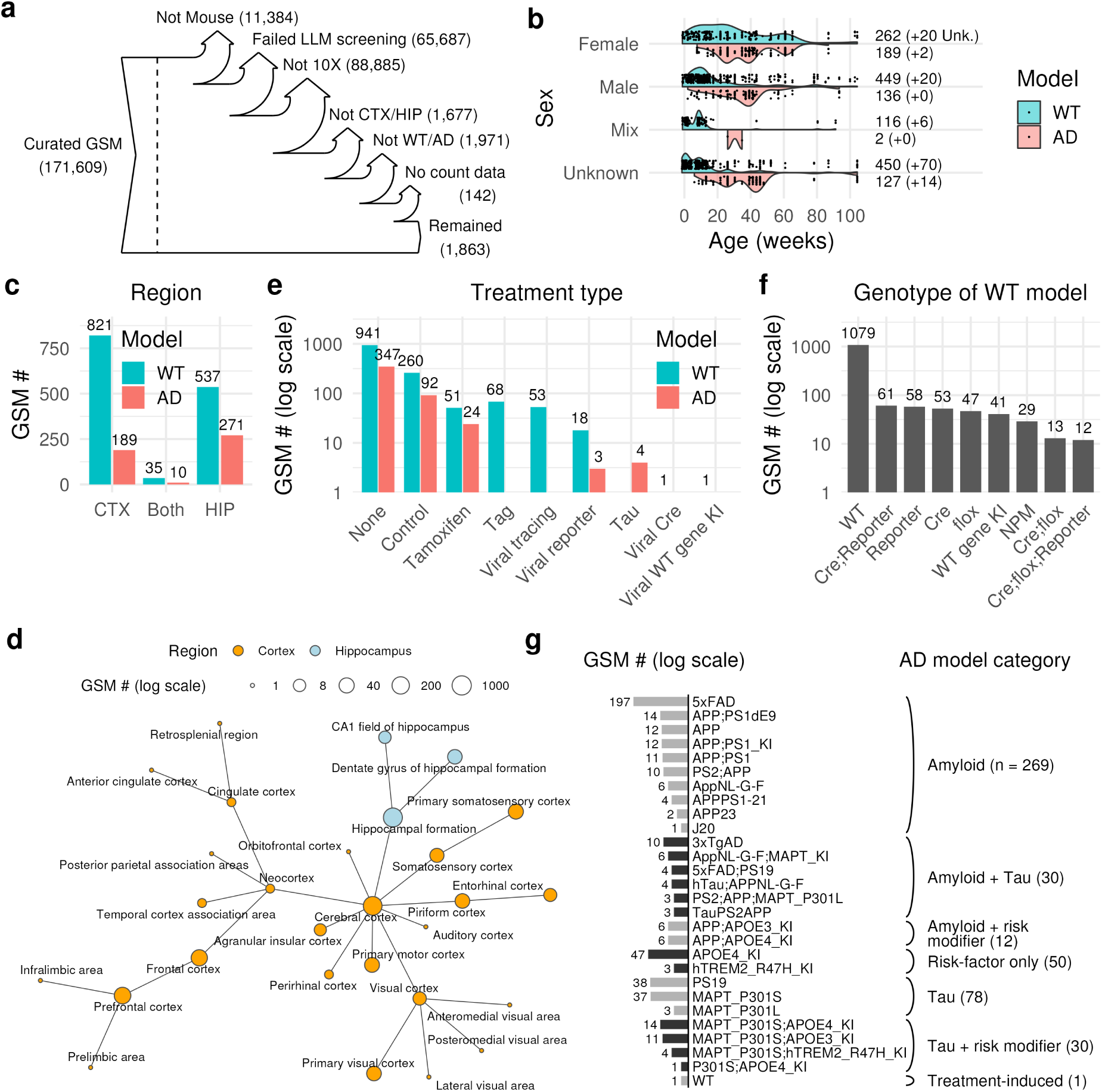
Construction and composition of the harmonized metadata resource. (a) Screening and selection flow from the GEO inventory to the metadata and count-available cohorts. (b) Age distributions by sex and WT/AD model group. Numbers give GSMs with known age and, in parentheses, additional GSMs with unknown age. (c) Numbers of GSMs by mesoregion and model group. Both indicates a combined cortex/hippocampus assignment. (d) UBERON representation of the curated regions, with node sizes indicating record counts on a logarithmic scale. (e) Standardized treatment categories. None includes some unknown or unreported treatment descriptions. (f) Standardized genotypes of the WT/control group. NPM: non-phenotypic mutant. (g) Standardized genotypes of the AD group, organized by broad model category. Panels (b)–(g) describe the 1,863 count-available GSMs.

#### Manual curation rules

One curator annotated the retained GSMs using GEO sample descriptions and, when needed, associated publications and supplementary files. For selected studies, information from the ENCODE portal was also consulted.

##### Brain region

Anatomical descriptions were mapped to UBERON terms [22] at the finest supported resolution. When the description was more specific than an available term, the corresponding parent term was retained. Multiple regions were preserved as multiple ontology identifiers rather than reduced to one dissection label. In the broader annotated inventory, 67 GSMs with ambiguous tissue descriptions were classified as undetermined. To determine mesoregion, UBERON parent–child relations and curated anatomical labels were used to assign broad Cortex and Hippocampus categories. GSMs mapping to both groups were labeled Cortex;Hippocampus (Both).

##### Sex

Sex was recorded as Female, Male, Mix, or Unknown. Mix denotes a sequencing sample pooled from animals of both sexes. A pool of animals of one sex retained that sex label. When a study included both sexes but did not identify the sex of an individual sample, that sample was labeled Unknown. For GSE185862, library-specific sex information was used after the deposited GSM-level pools were divided into sequencing libraries.

##### Age

Age at sample collection was expressed in weeks. Days, months, and years were converted as *d*/7, 365*m*/(12 *×* 7), and 365*y*/7, respectively. Postnatal day 0 was set to zero, and embryonic day *e* was represented as (*e −* 21)/7 weeks, giving negative values for embryonic samples. The midpoint was used when a bounded age range was reported. When the collection age was given as age at treatment plus a subsequent interval, the two intervals were summed. Descriptions such as “adult” were left Unknown when no numerical age could be established, even from the publication.

##### Platform

Samples were labeled 10X when processed using 10x Genomics 3*^r^* or 5*^r^* gene-expression chemistry. Other assay types, including 10x ATAC-seq, Flex, Visium, Xenium, Smart-seq, ATAC-seq, and Drop-seq, were not included in this category.

##### sc/sn protocol

Samples were labeled sn when isolated nuclei were profiled and sc when dissociated whole cells were profiled. When GEO descriptions disagreed (*e.g.*, a sample title indicated snRNA-seq whereas the protocol mentioned a single-cell suspension), information in the associated publication was used to resolve the discrepancy.

##### Strain

Mouse genetic backgrounds, such as *C57BL/6*, *B129*, and *CD1*, were annotated from GEO descriptions or the associated publications.

##### Genotype

Genetic information was manually annotated from GEO descriptions and associated publications. The Genotype field retained the reported genetic constructs, including disease-associated alleles, gene knockouts and knock-ins, Cre drivers, floxed alleles, and reporter constructs. When the genotype could not be determined from the available sources, it was labeled Unknown. *Treatment.* Experimental treatments and relevant procedural conditions were manually annotated from GEO descriptions and associated publications. The Treatment field retained specific descriptions of interventions, including pharmacological treatments, vehicle administration, viral injections, tamoxifen induction, and other experimental procedures or contexts. Control conditions, the absence of a reported treatment, and insufficient treatment information were recorded.

##### Model

Experimental models were manually annotated by considering genotype, treatment, and reported experimental context together. Samples were assigned to WT, AD, other experimental-model categories, or Unknown when the available information was insufficient to establish a model classification. In particular, whole-body or conditional genetic modifications, such as gene-specific knockouts, were not automatically interpreted as disease models or WT controls and were generally assigned to Unknown unless the experimental context supported a more specific classification.

WT represented wild-type animals and appropriate non-disease comparison groups, including some Cre, floxed, reporter, and non-AD knock-in genotypes. Thus, WT was an analysis label and did not necessarily indicate genetically unmodified or untreated animals. AD encompassed amyloid- and tau-associated models, their combinations with AD risk-associated alleles, risk-factor models such as human *APOE4* and *TREM2* R47H knock-ins, and selected tau-administration models. Samples assigned to other diseases, injuries, or experimental perturbations were excluded from the WT/AD collection.

For visualization, genotype and treatment annotations were grouped into broader descriptive categories that are separate from the manually annotated fields (Supplementary Table S4). In particular, the display category None combines several unspecified or unreported treatment.

#### Final collection

The final curated metadata collection comprised 2,005 GSMs from 389 GSEs, including 1,511 WT and 494 AD models.

#### Count acquisition

Forty-nine GSMs lacked usable count data, including absent or normalized-only count matrices, and 93 could not be assigned reliably to GSMs from the deposited barcodes or metadata. Thus, usable, sample-assignable count data were obtained for 1,863 of the 2,005 curated GSMs from 360 GSEs.

Counts for 414 of these GSMs were regenerated from sequencing reads with scRecounter [15], using GeneFull counting for every sample. In scRecounter, sequencing chemistry was determined from the first 1,000,000 SRA spots of each sequencing run by evaluating chemistry-specific barcode whitelists. The chemistry yielding the highest fraction of valid barcodes was selected, requiring a whitelist match rate of at least 30%, following the approach described for scBaseCount. Read alignment and quantification were performed using STARsolo (STAR v2.7.11b). Author-provided count matrices in GEO supplied the remaining 1,449 GSMs. Cell calling was determined separately from the barcode-count distribution, as described below.

Reader implementations, sample-assignment mappings for merged series-level objects, and feature harmonization to mouse gene symbols are described in the Supplementary Methods, and the source code is listed under Data and code availability. Most matrices contain UMI counts, but the GSE314059 long-read matrices are deposited as read-count matrices; count totals are therefore used as the general term for quantities computed across the collection.

Twenty-four GSMs contained multiple sequencing libraries and were divided into 167 library-level samples (GSM_k) before doublet detection. Five placeholder GSMs from GSE99888 and GSE104323 yielded ten libraries, two GSMs from GSE156201 yielded six, and 17 GSMs from GSE185862 yielded 151. Together with the unsplit GSMs, these yielded 2,006 processing libraries from 1,863 GSMs. Metadata were inherited from the parent record except where library-specific information was available, including the sex of GSE185862 libraries.

#### Cell calling, quality control and doublet removal

Matrices with more than 1,000 barcodes carrying fewer than 100 counts were treated as unfiltered for cell calling, a criterion met by 531 libraries. For these libraries, nonzero-count barcodes were processed with STAR 2.7.11b --runModesoloCellFiltering--soloCellFilterEmptyDrops_CR, using its Cell Ranger 3 parameters, including an expected cell number of 3,000 and a false-discovery rate of 0.01 [23]. For the other libraries, all nonzero-count barcodes were retained without an additional total-count threshold.

Low-quality cells were removed using data-driven quality control (ddqc) [24], with thresholds defined within preliminary clusters. Clustering used Scanpy [25], library-size normalization to 10,000 counts, log transformation, 2,000 highly variable genes, scaling, PCA, a ten-neighbor graph and Leiden clustering at the default resolution of 1. Within each cluster, the ddqc configuration applied two scaled median absolute deviations to low total counts, low detected-gene counts and high mitochondrial or ribosomal count percentages, without fixed count thresholds or an additional mitochondrial-percentage cap. Mitochondrial and ribosomal genes were taken from the mouse reference supplied with the ddqc implementation.

Scrublet [26], as implemented in Scanpy, was applied to cells passing ddqc with an expected doublet rate of 0.05, a simulated-to-observed ratio of 2, 30 principal components and automatic threshold selection. When automatic threshold selection returned no threshold, the implementation retained all cells as singlets.

#### Cell-type annotation

Cell types were assigned independently within each library using a Python implementation of the positive-marker scoring procedure of CellMarkerAccordion (CMA) [27], with its mouse healthy-tissue database. Markers were restricted to the exact terms cerebral cortex (UBERON:0000956), hippocampal formation (UBERON:0002421), or their union according to the curated tissue label. Candidate cell types required at least five markers, and at most 30 weighted markers were retained per type. Singlets were reclustered using the same procedure as for ddqc. CMA combined marker evidence consistency, marker specificity and scaled log-normalized expression to score cells for each candidate type. Marker weighting, cell scoring and cluster assignment, including the exact equations, are detailed in the Supplementary Methods. Clusters failing the assignment criteria, libraries with at most 50 singlets, and libraries without eligible marker sets were labeled Unknown.

Cell Ontology terms [28] were mapped to excitatory neurons (EN), inhibitory neurons (IN), oligodendrocyte precursor cells (OPC), astrocytes (AST), microglia (MG), oligodendrocytes (ODC), vascular cells (VC), or Others. Others includes unassigned cells and perivascular macrophages, which were distinguished from parenchymal microglia [29, 30].

#### Transcriptomic consistency checks for sc/sn and sex annotations

Each library was summarized by the mean expression of the nuclear-enriched long noncoding RNAs *Malat1* and *Neat1* [31] and by its mean mitochondrial count percentage, and these were compared between annotated sc and sn libraries. Libraries lacking a metric were excluded from the corresponding comparison (68 libraries in total). For sex, a metric was the ratio of the library-level mean numbers of detected Y-linked and X-linked genes, using a mouse BioMart-derived reference [32] containing 2,625 X-linked and 1,567 Y-linked gene symbols, and one library lacked a usable ratio. Female- and Male-labeled libraries lying on the opposite side of the mean ratio of Mix libraries were flagged for source review.

An expression-based Y/X indicator, computed from summed log-normalized expression of Y-linked and X-linked genes, was used for libraries with Unknown sex. Sc- and sn-specific Gaussian reference distributions fitted to Male- and Female-labeled libraries gave a continuous probability of Male for each library (Fig. 2e).

**Figure 2:**
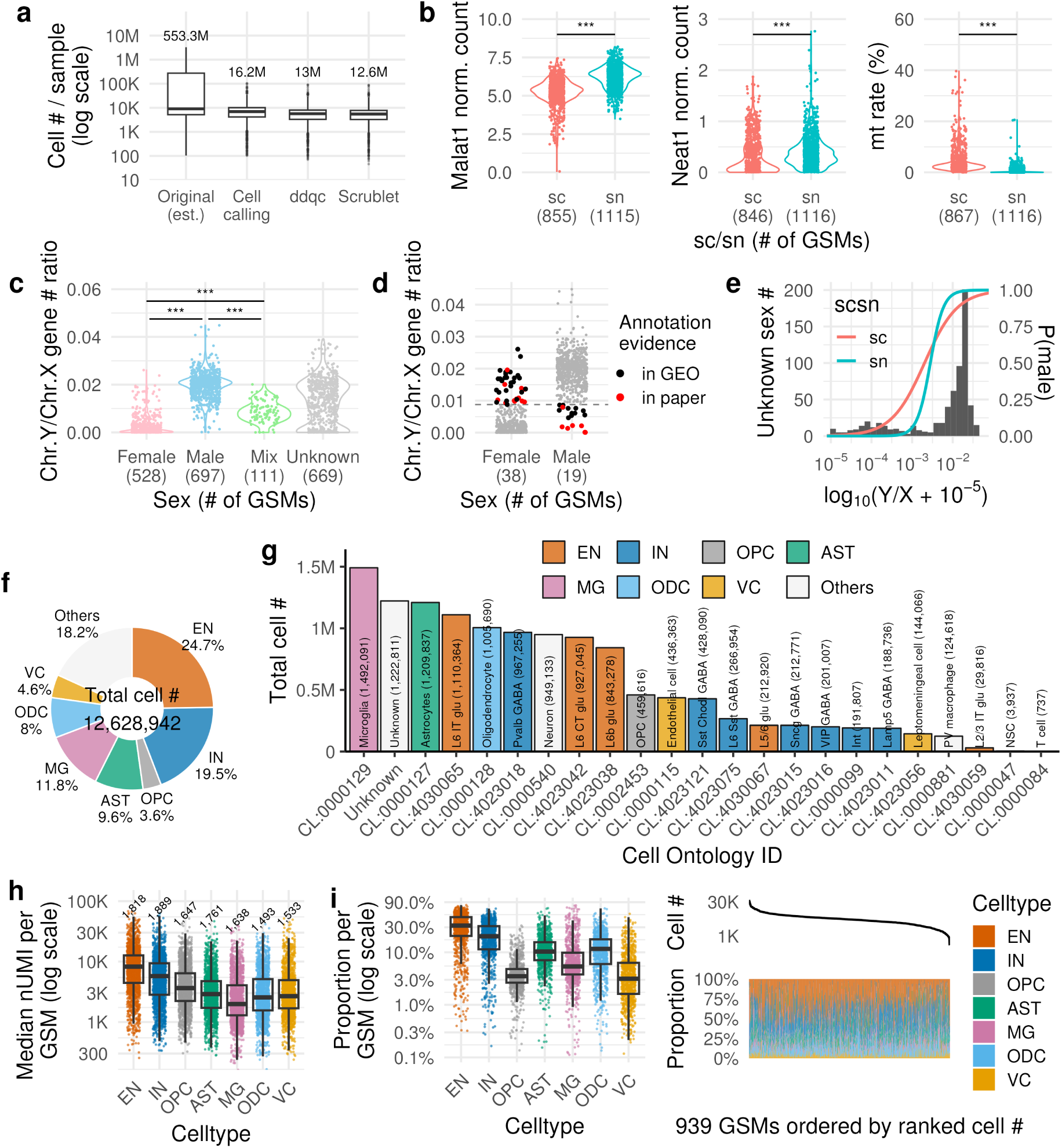
Processing and transcriptomic consistency checks across 2,006 libraries. (a) Nonzero input barcodes and retained profiles at each processing step, with totals above the distributions. The original input count is calculated after feature harmonization and includes author-filtered matrices. (b) Mean cell-level log1p expression per 10,000 counts of *Malat1* and *Neat1*, and mean mitochondrial count percentage, by sc/sn annotation. Numbers in parentheses give usable libraries for each metric. Two-sided Welch’s *t*-tests were used for comparisons. (c) Ratio of library mean numbers of detected Y-linked to X-linked genes, by curated sex. (d) Female- or Male-labeled libraries respectively above or below the mean detected-gene ratio of Mix libraries were flagged for source review. The dashed line indicates this diagnostic threshold, and colors indicate the source supporting the annotation. (e) Distribution of the expression-based Y/X ratio in 669 Unknown-sex libraries, with sc- and sn-specific *P* (Male *| x*) curves on the right axis. Here *x* = log_10_(Y/X+10*^−^*^5^), and Y/X is based on summed log-normalized expression rather than detected-gene counts. Curves were fitted separately by sc/sn protocol to Male- and Female-labeled libraries. (f) Major-class proportions among all 12,628,942 retained cells or nuclei. (g) Counts by assigned Cell Ontology term, including Unknown. (h) Library-level median UMI counts within each major class. Numbers indicate libraries containing the class. (i) Class proportions in the 939 libraries containing all seven major classes, calculated among those seven classes and excluding Others, with libraries ranked by their corresponding total cell count. EN, excitatory neurons; IN, inhibitory neurons; OPC, oligodendrocyte precursor cells; AST, astrocytes; MG, microglia; ODC, oligodendrocytes; VC, vascular cells.

#### Data integration

We integrated observed counts with scVI [19], using the sequencing library (GSM or GSM_k) as the batch identifier. For a global model (Supplementary Fig. S7–S10), at most 100 cells were sampled per library, distributed approximately evenly among the classes present, yielding 200,425 cells from 2,006 libraries. For each class-specific model (Fig. 3), at most 500 cells of the target class were sampled per library. Gene symbols were retained if present in at least 99% of the libraries in the corresponding analysis, with absent genes assigned zero counts. Highly variable genes (HVGs) were selected from raw counts using Seurat v3 ranking [33]. GSEs with fewer than 50 sampled cells or an unsuccessful variance-model fit were omitted from HVG ranking. The global model used 5,000 HVGs, and the class-specific models used 2,000 HVGs. Training used a 90%/10% cell split, early stopping and restoration of the best validation checkpoint. Architecture and optimization settings are provided in the Supplementary Methods. The posterior mean of the latent variables was used as the integrated embedding.

**Figure 3:**
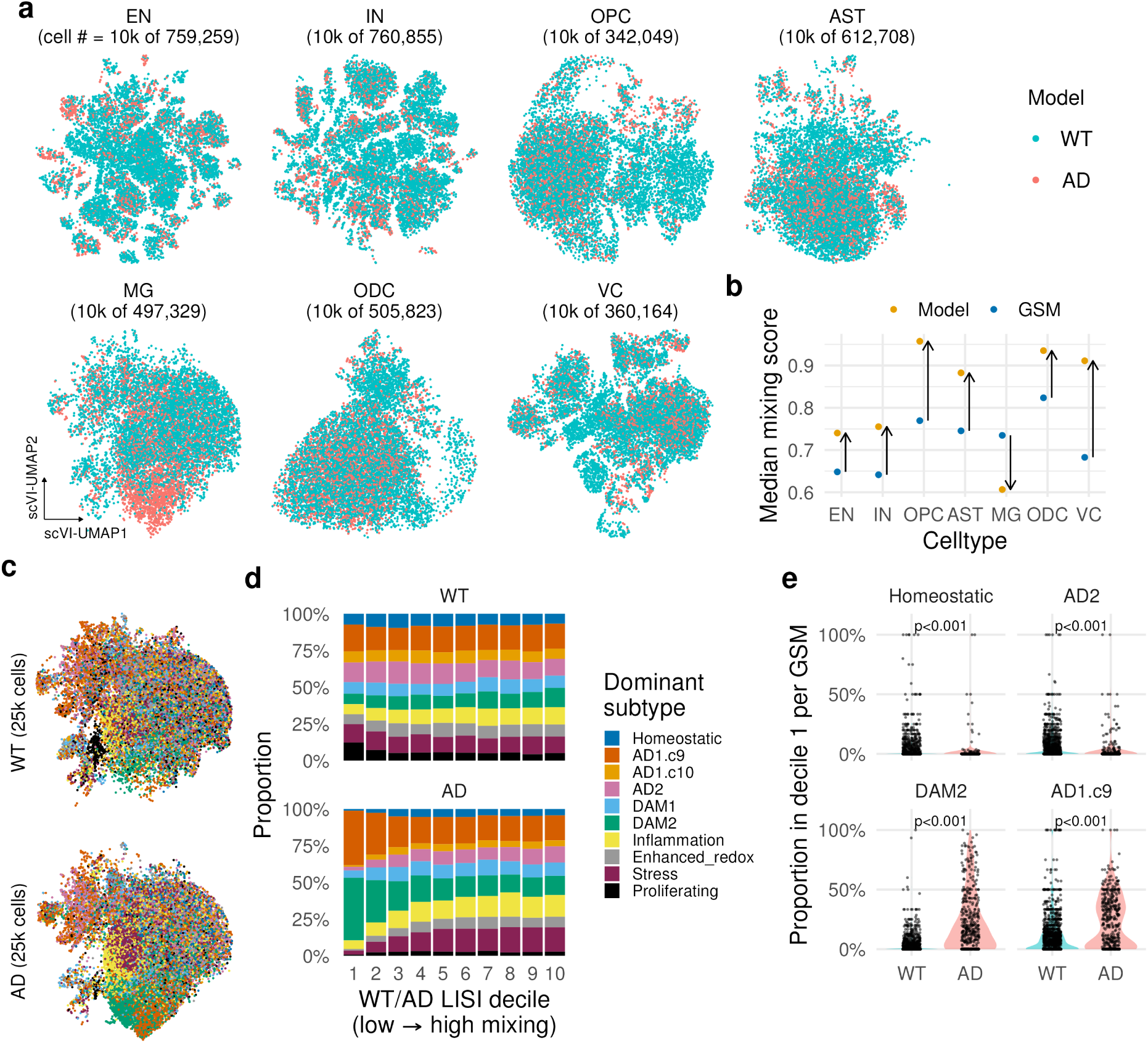
Integration recovers AD-associated microglial gene signatures. (a) Class-specific scVI UMAPs colored by WT/AD Model, displaying 10,000 randomly sampled cells per class from the complete class-specific analysis cohorts indicated in each panel. (b) Median permutation-normalized LISI mixing scores for Model and library identity, calculated using all cells in each class-specific cohort. Arrows indicate the change from library to Model mixing. (c) Microglial scVI UMAPs displaying 25,000 sampled WT and 25,000 sampled AD cells, colored by the highest z-score among the ten displayed signatures without a minimum-score threshold. (d) Dominant-signature proportions within Model LISI deciles for the same displayed subsets and assignment rule as in (c). Deciles were assigned using the complete microglial cohort separately within WT and AD, from least to most mixed. (e) Library-level proportions of four selected dominant signatures in decile 1 (13 candidate signatures, see Methods). Each point is a library. GSM labels in the panels denote processing-library identifiers. Two-sided Welch’s *t*-tests with Bonferroni adjustment across ten comparisons.

For the unintegrated baseline, we normalized the same cells and HVGs to 10,000 counts per cell, applied log1p transformation, and computed principal components without gene-wise scaling, using the latent dimensionality as the number of components. Uniform manifold approximation and projection (UMAP) [34] was computed from a 15-nearest-neighbor graph of each embedding. Class-specific UMAPs displayed 10,000 randomly selected cells per class, whereas mixing statistics used the complete sampled class-specific datasets.

#### Local inverse Simpson’s index and mixing score

To quantify how well each annotation is mixed in an embedding, we computed a mixing score for every cell. LISI (local inverse Simpson’s index) [35] counts how many categories of an annotation, such as library, GSE or sc/sn protocol, are effectively represented among a cell’s 90 nearest neighbors. Because this count depends on the number and proportions of categories, we divided it by the value obtained after randomly permuting the labels, giving a mixing score that is comparable across annotations. A score of zero means that a cell’s neighbors all share its label, and a score of one means that neighbors are as mixed as expected from random labels. High mixing is desirable for technical annotations (library, GSE, sc/sn protocol), whereas for biological annotations (Model, tissue, sex, age category) it is not necessarily desirable, because they represent variation that should be retained. The same procedure was applied to observed-count PCA, scVI latent coordinates and PCA of the Kanade expression layers, excluding Unknown labels and treating Mix as a separate sex category. The LISI calculation, covariate definitions (including operational age categories) and stratification are given in the Supplementary Methods.

#### Microglial gene-signature scoring

Microglia were scored for published gene signatures from Green et al. [36], including the AD1/AD2 signatures of Gerrits et al. [37] and the DAM signatures of Keren-Shaul et al. [3]. Each cell’s score for a signature was the mean log-normalized expression of its up-regulated markers, referenced to WT cells from the same study and expressed as a z-score. Ten signatures with at least five available markers were used for the main displays. Cells were ranked by Model LISI within WT and AD, and signature scores were compared across the resulting deciles. The scoring formula, standardization and both assignment rules are given in the Supplementary Methods.

### Exploratory analysis of age-associated microglial expression

#### Analysis cohort and Kanade training

We used Kanade [38] to explore age-matched AD–WT expression patterns after adjusting the reconstructed representation for study, sc/sn protocol and sex, as an application of the harmonized metadata resource. The analysis cohort comprised 433,128 sampled microglia from 1,373 libraries (1,345 GSMs and 263 GSEs), including 434 AD and 939 WT libraries with recorded ages spanning 2.0–104.29 weeks. At most 500 microglia were sampled per library with deterministic library-specific seeds. GSE was the technical batch identifier.

The training covariates were Model, log_2_ age, tissue, sex and sc/sn protocol, with a separate unsupervised latent component. The 28-dimensional latent representation contained supervised blocks for each covariate, an unsupervised block and a GSE block (dimensions in the Supplementary Methods). Tissue was coded as cortex = 1, hippocampus = 0 and combined tissue = 0.5. Sex was encoded as Male = 1, Female = 0 and Mix = 0.5. For the 373 Unknown-sex libraries, the continuous probability of Male (Fig. 2e) was used. Continuous covariates were standardized using training cells. The common input comprised 2,000 HVGs selected by within-GSE Seurat v3 ranking [33], excluding genes matching the mitochondrial and ribosomal-symbol filters specified in the Supplementary Methods. Models were trained with three initialization seeds (0, 1 and 2). Latent-block dimensions, loss weights, optimization settings and stopping criteria are provided in the Supplementary Methods.

#### Expression reconstruction and adjustment

The unadjusted reconstruction, R0, retained all posterior-mean latent coordinates. For the adjusted layer, R2, the GSE-associated block was set to zero, the sex block to the midpoint of the Male and Female latent centroids, and the sc/sn block to the mean of the sc and sn centroids weighted by their cell fractions in the analysis input. Model-, age- and tissue-associated coordinates and the unsupervised component were retained. Decoder outputs were converted to library-size-independent expected log-scale expression estimates over the 2,000 HVGs.

Covariate mixing levels were assessed using the same permutation-normalized LISI procedure applied to 30-component PCA representations of the complete raw, R0 and R2 expression layers. Expression fidelity was assessed by within-cell Spearman correlations between layers, across either the 480 genes subsequently selected for pattern analysis or all 2,000 HVGs, and by the paired change in correlation with the observed data between R0 and R2. The raw expression layer used observed counts normalized over the same 2,000 HVGs. The cell-sampling cap, correlation definitions and aggregation are detailed in the Supplementary Methods.

#### Age contrasts and gene-pattern analysis

Adjusted expression was averaged per library (and across the three seeds), then within GSE and equally across GSEs, separately for WT and AD within each tissue and age band. The AD–WT contrast in each supported tissue-by-age stratum was accompanied by percentile intervals from GSE-level bootstrap resampling. A contrast passed an exploratory consistency criterion when its 95% bootstrap interval excluded zero, its direction agreed across all three seeds, and its direction matched that of observed-count pseudobulk. Within-GSE contrasts matched by tissue, age band and sc/sn protocol were calculated as a sensitivity analysis.

The pattern analysis included 480 genes passing the above consistency criterion with |Δ*| ≥* 0.25 in at least one supported cortex or hippocampus stratum. The normalized contrast profiles of these genes across the 14 supported strata were grouped by PCA and Ward clustering into three clusters. GO Biological Process over-representation, followed by Bonferroni correction, was assessed for each cluster and displayed GO terms were selected post hoc for interpretability. Age-band definitions, stratum-support rules, normalization, and GO annotation and testing details are given in the Supplementary Methods. Genes labeled as AD-associated in the displays were taken from the effector-gene nominations of Uffelmann et al. [39] and the ADSP Gene Verification Committee list [40, 41], mapped to mouse symbols; these labels were not used to select genes (provenance in the Supplementary Methods).

#### Statistical analysis

Transcriptomic protocol and sex indicators were compared between annotation groups using two-sided Welch’s *t*-tests on library-level summaries. The microglial dominant-signature comparisons also used Welch’s tests with Bonferroni adjustment across ten comparisons. These comparisons are descriptive because multiple libraries can derive from one study or animal. For the 480 selected genes, cell-level expression (observed log-normalized or Kanade-adjusted R2) in 415,579 microglia from 1,329 libraries was modeled with a linear mixed-effects model, with age, tissue, Model (WT/AD), their interactions and sc/sn protocol as fixed effects and library as a random intercept. P values used GSE-cluster-robust standard errors with Benjamini–Hochberg adjustment. The model specification, robustness analyses and null calibration are given in the Supplementary Methods, and gene counts and null-calibration results in Tables S1–S2.

#### Software and computational provenance

Data collection and figure generation were implemented in R, and count processing and advanced analyses in Python. Computations were run on a Linux server with an NVIDIA A100 80 GB GPU. Software versions are summarized in Supplementary Table S3.

## 3 Results

### A harmonized metadata resource linked to reusable count data

The cumulative inventory contained 2,249 GSEs and 184,385 GSMs (171,609 after regrouping individual-cell deposits), including 160,225 mouse GSMs from 2,103 GSEs. LLM-assisted preliminary annotation and screening, followed by manual curation, yielded 2,005 eligible GSMs from 389 GSEs representing 10x Genomics sc/snRNA-seq of WT/control or AD-related mouse cortex and hippocampus samples (Fig. 1a). Usable, sample-assignable counts were obtained for 1,863 GSMs from 360 GSEs: 1,393 WT/control and 470 AD GSMs, comprising 731 scRNA-seq and 1,132 snRNA-seq samples. The broader annotated inventory also contained other experimental models and anatomical regions that were excluded from this focused resource (Supplementary Figs. S3 and S4).

Among the 1,863 count-available GSMs, age was known for 1,731 (92.9%), spanning embryonic stages to 104.3 weeks, and sex was recorded as Female, Male or Mix for 1,202 (64.5%). The remaining 132 GSMs had unknown age and 661 had unknown sex (Fig. 1b). The broad tissue assignments were Cortex for 1,010 GSMs, Hippocampus for 808 and Both for 45, while the retained UBERON annotations preserved more detailed anatomical information (Fig. 1c,d).

Treatment and genotype annotations exposed distinctions within the broad WT/AD groups (Fig. 1e–g). The largest standardized treatment category was None, containing 1,288 count-available GSMs, including 403 with an Unknown or unknown value in the underlying Treatment field. Other categories retained control conditions and procedures such as tamoxifen administration and viral manipulation. Treatment annotations outside the WT/AD collection were more diverse and included pharmacological, injury and other perturbation contexts (Supplementary Fig. S5). The WT/control group included Cre, floxed and reporter genotypes, illustrating that this analysis label does not imply an unmodified animal. The 470 count-available AD GSMs comprised 28 standardized genotype labels, and 5xFAD accounted for 197 GSMs (41.9%).

### Transcriptomic checks and cell-type annotation across 12.6 million retained profiles

The 1,863 count-available GSMs yielded 2,006 processing libraries after separation of merged libraries. Across the harmonized input matrices, 553,258,489 barcodes had nonzero counts. Cell calling and retention of already filtered inputs yielded 16,186,104 profiles, ddqc retained 12,992,739, and Scrublet retained 12,628,942 putative singlet cells or nuclei (Fig. 2a).

Libraries annotated as sn showed higher *Malat1* and *Neat1* expression and lower mitochondrial count percentages than sc libraries (Fig. 2b, all *p <* 0.001). The ratio of the mean numbers of detected Y-linked and X-linked genes was higher in Male than Female libraries, with Mix libraries intermediate between these groups (Fig. 2c, all pairwise *p <* 0.001). The source-review heuristic flagged 38 Female libraries above, and 19 Male libraries below, the mean ratio of Mix libraries. Their annotated sex labels were supported by explicit GEO or publication descriptions (Fig. 2d). For the 669 Unknown-sex libraries with a usable expression-based Y/X ratio, the protocol-specific Gaussian reference fits assigned 496 (74.1%) a *P* (Male *| x*) *>* 0.5 and 173 (25.9%) a value below 0.5 (Fig. 2e).

CMA identified 22 cell types mapped to Cell Ontology terms, with unassigned cells labeled as Unknown (Fig. 2f,g, Supplementary Fig. S6). EN and IN represented 24.7% and 19.5% of retained profiles, followed by MG (11.8%), AST (9.6%), ODC (8.0%), VC (4.6%) and OPC (3.6%). Others represented 18.2%, including 1,222,811 Unknown profiles (9.7% of the complete collection). Library-level total-count distributions differed among major classes, and the relative abundance of those classes varied substantially across libraries that were broadly similar with profiles reported by Green et al. [36](Fig. 2h,i). Together, these summaries describe the composition of the processed resource and provide internal checks on its protocol, sex and cell-class annotations.

### Integration recovers established AD-associated microglial signatures

We first examined how global scVI integration changed the organization of technical and biological annotations in 200,425 sampled profiles. Relative to the observed-count PCA baseline, median library mixing scores increased by 0.198–0.278 and GSE mixing scores by 0.205–0.356 across the seven major classes (Supplementary Fig. S7). Cell Ontology and major-class mixing remained low, changing from 0.093 to 0.104 and from 0.072 to 0.088, respectively, across labeled cells (Supplementary Fig. S8a). Protocol-associated separation remained substantial: sc/sn mixing increased in every class, but the integrated median scores ranged from only 0.015 to 0.142, with the lowest values in MG and ODC (Supplementary Fig. S8b).

Changes in tissue organization differed among classes. Median tissue mixing increased by 0.254–0.351 in OPC, MG, ODC and VC, compared with 0.021–0.100 in EN, IN and AST (Supplementary Fig. S9a). Sex and age-category mixing increased by 0.143–0.275 and 0.134–0.242, respectively (Supplementary Figs. S9b and S10a). WT/AD mixing increased by 0.184–0.301 in six classes, whereas the increase in MG was 0.042 (Supplementary Fig. S10b). Thus, reduced study-associated organization coexisted with residual protocol, tissue and model structure.

To examine the relatively persistent WT/AD separation in MG, we trained separate scVI models for each major cell class. Microglia were the only class for which median Model mixing was lower than library mixing in this class-specific comparison (Fig. 3a,b). The microglial cohort comprised 497,329 profiles: 321,475 WT and 175,854 AD. Published microglial gene sets were used to assign dominant-signature labels, providing a reference for interpreting the integrated expression patterns (Supplementary Table S8). The least-mixed AD decile contained a greater representation of DAM2- and AD1.c9-dominant profiles (Fig. 3c,d). Within decile 1, library-level proportions of DAM2- and AD1.c9-dominant profiles were higher in AD, whereas Homeostatic- and AD2-dominant profiles were more frequent in WT (all displayed adjusted *p <* 0.001, Fig. 3e). These results recover recognizable AD-associated microglial signatures within cohorts defined by the curated metadata.

### Exploratory analysis of age-associated AD–WT expression patterns

We next asked whether microglial AD–WT expression differences varied across age and between cortex and hippocampus. We used Kanade to construct a representation of expression adjusted for study, sex and sc/sn protocol (R2), and compared age-by-tissue AD–WT contrasts in this representation. The analysis included 1,373 libraries from 263 GSEs: 54 GSEs contained both WT and AD libraries, 181 contained WT only and 28 contained AD only. Age, tissue, protocol and genotype were incompletely crossed (Supplementary Fig. S11). Adjustment increased library, GSE, sc/sn and sex mixing and removed Model, tissue and, to a lesser degree, age structure from the PCA representation, while retaining moderate expression-rank correspondence with the observed data (median library-level Spearman *ρ* 0.417–0.427 for R2 versus 0.491 for R0, across the 480 genes analysed below; Supplementary Figs. S12 and S13a).

We selected 480 of 2,000 highly variable genes with a consistent AD–WT contrast in at least one supported tissue-by-age stratum. Normalized contrast profiles across 14 supported strata yielded three clusters of 203, 178 and 99 genes (Fig. 4b). Cluster 1 had negative mean AD–WT contrasts across the analysed strata and included *P2ry12*, *Tmem119*, *Bin1*, *Clu* and *Mrc1*. Cluster 2 had positive mean contrasts in both tissues across the age bands and included *Apoe*, *Igf1*, *Trem2*, *Ctsb* and *Ank*, consistent with established disease-associated microglial signatures (Fig. 4a–c). GO Biological Process terms in cluster 2 were enriched for immune-system processes, whereas cluster 1 terms included negative regulation of programmed cell death (Fig. 4d). Cluster 3, including *C4b*, *Grm3* and *Cntn4*, showed region-dependent age-associated patterns: its mean cortical contrast decreased from 0.203 at 4–9 weeks to *−*0.024 at 44–65 weeks, whereas its mean hippocampal contrast increased from 0.057 at 4–9 weeks to 0.255 at 78–105 weeks (Fig. 4c). Genes from the AD-associated reference list, compiled from GWAS effector-gene nominations (Supplementary Data 2) [39] and an ADSP-attributed gene list [40, 41], occurred in both clusters 1 and 2. These profiles suggest that AD-associated microglial expression differences are not uniform across genes, ages or brain regions. Several negatively contrasted genes encoded neuronal, oligodendroglial, haemoglobin or vascular-associated transcripts, potentially representing ambient RNA or other-cell contamination.

**Figure 4:**
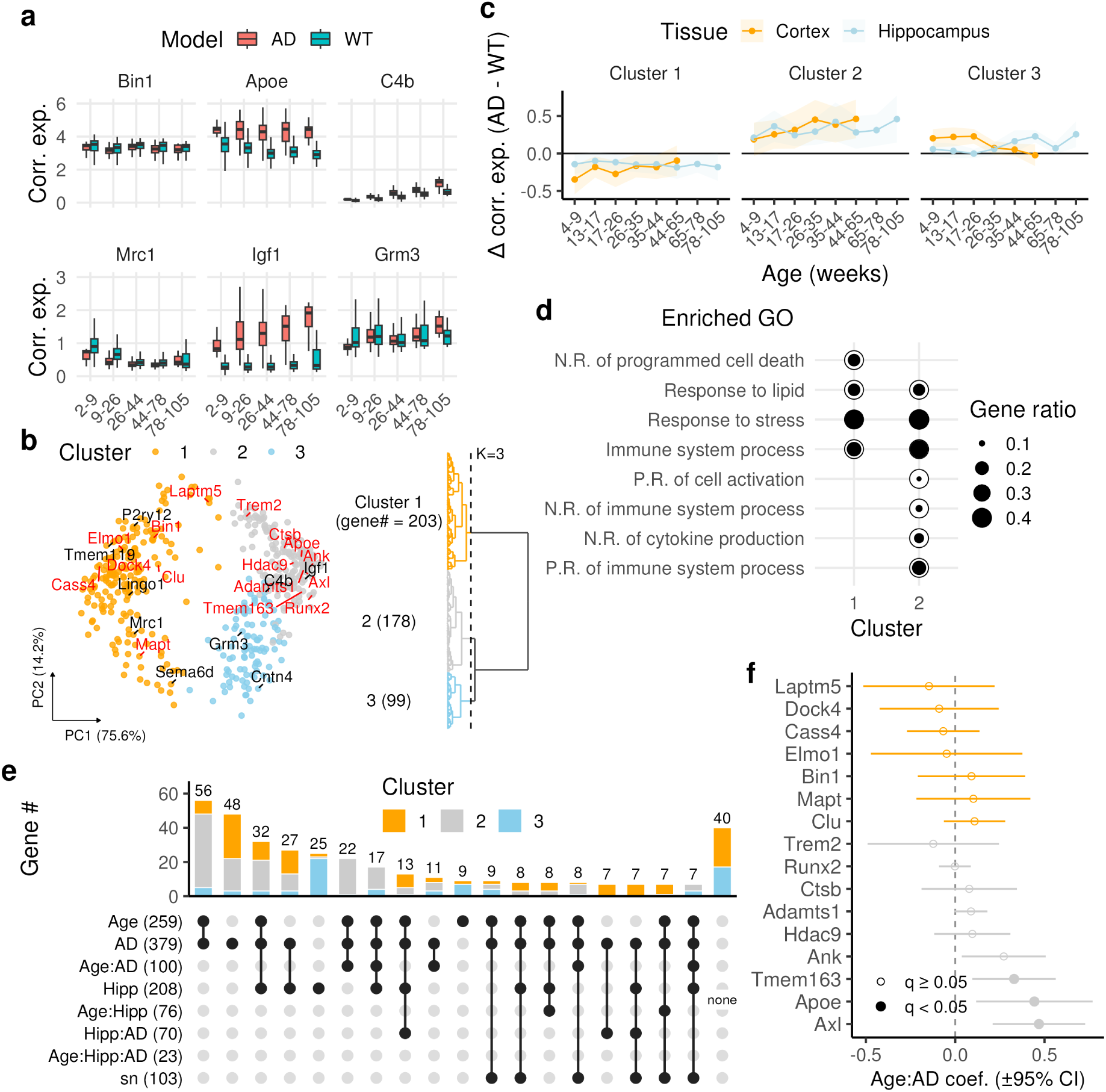
Exploratory age-associated AD–WT differences in reconstructed microglial expression. (a) Library-level mean adjusted expression (R2, seed 0) of *Bin1*, *Apoe*, *C4b*, *Mrc1*, *Igf1* and *Grm3*. All retained tissue groups were pooled for these illustrative boxplots, and adjacent age bands were combined into [2, 9), [9, 26), [26, 44), [44, 78) and [78, 105) weeks. (b) PCA and Ward hierarchy for the normalized AD–WT difference profiles of 480 selected genes across 14 supported tissue-by-age strata. Contrasts used the mean of three training seeds. Red labels denote genes in the AD-associated reference list. (c) Mean AD–WT difference across the genes in each cluster, separated by tissue and original age band. Ribbons show one standard deviation across genes, not uncertainty intervals for a between-Model effect. (d) Selected GO Biological Process terms from the cluster over-representation analysis. Dot area denotes the proportion of genes in the annotated hit set assigned to the term. (e) UpSet representation of overlaps among eight fitted mixed-model terms for the 480 selected genes on the adjusted R2 layer, separated by the displayed R2-derived gene clusters. Bars count genes in exclusive significance combinations; the rightmost column represents genes with no significant term. Significance is *q*_BH_ *<* 0.05 from GSE-cluster-robust *p*-values, adjusted within each model term and expression layer (Supplementary Methods). (f) Estimated log-age-by-AD interaction coefficients for genes matching the AD-associated reference list within the selected 480-gene panel, grouped by the adjusted-expression cluster. Error bars are approximate 95% intervals (*β*^^^ *±* 1.96 SE_GSE_, GSE-cluster-robust), and point style indicates *q*_BH_ *<* 0.05. The coefficient is per unit log_10_(age/20 weeks) and must not be interpreted as a per-week effect. P.R., positive regulation; N.R., negative regulation.

To quantify these patterns with continuous age, we fitted a gene-wise mixed-effects model (LMM) with log-age, Model, tissue, their interactions and sc/sn protocol to the same 480 genes, using observed or R2 expression (415,579 microglia from 1,329 libraries and 255 GSEs; Fig. 4e,f, Supplementary Table S1). An age-by-AD interaction was BH-significant for 100 genes in R2 and nine in observed expression, with six shared, and interaction coefficients from the two representations were only weakly concordant (*ρ ≈* 0.2–0.3). Of the 16 selected genes in the AD-associated reference list, *Axl*, *Tmem163* and *Apoe* had BH-significant age-by-AD interactions in R2, and only *Axl* also in observed expression (Fig. 4f, Supplementary Fig. S13e). The formal age-by-tissue-by-AD term, which corresponds to the regional pattern of cluster 3, was significant for only 23 genes in R2 (none in observed expression), so that pattern is descriptive. Adjusting the expression representation thus materially changed the age-by-AD candidate list. Because this analysis used the same data and genes, and a GSE-block permutation audit gave a null rejection rate of 0.104 for the R2 age-by-AD term (Supplementary Table S2), we regard the R2-specific associations as exploratory.

## 4 Discussion

This study links harmonized sample metadata to processed mouse cortex and hippocampus sc/snRNA-seq data, allowing cohorts to be defined using the experimental context of individual samples (Fig. 1). Its main contribution is to retain collection age, anatomical region, sex, protocol, genotype, treatment and model as separate fields, rather than treating broad disease labels as sufficient descriptions. In particular, WT/control does not necessarily denote an unmodified or untreated animal, and AD-related models differ in their genetic and experimental contexts (Fig. 1e–g). Because these fields remain distinct, the same resource can support differently constrained comparisons without silently collapsing experimental controls into the same category. The resource therefore complements AD-focused databases and larger single-cell infrastructures by enabling more explicit sample selection and examination of heterogeneity within nominally comparable groups [11–15].

The curation process also illustrates why extracting a value from a sample description is not equivalent to interpreting its biological meaning. For example, an age reported in GEO could refer to treatment initiation rather than tissue collection, requiring consultation of the accompanying publication to determine the relevant age. GPT-4o-assisted pre-annotation and Gemma-based eligibility screening helped organize the large candidate inventory, but manual assessment remained central to interpreting the retained GSMs. The Gemma benchmark measured eligibility screening, rather than the accuracy of each annotated field (Supplementary Fig. S2). This distinction matters because study-level descriptions can cover biologically different samples, including treated animals and their controls. Separating genotype, treatment and model was particularly valuable when deciding whether an experimental control belonged in the WT group or whether a disease-associated intervention justified an AD label.

The expression data provided complementary checks on the curated metadata (Fig. 2). Nuclear-enriched transcripts and mitochondrial fractions differed between annotated sc and sn libraries, while sex-chromosome indicators broadly followed the recorded sex groups (Fig. 2b–e). Review of the 57 flagged Female- or Male-labeled libraries identified source support for their annotations, rather than clear evidence of errors. For libraries with unreported sex, protocol-specific expression ratios supplied continuous reference covariates for the exploratory analysis without changing the curated Unknown labels (Fig. 2e). CellMarkerAccordion recovered major brain cell classes and finer Cell Ontology assignments, with marker-score distributions providing internal checks of their consistency (Fig. 2f–i, Supplementary Fig. S6). The broad cell-class abundance and UMI-count rankings are also qualitatively similar to the independent human prefrontal-cortex snRNA-seq observations of Green et al. [36] (their Extended Data Fig. 1l,m). This cross-species resemblance strengthens plausibility but cannot establish assignment accuracy, because species, region, study selection, cell or nucleus capture and classifier methodology all differ. Together, these analyses support the usability of the processed collection without equating transcriptomic agreement with independent ground-truth validation.

Integration provided a biological test of metadata-defined cohorts. Global scVI increased mixing by library and GSE, while sc/sn and regional structure remained evident and varied among cell classes (Supplementary Figs. S7–S10). This residual organization is compatible with known protocol-associated and anatomical differences, and emphasizes that maximum mixing is not necessarily desirable for biological covariates [17, 18]. WT/AD mixing increased less in microglia than in the other major classes, motivating a more focused evaluation (Supplementary Fig. S10b). In the class-specific analysis, microglia retained relatively strong WT/AD separation (Fig. 3a,b). Less-mixed AD neighborhoods were enriched for DAM2- and AD1.c9-dominant profiles, consistent with established AD-associated microglial programs [3, 4, 37] (Fig. 3c–e). The recovery of these signatures supports the biological coherence of the curated WT/AD cohorts. It is not presented as discovery of a new microglial state.

The Kanade application shows that the curated metadata allow cross-study microglial comparisons stratified by AD model, age and brain region (Fig. 4). The negative and positive AD– WT gene groups organize familiar microglial expression programs: cluster 2, containing *Apoe* and *Trem2*, is consistent with established disease-associated microglial signatures [3, 42], and cluster 1 contains homeostatic genes such as *P2ry12* and *Tmem119*. This recovers established biology rather than defining a new cell state. The third group showed region-dependent age-associated patterns that would be difficult to see in an undifferentiated pooled AD–WT comparison, and raises the question of whether specific genes have distinct age-by-AD associations in cortex and hippocampus. These observations make specific gene-by-age-by-region hypotheses worth testing in matched samples or independent cohorts. The comparison of observed and reconstructed expression further shows how strongly candidate age-by-AD associations can depend on adjustment choices. Several limitations and future directions follow from the design of this resource. The search and screening procedure does not establish exhaustive coverage: studies without Gemma-positive GSMs were not fully reviewed, and a single curator annotated the retained samples without an independent second review. Publication evidence was consulted selectively, and unresolved age and sex fields remain (Fig. 1b). Usable or reliably assignable counts were unavailable for 142 eligible GSMs. Author-provided matrices differed in prior processing, a small subset contained read rather than UMI counts, and some retained profiles lacked a confident cell-class assignment (Fig. 2g). Transcriptomic sex indicators are not verified animal-sex labels, and marker-based checks are not independent validation of cell-class calls. More complete sample-level provenance and independent annotation audits would strengthen future versions of the collection.

Kanade analyses carry material limitations. Genotype, age, tissue and protocol were incompletely crossed across public studies, age comparisons are cross-sectional rather than within-animal trajectories, and GSE-level dependence limits inference (Supplementary Fig. S11). Kanade received age, Model and tissue as supervised covariates, so their separation after reconstruction does not independently establish biological fidelity, and representation-specific age and AD differences may reflect decoder behaviour as well as signal. Because interaction coefficients agreed only weakly between representations and the GSE-block null audit was not perfectly calibrated, the R2-specific interaction hits are exploratory and not validated age-by-AD effects. The 480-gene selection and clustering were data-dependent, so cluster patterns, enrichment and hit counts are not independent confirmatory tests.

Broader coverage of disease models and interventions, alongside matched experimental validation, would extend the resource’s utility. Its present value is the transparent link between sample context and reusable transcriptomes, illustrated by recovery of known microglial signatures and the generation of testable age- and region-associated hypotheses (Figs. 3 and 4).

## Supporting information

Supplementary Methods, Tables, and Results

## Funding

This work was supported by JST, NBDC Japan Grant Number JPMJND2402 to H.O., JSPS KAKENHI Grant Number JP23H04149 (Glia Decoding) to H.O., and the RIKEN TRIP Advanced General Intelligence for Science Program (AGIS) to H.O.

## Declaration of competing interest

The authors declare no conflicts of interest for this work.

## Data and code availability

The original datasets are available through GEO and the additional repositories identified by the curated source links. Metadata-curation, count-reader and figure-generation code is available at https://github.com/bioinfo-tsukuba/Brain_Blast.

## CRediT authorship contribution statement

H.I.: Conceptualization, Methodology, Software, Formal analysis, Investigation, Data curation, Data preparation, Visualization, Writing – original draft. H.K.: Data preparation. H.O.: Conceptualization, Resources, Supervision, Writing – review & editing.

## Acknowledgements

We thank members of the Ozaki laboratory at the University of Tsukuba and RIKEN BDR for helpful discussions and constructive feedback.

## Declaration of generative AI in the writing process

During preparation of this work, the authors used ChatGPT (OpenAI) and Claude Code (Anthropic) for drafting, language editing, literature and code review, and checking consistency between the manuscript and analysis outputs. The LLM-assisted data-screening procedure is described separately in the Methods. The authors reviewed and edited the generated material and take responsibility for the final content.

## Notes

### Competing Interest Statement

The authors have declared no competing interest.

## References

[1] Amit Zeisel, Hannah Hochgerner, Peter Lönnerberg, Anna Johnsson, Fatima Memic, Job van der Zwan, Martin Häring, Emelie Braun, Lars E. Borm, Gioele La Manno, Simone Codeluppi, Alessandro Furlan, Kawai Lee, Nathan Skene, Kenneth D. Harris, Jens Hjerling-Leffler, Ernest Arenas, Patrik Ernfors, Ulrika Marklund, and Sten Linnarsson. Molecular Architecture of the Mouse Nervous System. Cell, 174(4):999–1014.e22, 2018. doi: 10.1016/j.cell.2018.06.021. URL 10.1016/j.cell.2018.06.021.

[2] Zizhen Yao, Cindy T. J. van Velthoven, Michael Kunst, Meng Zhang, Delissa McMillen, Changkyu Lee, Won Jung, Jeff Goldy, Aliya Abdelhak, Matthew Aitken, Katherine Baker, Pamela Baker, Eliza Barkan, Darren Bertagnolli, Ashwin Bhandiwad, Cameron Bielstein, Prajal Bishwakarma, Jazmin Campos, Daniel Carey, Tamara Casper, Anish Bhaswanth Chakka, Rushil Chakrabarty, Sakshi Chavan, Min Chen, Michael Clark, Jennie Close, Kirsten Crichton, Scott Daniel, Peter DiValentin, Tim Dolbeare, Lauren Ellingwood, Elysha Fiabane, Timothy Fliss, James Gee, James Gerstenberger, Alexandra Glandon, Jessica Gloe, Joshua Gould, James Gray, Nathan Guilford, Junitta Guzman, Daniel Hirschstein, Windy Ho, Marcus Hooper, Mike Huang, Madie Hupp, Kelly Jin, Matthew Kroll, Kanan Lathia, Arielle Leon, Su Li, Brian Long, Zach Madigan, Jessica Malloy, Jocelin Malone, Zoe Maltzer, Naomi Martin, Rachel McCue, Ryan McGinty, Nicholas Mei, Jose Melchor, Emma Meyerdierks, Tyler Mollenkopf, Skyler Moonsman, Thuc Nghi Nguyen, Sven Otto, Trangthanh Pham, Christine Rimorin, Augustin Ruiz, Raymond Sanchez, Lane Sawyer, Nadiya Shapovalova, Noah Shepard, Cliff Slaughterbeck, Josef Sulc, Michael Tieu, Amy Torkelson, Herman Tung, Nasmil Valera Cuevas, Shane Vance, Katherine Wadhwani, Katelyn Ward, Boaz Levi, Colin Farrell, Rob Young, Brian Staats, Ming-Qiang Michael Wang, Carol L. Thompson, Shoaib Mufti, Chelsea M. Pagan, Lauren Kruse, Nick Dee, Susan M. Sunkin, Luke Esposito, Michael J. Hawrylycz, Jack Waters, Lydia Ng, Kimberly Smith, Bosiljka Tasic, Xiaowei Zhuang, and Hongkui Zeng. A high-resolution transcriptomic and spatial atlas of cell types in the whole mouse brain. Nature, 624(7991):317–332, 2023. doi: 10.1038/s41586-023-06812-z. URL 10.1038/s41586-023-06812-z.

[3] Hadas Keren-Shaul, Amit Spinrad, Assaf Weiner, Orit Matcovitch-Natan, Raz Dvir-Szternfeld, Tyler K. Ulland, Eyal David, Kuti Baruch, David Lara-Astaiso, Beata Toth, Shalev Itzkovitz, Marco Colonna, Michal Schwartz, and Ido Amit. A Unique Microglia Type Associated with Restricting Development of Alzheimer’s Disease. Cell, 169(7):1276–1290.e17, 2017. doi: 10.1016/j.cell.2017.05.018. URL 10.1016/j.cell.2017.05.018.

[4] Carlo Sala Frigerio, Leen Wolfs, Nicola Fattorelli, Nicola Thrupp, Iryna Voytyuk, Inga Schmidt, Renzo Mancuso, Wei-Ting Chen, Maya E. Woodbury, Gyan Srivastava, Thomas Möller, Eloise Hudry, Sudeshna Das, Takaomi Saido, Eric Karran, Bradley Hyman, V. Hugh Perry, Mark Fiers, and Bart De Strooper. The Major Risk Factors for Alzheimer’s Disease: Age, Sex, and Genes Modulate the Microglia Response to A*β* Plaques. Cell Reports, 27(4): 1293–1306.e6, 2019. doi: 10.1016/j.celrep.2019.03.099. URL 10.1016/j.celrep.2019.03.099.

[5] Timothy R. Hammond, Connor Dufort, Lasse Dissing-Olesen, Stefanie Giera, Adam Young, Alec Wysoker, Alec J. Walker, Frederick Gergits, Michael Segel, James Nemesh, Samuel E. Marsh, Arpiar Saunders, Evan Macosko, Florent Ginhoux, Jinmiao Chen, Robin J.M. Franklin, Xianhua Piao, Steven A. McCarroll, and Beth Stevens. Single-Cell RNA Sequencing of Microglia throughout the Mouse Lifespan and in the Injured Brain Reveals Complex Cell-State Changes. Immunity, 50(1):253–271.e6, 2019. doi: 10.1016/j.immuni.2018.11.004. URL 10.1016/j.immuni.2018.11.004.

[6] Holly Oakley, Sarah L. Cole, Sreemathi Logan, Erika Maus, Pei Shao, Jeffery Craft, Angela Guillozet-Bongaarts, Masuo Ohno, John Disterhoft, Linda Van Eldik, Robert Berry, and Robert Vassar. Intraneuronal *β*-Amyloid Aggregates, Neurodegeneration, and Neuron Loss in Transgenic Mice with Five Familial Alzheimer’s Disease Mutations: Potential Factors in Amyloid Plaque Formation. The Journal of Neuroscience, 26(40):10129–10140, 2006. doi: 10.1523/JNEUROSCI.1202-06.2006. URL 10.1523/JNEUROSCI.1202-06.2006.

[7] Eric D. Sun, Rahul Nagvekar, Angela N. Pogson, and Anne Brunet. Brain aging and rejuvenation at single-cell resolution. Neuron, 113(1):82–108, 2025. doi: 10.1016/j.neuron.2024.12.007. URL 10.1016/j.neuron.2024.12.007.

[8] Tanya Barrett, Stephen E. Wilhite, Pierre Ledoux, Carlos Evangelista, Irene F. Kim, Maxim Tomashevsky, Kimberly A. Marshall, Katherine H. Phillippy, Patti M. Sherman, Michelle Holko, Andrey Yefanov, Hyeseung Lee, Naigong Zhang, Cynthia L. Robertson, Nadezhda Serova, Sean Davis, and Alexandra Soboleva. NCBI GEO: archive for functional genomics data sets—update. Nucleic Acids Research, 41(D1):D991–D995, 2013. doi: 10.1093/nar/gks1193. URL 10.1093/nar/gks1193.

[9] Anja Füllgrabe, Nancy George, Matthew Green, Parisa Nejad, Bruce Aronow, Silvie Korena Fexova, Clay Fischer, Mallory Ann Freeberg, Laura Huerta, Norman Morrison, Richard H. Scheuermann, Deanne Taylor, Nicole Vasilevsky, Laura Clarke, Nils Gehlenborg, Jim Kent, John Marioni, Sarah Teichmann, Alvis Brazma, and Irene Papatheodorou. Guidelines for reporting single-cell RNA-seq experiments. Nature Biotechnology, 38(12):1384–1386, 2020. doi: 10.1038/s41587-020-00744-z. URL 10.1038/s41587-020-00744-z.

[10] Sidhant Puntambekar, Jay R. Hesselberth, Kent A. Riemondy, and Rui Fu. Cell-level metadata are indispensable for documenting single-cell sequencing datasets. PLOS Biology, 19(5):e3001077, 2021. doi: 10.1371/journal.pbio.3001077. URL 10.1371/ journal.pbio.3001077.

[11] Jing Jiang, Cankun Wang, Ren Qi, Hongjun Fu, and Qin Ma. scREAD: A Single-Cell RNA-Seq Database for Alzheimer’s Disease. iScience, 23(11):101769, 2020. doi: 10.1016/j.isci.2020.101769. URL 10.1016/j.isci.2020.101769.

[12] Cankun Wang, Diana Acosta, Megan McNutt, Jiang Bian, Anjun Ma, Hongjun Fu, and Qin Ma. A single-cell and spatial RNA-seq database for Alzheimer’s disease (ssREAD). Nature Communications, 15(1):4710, 2024. doi: 10.1038/s41467-024-49133-z. URL 10.1038/s41467-024-49133-z.

[13] Xin-Wen Li, Ting-Ting Duan, Jin-Yu Chu, Shi-Yao Pan, Yan Zeng, and Fei-Fei Hu. SCAD-Brain: a public database of single cell RNA-seq data in human and mouse brains with Alzheimer’s disease. Frontiers in Aging Neuroscience, 15:1157792, 2023. doi: 10.3389/fnagi.2023.1157792. URL 10.3389/fnagi.2023.1157792.

[14] CZI Cell Science Program, Shibla Abdulla, Brian Aevermann, Pedro Assis, Seve Badajoz, Sidney M Bell, Emanuele Bezzi, Batuhan Cakir, Jim Chaffer, Signe Chambers, J Michael Cherry, Tiffany Chi, Jennifer Chien, Leah Dorman, Pablo Garcia-Nieto, Nayib Gloria, Mim Hastie, Daniel Hegeman, Jason Hilton, Timmy Huang, Amanda Infeld, Ana-Maria Istrate, Ivana Jelic, Kuni Katsuya, Yang Joon Kim, Karen Liang, Mike Lin, Maximilian Lombardo, Bailey Marshall, Bruce Martin, Fran McDade, Colin Megill, Nikhil Patel, Alexander Predeus, Brian Raymor, Behnam Robatmili, Dave Rogers, Erica Rutherford, Dana Sadgat, Andrew Shin, Corinn Small, Trent Smith, Prathap Sridharan, Alexander Tarashansky, Norbert Tavares, Harley Thomas, Andrew Tolopko, Meghan Urisko, Joyce Yan, Garabet Yeretssian, Jennifer Zamanian, Arathi Mani, Jonah Cool, and Ambrose Carr. CZ CELLxGENE Discover: a single-cell data platform for scalable exploration, analysis and modeling of aggregated data. Nucleic Acids Research, 53(D1):D886–D900, 2025. doi: 10.1093/nar/gkae1142. URL 10.1093/nar/gkae1142.

[15] Nicholas D. Youngblut, Christopher Carpenter, Arshia Nayebnazar, Abhinav Adduri, Rohan Shah, Chiara Ricci-Tam, Jaanak Prashar, Rajesh Ilango, Noam Teyssier, Silvana Konermann, Patrick D. Hsu, Alexander Dobin, Dave P. Burke, Hani Goodarzi, and Yusuf H. Roohani. scBaseCount: An AI agent-curated, standardized, auto-updated single-cell data repository. Cell, 189(19):5932–5944.e6, 2026. doi: 10.1016/j.cell.2026.08.025. URL 10.1016/j.cell.2026.08.025.

[16] Arc Institute. scBaseCount: metadata documentation and disease-annotation confidence audit. Project documentation, 2026. URL https://github.com/ArcInstitute/arc-virtual-cell-atlas/blob/main/scBaseCount/README.md. Disease annotations and per-SRX confidence audit; accessed 24 October 2026.

[17] Nicola Thrupp, Carlo Sala Frigerio, Leen Wolfs, Nathan G. Skene, Nicola Fattorelli, Suresh Poovathingal, Yannick Fourne, Paul M. Matthews, Tom Theys, Renzo Mancuso, Bart de Strooper, and Mark Fiers. Single-Nucleus RNA-Seq Is Not Suitable for Detection of Microglial Activation Genes in Humans. Cell Reports, 32(13):108189, 2020. doi: 10.1016/j.celrep.2020.108189. URL 10.1016/j.celrep.2020.108189.

[18] Samuel E. Marsh, Alec J. Walker, Tushar Kamath, Lasse Dissing-Olesen, Timothy R. Hammond, T. Yvanka de Soysa, Adam M. H. Young, Sarah Murphy, Abdulraouf Abdulraouf, Naeem Nadaf, Connor Dufort, Alicia C. Walker, Liliana E. Lucca, Velina Kozareva, Charles Vanderburg, Soyon Hong, Harry Bulstrode, Peter J. Hutchinson, Daniel J. Gaffney, David A. Hafler, Robin J. M. Franklin, Evan Z. Macosko, and Beth Stevens. Dissection of artifactual and confounding glial signatures by single-cell sequencing of mouse and human brain. Nature Neuroscience, 25(3):306–316, 2022. doi: 10.1038/s41593-022-01022-8. URL 10.1038/s41593-022-01022-8.

[19] Romain Lopez, Jeffrey Regier, Michael B. Cole, Michael I. Jordan, and Nir Yosef. Deep generative modeling for single-cell transcriptomics. Nature Methods, 15(12):1053–1058, 2018. doi: 10.1038/s41592-018-0229-2. URL 10.1038/s41592-018-0229-2.

[20] Malte D. Luecken, M. Büttner, K. Chaichoompu, A. Danese, M. Interlandi, M. F. Mueller, D. C. Strobl, L. Zappia, M. Dugas, M. Colomé-Tatché, and Fabian J. Theis. Benchmarking atlas-level data integration in single-cell genomics. Nature Methods, 19(1):41–50, 2022. doi: 10.1038/s41592-021-01336-8. URL 10.1038/s41592-021-01336-8.

[21] Sean Davis and Paul S. Meltzer. GEOquery: a bridge between the Gene Expression Omnibus (GEO) and BioConductor. Bioinformatics, 23(14):1846–1847, 2007. doi: 10.1093/bioinformatics/btm254. URL 10.1093/bioinformatics/btm254.

[22] Christopher J Mungall, Carlo Torniai, Georgios V Gkoutos, Suzanna E Lewis, and Melissa A Haendel. Uberon, an integrative multi-species anatomy ontology. Genome Biology, 13(1): R5, 2012. doi: 10.1186/gb-2012-13-1-r5. URL 10.1186/gb-2012-13-1-r5.

[23] Aaron T. L. Lun, Samantha Riesenfeld, Tallulah Andrews, The Phuong Dao, Tomas Gomes, participants in the 1st Human Cell Atlas Jamboree, and John C. Marioni. EmptyDrops: distinguishing cells from empty droplets in droplet-based single-cell RNA sequencing data. Genome Biology, 20(1):63, 2019. doi: 10.1186/s13059-019-1662-y. URL 10.1186/s13059-019-1662-y.

[24] Ayshwarya Subramanian, Mikhail Alperovich, Yiming Yang, and Bo Li. Biology-inspired data-driven quality control for scientific discovery in single-cell transcriptomics. Genome Biology, 23(1):267, 2022. doi: 10.1186/s13059-022-02820-w. URL 10.1186/s13059-022-02820-w.

[25] F. Alexander Wolf, Philipp Angerer, and Fabian J. Theis. SCANPY: large-scale single-cell gene expression data analysis. Genome Biology, 19(1):15, 2018. doi: 10.1186/s13059-017-1382-0. URL 10.1186/s13059-017-1382-0.

[26] Samuel L. Wolock, Romain Lopez, and Allon M. Klein. Scrublet: Computational Identification of Cell Doublets in Single-Cell Transcriptomic Data. Cell Systems, 8(4):281–291.e9, 2019. doi: 10.1016/j.cels.2018.11.005. URL 10.1016/j.cels.2018.11.005.

[27] Emma Busarello, Giulia Biancon, Ilaria Cimignolo, Fabio Lauria, Zuhairia Ibnat, Christian Ramirez, Gabriele Tomè, Marianna Ciuffreda, Giorgia Bucciarelli, Alessandro Pilli, Stefano Maria Marino, Vittorio Bontempi, Federica Ress, Kristin R. Aass, Jennifer VanOudenhove, Luca Tiberi, Maria Caterina Mione, Therese Standal, Paolo Macchi, Gabriella Viero, Stephanie Halene, and Toma Tebaldi. Cell Marker Accordion: interpretable single-cell and spatial omics annotation in health and disease. Nature Communications, 16(1):5399, 2025. doi: 10.1038/s41467-025-60900-4. URL 10.1038/s41467-025-60900-4.

[28] Alexander D. Diehl, Terrence F. Meehan, Yvonne M. Bradford, Matthew H. Brush, Wasila M. Dahdul, David S. Dougall, Yongqun He, David Osumi-Sutherland, Alan Ruttenberg, Sirarat Sarntivijai, Ceri E. Van Slyke, Nicole A. Vasilevsky, Melissa A. Haendel, Judith A. Blake, and Christopher J. Mungall. The Cell Ontology 2016: enhanced content, modularization, and ontology interoperability. Journal of Biomedical Semantics, 7(1):44, 2016. doi: 10.1186/s13326-016-0088-7. URL 10.1186/s13326-016-0088-7.

[29] Tobias Goldmann, Peter Wieghofer, Marta Joana Costa Jordão, Fabiola Prutek, Nora Hagemeyer, Kathrin Frenzel, Lukas Amann, Ori Staszewski, Katrin Kierdorf, Martin Krueger, Giuseppe Locatelli, Hannah Hochgerner, Robert Zeiser, Slava Epelman, Frederic Geissmann, Josef Priller, Fabio M V Rossi, Ingo Bechmann, Martin Kerschensteiner, Sten Linnarsson, Steffen Jung, and Marco Prinz. Origin, fate and dynamics of macrophages at central nervous system interfaces. Nature Immunology, 17(7):797–805, 2016. doi: 10.1038/ni.3423. URL 10.1038/ni.3423.

[30] Takahiro Masuda, Lukas Amann, Gianni Monaco, Roman Sankowski, Ori Staszewski, Martin Krueger, Francesca Del Gaudio, Liqun He, Neil Paterson, Elisa Nent, Francisco Fernández-Klett, Ayato Yamasaki, Maximilian Frosch, Maximilian Fliegauf, Lance Fredrick Pahutan Bosch, Hatice Ulupinar, Nora Hagemeyer, Dietmar Schreiner, Cayce Dorrier, Makoto Tsuda, Claudia Grothe, Anne Joutel, Richard Daneman, Christer Betsholtz, Urban Lendahl, Klaus-Peter Knobeloch, Tim Lämmermann, Josef Priller, Katrin Kierdorf, and Marco Prinz. Specification of CNS macrophage subsets occurs postnatally in defined niches. Nature, 604 (7907):740–748, 2022. doi: 10.1038/s41586-022-04596-2. URL 10.1038/s41586-022-04596-2.

[31] John N. Hutchinson, Alexander W. Ensminger, Christine M. Clemson, Christopher R. Lynch, Jeanne B. Lawrence, and Andrew Chess. A screen for nuclear transcripts identifies two linked noncoding RNAs associated with SC35 splicing domains. BMC Genomics, 8:39, 2007. doi: 10.1186/1471-2164-8-39. URL 10.1186/1471-2164-8-39.

[32] Damian Smedley, Syed Haider, Benoit Ballester, Richard Holland, Darin London, Gudmundur Thorisson, and Arek Kasprzyk. BioMart – biological queries made easy. BMC Genomics, 10(1):22, 2009. doi: 10.1186/1471-2164-10-22. URL 10.1186/14 71-2164-10-22.

[33] Tim Stuart, Andrew Butler, Paul Hoffman, Christoph Hafemeister, Efthymia Papalexi, William M. Mauck, III, Yuhan Hao, Marlon Stoeckius, Peter Smibert, and Rahul Satija. Comprehensive Integration of Single-Cell Data. Cell, 177(7):1888–1902.e21, 2019. doi: 10.1016/j.cell.2019.05.031. URL 10.1016/j.cell.2019.05.031.

[34] Leland McInnes, John Healy, and James Melville. UMAP: Uniform Manifold Approximation and Projection for Dimension Reduction. arXiv, 2018. doi: 10.48550/arXiv.1802.03426. URL https://arxiv.org/abs/1802.03426. Preprint.

[35] Ilya Korsunsky, Nghia Millard, Jean Fan, Kamil Slowikowski, Fan Zhang, Kevin Wei, Yuriy Baglaenko, Michael Brenner, Po-ru Loh, and Soumya Raychaudhuri. Fast, sensitive and accurate integration of single-cell data with Harmony. Nature Methods, 16(12):1289–1296, 2019. doi: 10.1038/s41592-019-0619-0. URL 10.1038/s41592-019-0619-0.

[36] Gilad Sahar Green, Masashi Fujita, Hyun-Sik Yang, Mariko Taga, Anael Cain, Cristin McCabe, Natacha Comandante-Lou, Charles C. White, Anna K. Schmidtner, Lu Zeng, Alina Sigalov, Yangling Wang, Aviv Regev, Hans-Ulrich Klein, Vilas Menon, David A. Bennett, Naomi Habib, and Philip L. De Jager. Cellular communities reveal trajectories of brain ageing and Alzheimer’s disease. Nature, 633(8030):634–645, 2024. doi: 10.1038/s41586-024-07871-6. URL 10.1038/s41586-024-07871-6.

[37] Emma Gerrits, Nieske Brouwer, Susanne M. Kooistra, Maya E. Woodbury, Yannick Vermeiren, Mirjam Lambourne, Jan Mulder, Markus Kummer, Thomas Möller, Knut Biber, Wilfred F. A. den Dunnen, Peter P. De Deyn, Bart J. L. Eggen, and Erik W. G. M. Boddeke. Distinct amyloid-*β* and tau-associated microglia profiles in Alzheimer’s disease. Acta Neuropathologica, 141(5):681–696, 2021. doi: 10.1007/s00401-021-02263-w. URL 10.1007/s00401-021-02263-w.

[38] Shunta Sakaguchi, Masato Tsutsumi, Kentaro Nishi, and Honda Naoki. Disentangling batch effects and biological signals across conditions in single-cell transcriptome. Patterns, 7(10): 101638, 2026. doi: 10.1016/j.patter.2026.101638. URL 10.1016/j.patter.2026.101638.

[39] Emil Uffelmann, Douglas P. Wightman, et al. Alzheimer’s disease genetics converges on microglial and neuronal dysregulation. Nature, 2026. doi: 10.1038/s41586-026-11206-y. URL 10.1038/s41586-026-11206-y.

[40] LRAPA datasets contributors. AD-associated genes reference list. https://github.com/yalanyang/LRAPA_datasets/blob/main/reference/genelist/AD_associated_genes.txt. GitHub repository. Accessed: 2026-11-03.

[41] ADSP Gene Verification Committee. Ad/adrd top genes repository. https://topgenes.niagads.org/. Accessed: 2026-11-03.

[42] Susanne Krasemann, Charlotte Madore, Ron Cialic, Caroline Baufeld, Narghes Calcagno, Rachid El Fatimy, Lien Beckers, Elaine O’Loughlin, Yang Xu, Zain Fanek, David J. Greco, Scott T. Smith, George Tweet, Zachary Humulock, Tobias Zrzavy, Patricia Conde-Sanroman, Mar Gacias, Zhiping Weng, Hao Chen, Emily Tjon, Fargol Mazaheri, Kristin Hartmann, Asaf Madi, Jason D. Ulrich, Markus Glatzel, Anna Worthmann, Joerg Heeren, Bogdan Budnik, Cynthia Lemere, Tsuneya Ikezu, Frank L. Heppner, Vladimir Litvak, David M. Holtzman, Hans Lassmann, Howard L. Weiner, Jordi Ochando, Christian Haass, and Oleg Butovsky. The TREM2-APOE Pathway Drives the Transcriptional Phenotype of Dysfunctional Microglia in Neurodegenerative Diseases. Immunity, 47(3):566–581.e9, 2017. doi: 10.1016/j.immuni.2017.08.008. URL 10.1016/j.immuni.2017.08.008.

