## Supplementary Methods, Tables, and Results for "Harmonized sample metadata and processed cortical and hippocampal transcriptomes from wild-type and Alzheimer’s disease model mice"

##### Data structure and analysis units

The GSM-level metadata schema describes 2,005 biologically eligible records, including 142 without usable or reliably assignable expression data (Supplementary Table S4). The 1,863 count-available records yielded 2,006 processing libraries. Local GSM000 identifiers and numeric library suffixes preserve the relationship between count-matrix groups, processing libraries and their source studies. GEO links, available publication links and curatorial notes support record-level inspection, but do not constitute complete field-level provenance. Library-specific information was used when available, including sex assignments for the split GSE185862 libraries.

##### LLM-assisted screening: input preparation, decoding and parsing

Each GSM was represented by three study-level and six sample-level fields (Supplementary Table S5). Whitespace was collapsed within each field, and the GSE/GSM identifier fields were omitted from the classifier input. Accession identifiers or URLs embedded in free-text metadata were not systematically stripped. The nominal budget was 16,000 characters for the combined field values. If that budget was exceeded, text was trimmed from the end of fields in the following order, from lowest to highest priority: growth protocol, overall summary, study title, overall design, treatment protocol, extraction protocol, sample source, sample title, and sample characteristics. Trimming did not reduce any field below 400 characters when at least 400 characters were originally available. A truncation marker was appended to shortened fields. Field labels and truncation markers were added after the budget was calculated, so the final assembled string could slightly exceed the nominal character budget.

The metadata were inserted at the `\{\{METADATA\}\}` placeholder in prompts, and the model-specific chat template was applied.

Gemma was executed in offline vLLM mode with prefix caching, temperature 0, seed 42, a maximum of 256 generated tokens, and a maximum model length of 16,384 tokens. The batch cap was 32. Prompt optimization and the final test reused that frozen configuration. Inputs whose tokenized length plus the generation budget exceeded the context window were not submitted and were flagged. The model was instructed to return one JSON object with boolean keys `assay_10x_sc_or_sn`, `target_anatomy` and `wt_or_ad_sample`, together with a binary `label`, `confidence` and `failure_reasons`. Responses were parsed as JSON, allowing removal of Markdown code fences or extraction of the outermost braces. For unparseable responses, a second inference call appended the instruction "Your answer must be ONLY one valid JSON object with the requested keys. No other text." Unparseable responses counted as misclassifications in the benchmark. During cohort-2 screening, these were treated as candidates for manual inspection. No parse errors were reported in the held-out test or cohort-2 screening.

#### LLM-assisted screening: implementation of prompt optimization

The reference set comprised 3,402 manually annotated cohort-1 GSMs from 702 GSEs (1,701 positive and 1,701 negative). It was partitioned by GSE, so that samples from one study never appeared in different sets, into training (2,050 GSMs, 302 GSEs), validation (676 GSMs, 225 GSEs) and held-out test (676 GSMs, 175 GSEs) sets.

Generation 0 consisted of a zero-shot prompt describing three necessary criteria for GSM eligibility: the assay, the anatomy, and the genotype or experimental condition. The prompt also established sample-level precedence over study-level metadata and required the structured JSON output. Generation 0 was evaluated on the complete training and validation partitions before any revisions.

For each revision round $g = 1, \ldots, 15$, incorrect training predictions from generation $g-1$ were identified, including failures to parse the JSON output. To construct Claude's instruction-revision request, up to eight false negatives and eight false positives were sampled from the training errors, with preference for one error per GSE before sampling additional errors from the same studies (sampling seed $42 + 10{,}000 + g$). Claude received the preceding instruction, training-set accuracy, precision, recall, $F_1$, confusion counts and parse-error count, criterion-specific summaries of errors using Gemma's own Boolean responses, summaries of the five most recent generations, and the selected misclassified training records with their true labels and Gemma predictions. The fixed Claude instruction asked for generalized changes explaining the errors, preservation of working rules, no sample-specific identifiers, and an unchanged JSON schema. Claude returned a short change summary and a full revised instruction in tagged blocks, which were programmatically validated. Invalid outputs were returned to Claude for revision, up to three optimizer attempts.

A separate few-shot synthesis request was assembled from up to five false-negative and five false-positive records in the preceding generation, selected randomly with seed $42 + g$. The example set was reconstructed anew for every generation rather than accumulated. Claude was asked to create one synthetic analogue per selected training record, maintaining its label, the distribution of evidence between study-level and sample-level fields, and the ambiguity and difficulty of the source record, while changing incidental details and excluding accession identifiers, URLs and author names. Each synthetic record was checked automatically for GSE/GSM identifiers and URLs. For copying detection, the set of distinct word 8-grams in the synthetic record was compared with the union of 8-grams from training records belonging to its source GSE. A record was rejected if more than 15% of its 8-grams appeared in that union. Rejected synthesis results were sent back to Claude for regeneration (up to three optimizer attempts). The optimizer also verified that each synthetic label matched its training source and that the source was an error from the preceding generation. This check verified textual overlap and label inheritance, but did not independently verify biological consistency of the generated records.

The resulting full prompt consisted of the revised instruction, a few-shot block containing each synthetic record with its assigned label, and the metadata for the GSM to be classified. Claude was used without tools and was asked to act as a prompt engineer and return only the requested tagged format. Claude Opus 5.5 (Anthropic) was the prompt-revision model. The exact resolved model identifier was not recorded. Gemma's inference configuration was fixed throughout optimization, and the full training and validation splits were evaluated at each generation. Only training data were provided to Claude, that is, validation data were used exclusively to select among completed prompts, and the test set was not loaded during optimization.

Prompts were compared using positive-class (eligible GSM) precision, recall and $F_1$, defined as

$$\text{Precision} = \frac{TP}{TP + FP}, \qquad \text{Recall} = \frac{TP}{TP + FN}, \qquad F_1 = \frac{2TP}{2TP + FP + FN}, \tag{S1}$$

where $TP$, $FP$ and $FN$ denote the numbers of true positives, false positives and false negatives among the evaluated GSMs, and unparseable responses were counted as misclassifications. The final candidate was selected by validation $F_1$ and parsimony: among prompts within 0.005 of the maximum validation $F_1$, preference was given to the fewest few-shot examples, then the shortest prompt (in characters), and finally the earliest generation.

The maximum validation $F_1$ was 0.9851 (generation 1). Generations 1, 2, 3, 5 and 8 fell within 0.005 of this value. The predefined selection rule first minimized the number of few-shot examples within this near-optimal set, with prompt character count and then generation index as tie-breakers. Generation 3 was selected, with five synthetic positive examples and no negative examples, whose full prompt comprised 15,265 characters. Its training-set precision, recall and $F_1$ were 0.9941, 0.9863 and 0.9902, and the corresponding validation values were 0.9880, 0.9734 and 0.9806. On the independent held-out test ($n = 676$, 340 positive and 336 negative samples), the frozen prompt yielded 326 true positives, 8 false positives, 328 true negatives and 14 false negatives (precision 0.9760, recall 0.9588, $F_1$ 0.9674). No additional prediction-threshold or rescue-rule optimization was applied in this evaluation. Generation-wise performance is shown in Supplementary Fig. S2a.

Benchmark samples were approximately class-balanced, so their precision and $F_1$ are not estimates of performance at the lower prevalence of relevant GSMs expected in GEO. The reference labels were frozen from an earlier curation snapshot. Two reference-positive records (GSM4658252 and GSM6252678) are absent from the final resource, while two reference-negative training records (GSM8941782 and GSM8941789 from GSE295077) are included in the final resource. The latter were retained because vascular enrichment alone was not an exclusion criterion when the species, assay, tissue and model criteria were met. Their historical training labels were preserved for the frozen benchmark. The held-out benchmark was not retrospectively relabeled, and its reported performance describes the frozen reference task. After regrouping individual-cell deposits, cohort 2 comprised 66,165 records: 478 positive and 65,687 negative. The 117 GSEs containing at least one positive prediction comprised 2,883 unique records, including all 2,405 individually negative predictions reviewed in those studies. The 307 GSEs without a positive prediction were not manually reviewed at this stage. These counts match the main selection flow. The screening inference output uses the original-accession granularity and contains 78,941 rows, all with distinct original GSM accessions, including 13,254 positive predictions and 15,659 GSMs in the selected studies. The four individual-cell series GSE93421, GSE95315, GSE99888 and GSE104323 contributed 12,790 original accessions that were replaced by 14 local grouping identifiers, reducing the reported sample counts by 12,776. Thus, the larger inference totals reflect the original record granularity. They are not additional unique samples in the regrouped collection.

#### Preliminary GPT-4o metadata extraction for cohort 1

Before the Gemma-based screening, GPT-4o (OpenAI, `gpt-4o`, temperature 0) was accessed directly using the OpenAI Chat Completions API to pre-fill annotation templates. Six separate extraction requests obtained brain region, sex, age, strain, sequencing strategy / sc-versus-sn modality / platform, and genotype / treatment. Region, sex, age and strain requests used the GSM characteristics and title, whereas the sequencing and genotype/treatment requests used additional source and protocol fields. A seventh GPT-4o request inferred an experimental-model label from the previously extracted strain, genotype and treatment rather than the original GSM record. The returned values were preliminary and the final inclusion decisions and retained annotations were established by manual review. Some studies had empty preliminary extraction output files and were therefore not included among the earlier successfully processed records. The reasons for the missing outputs could not be established. Cohort membership was not randomized and might be associated with factors affecting extraction success. Prompt S5 reproduces the literal questions and message-role wrappers.

**Model and prompt provenance.** Gemma 4 31B-it was run with offline vLLM (maximum model length 16,384, maximum generation length 256, temperature 0, seed 42, batch cap 32). The numerical dtype and quantization were not explicitly set. Claude Opus 5.5 was the prompt-revision model. Its exact resolved identifier was not recorded. The generation-0 seed and generation-3 selected prompts contain 2,379 and 15,265 characters, respectively. The held-out split contained 676 GSMs from 175 GSEs (340 positive and 336 negative reference labels), and no GSE occurred in more than one split. Class balancing and study prioritization limit transfer of benchmark precision and recall to the complete GEO population.

#### Readers and mappings for deposited count matrices

Dedicated readers imported deposited 10x, HDF5, delimited, R-object and AnnData count matrices, and study-specific barcode and metadata mappings assigned cells to samples. The standard barcode–feature–matrix reader accepted compressed or uncompressed 10x Matrix Market files with either `genes.tsv` or `features.tsv`. The 10x HDF5 reader retained Gene Expression features and preferred a raw HDF5 file when raw and filtered versions were both present. Archive readers extracted compatible Matrix Market or HDF5 structures. When several matrix directories were present in an archive, the archive helper preferred a filtered subdirectory, so raw-file preference was not universal across readers. Delimited-table readers accepted explicit orientation or inferred gene and cell axes, removed nonnumeric annotation columns, summed repeated features and combined cluster-partitioned tables when necessary. R-object readers extracted RNA count slots or layers from Seurat objects, count assays from SummarizedExperiment-derived objects, or counts from supported matrix and data-frame representations. For AnnData, an explicitly requested layer took precedence. Otherwise the reader attempted `/layers/counts`, `/raw/X` and `/X` in that order. Sparse CSR, sparse CSC and dense encodings were supported, and count validity was checked downstream rather than inferred from the AnnData field name.

Series-level matrices were assigned to GSMs using explicit maps from barcode prefixes or suffixes, sample metadata, and GEO titles or characteristics. GSE185862 used a custom HDF5 reader for `/data/counts`, `/data/gene` and `/data/samples`, processed in blocks of 2,000 cells. GSE104323 was regrouped by the reported E16.5, P0, P120, P132, P18, P19, P23 and P5 age labels into local records GSM000007–GSM000014. GSE95315 used the deposited age-group divisions to define the local pooled-sample records GSM000003 and GSM000004. The GSE314059 reader handled the deposited long-read gene read-count tables for three mice, corresponding to GSM9380799–GSM9380801. These source counts are read counts and are not relabeled as UMIs. Normalized-only matrices from GSE138381 and GSE205936 were excluded from the count-available cohort. Split-library identifiers preserve their parent record with a numeric suffix.

Features were harmonized to mouse gene symbols. Only Gene Expression features were retained when feature types were available. Ensembl-only identifiers were converted using the `mmusculus_gene_ensembl` BioMart mapping after removal of version suffixes [1], retrieved from the Ensembl BioMart service (US East mirror) on 2026-10-14. Unmapped identifiers, spike-in controls and unnamed features were discarded, and features with the same symbol were summed. Reference symbols were matched exactly or, where unambiguous, without regard to case. Matrices containing non-integer or negative values were rejected.

#### Cell calling and exceptional cases

The empty-droplet screen counted all barcodes with total counts below 100, including zero-count barcodes, and triggered when this number exceeded 1,000. For libraries meeting this criterion, only barcodes with positive totals were passed to STAR's `soloCellFiltering` mode. The `EmptyDrops_CR` parameters were an expected cell number of 3,000, a maximum percentile of 0.99, a maximum-to-minimum ratio of 10, ambient-profile barcode indices of 45,000–90,000, a minimum count of 500, a minimum fraction of median counts of 0.01, at most 20,000 candidate barcodes, a false-discovery rate of 0.01 and 10,000 simulations. The retained set combined cells called by the ordinal-magnitude stage and by the subsequent EmptyDrops test. A result with zero EmptyDrops candidates and no additional EmptyDrops calls was accepted. The procedure required sufficient nonzero-count barcodes to estimate the ambient profile.

Libraries that did not meet the empty-droplet-screen criterion retained every positive-count barcode. Thus the cell-calling denominator combines barcodes called from unfiltered inputs with cells previously filtered by the original study. The original-barcode total is the sum of nonzero-count input barcodes after feature harmonization, rather than an estimate derived from the number of features or matrix dimensions.

#### Clustering, ddqc and doublet detection

For preliminary clustering and reclustering of singlets, zero-count genes were removed, counts were normalized to 10,000 per cell and log1p-transformed, and 2,000 highly variable genes were selected using Scanpy's default Seurat flavor. Expression values were scaled with a maximum value of 10, up to 50 principal components were computed with ARPACK, and a ten-neighbor graph used up to the first 30 components. The number of components was bounded by the input dimensions in the implementation. Leiden clustering used the default resolution of 1 and a deterministic library-specific random seed.

The `ddqc_scanpy` configuration used the MAD method with threshold 2, lower bounds for gene and total-count metrics, upper bounds for mitochondrial and ribosomal percentages, and no additional fixed thresholds or mitochondrial-percentage cap. Mitochondrial and ribosomal genes were taken from GENCODE vM38 (GRCm39)-derived mouse gene lists. When no mitochondrial or ribosomal reference gene was available in a library, the corresponding metric was recorded as missing and was inactive for ddqc.

Scrublet used an expected doublet rate of 0.05, a simulated-to-observed ratio of 2 and an automatically selected threshold. The initial principal-component count was 30. When the input dimensions were insufficient, the implementation retried with a smaller count. If no threshold was returned, all cells were retained as singlets. The within-library QC comparisons use removed versus retained cells for ddqc and predicted doublets versus singlets for Scrublet. For a metric $m$ describing total counts or detected genes, the displayed contrast was $\log_2\{\max[\mathrm{median}(m_{\mathrm{removed}}), 1]/\max[\mathrm{median}(m_{\mathrm{retained}}), 1]\}$. The lower bound of one did not alter any of the usable gene-count or total-count comparisons. For mitochondrial and ribosomal percentages it was $\mathrm{median}(m_{\mathrm{removed}}) - \mathrm{median}(m_{\mathrm{retained}})$ in percentage points. Libraries without a usable comparison were omitted for that metric only.

#### Exact Cell Marker Accordion weighting and assignment

The mouse healthy-tissue CMA database was restricted to the sample's exact tissue terms and to markers present among the library's gene symbols. Rows were deduplicated by cell type, Cell Ontology identifier, marker, marker direction and source resource across the selected tissues. Only positive markers were present in the selected tissue subset. The implementation explicitly rejects negative markers.

For a gene $g$ and candidate cell type $t$, evidence consistency $E_{gt}$ was the number of distinct supporting source resources. Specificity was based on $K_g$, the number of candidate cell types supported by that gene, calculated before the minimum-marker filter. If there were $L$ candidate types, the initial reverse rescaling was $r_g = L + 1 - K_g$. This value was rescaled over its observed range to the range of evidence consistencies,

$$r'_g = E_{\min} + \frac{r_g - r_{\min}}{r_{\max} - r_{\min}}(E_{\max} - E_{\min}), \tag{S2}$$

where $E_{\min}$ and $E_{\max}$ are taken over the gene–type marker table for that library. For a zero input range, rescaling returned the midpoint of the target range. The final weight was

$$w_{gt} = (1 + \log_{10} E_{gt})(1 + \log_{10} r'_g). \tag{S3}$$

Cell types with fewer than five available markers were then removed, and at most 30 markers per type were retained in decreasing weight order. Stable ties were resolved from the marker and cell-type ordering. These weights depend on the available feature set as well as the tissue-specific marker database.

For cell $c$, observed counts were normalized over all retained features and transformed as

$$x_{cg} = \log\left(1 + 10^4 \frac{n_{cg}}{\sum_h n_{ch}}\right). \tag{S4}$$

Each marker was centered and scaled across the library's singlet cells using the sample standard deviation, with a zero-standard-deviation marker assigned zero. Only the upper tail was clipped, so $z_{cg} = \min[(x_{cg} - \bar{x}_g)/s_g, 10]$. For retained marker set $M_t$, the cell score was

$$S_{ct} = \frac{\sum_{g \in M_t} w_{gt} z_{cg}}{\sqrt{\sum_{g \in M_t} w_{gt}}}. \tag{S5}$$

Each cell's preliminary best type was $b_c = \arg\max_t S_{ct}$. For each cluster $k$ and type $t$, $Q_{kt}$ was the 75th percentile of $S_{ct}$ among all cells in $k$, using the standard linearly interpolated quantile. A cell's preliminary label was changed to Unknown if $Q_{k,b_c} \leq 0$. The cluster candidate was the type with the highest $Q_{kt}$ among types selected as $b_c$ by at least one cluster cell. The cluster was assigned this candidate only if its score was positive and at least 10% of cells retained that type as their preliminary label. Otherwise it was labeled Unknown. Libraries with no eligible marker set or at most 50 singlets were also labeled Unknown.

#### Internal consistency displays for cell-type annotation

For each cluster and target major class, the CMA display first selected the maximum 75th-percentile score over candidate cell types in that class. For each library and identified major class, it then took the cell-count-weighted median of these scores over the assigned clusters. This is a library-level summary of cluster scores, rather than a direct per-cell mean.

The marker display selected five positive CMA markers per major class by the fraction of libraries voting for each marker among their clusters' top-contributing markers. Eligibility required a gene to be present in at least 50% of libraries containing at least ten cells of that class. Within each library and class with at least ten cells, the plotted value was the mean cell-level log1p-normalized expression, including zero expression. A gene absent from the library's features was treated as missing rather than as a measured zero. Because the marker selection used the same marker resource that generated the cell labels, this analysis evaluates internal consistency and does not independently measure annotation accuracy.

#### Transcriptomic indicators of sc/sn protocol

For each library, the mean cell-level log1p expression of the nuclear-enriched long noncoding RNAs *Malat1* and *Neat1* was computed after normalization to 10,000 total counts [2], together with the mean mitochondrial count percentage as a complementary protocol-associated indicator. Mitochondrial symbols were recognized by the prefixes *mt-* or *mt.*, without regard to case. Missing metrics were excluded separately for each comparison: 36 libraries lacked *Malat1*, 44 lacked *Neat1*, and 23 lacked the mitochondrial metric, affecting 68 distinct libraries in total. Protocol and sex indicators were compared between annotation groups using two-sided Welch's $t$-tests on library-level summaries. These comparisons are descriptive because multiple libraries can derive from one study or animal.

#### Sex-indicator definitions and reference fits

The reference table contains 2,651 X-linked and 1,586 Y-linked Ensembl records, corresponding to 2,625 and 1,567 unique nonmissing gene symbols, respectively. Reference genes were represented as sets of symbols, matched exactly or without regard to case, and repeated feature mappings were summed. The detected-gene indicator in Fig. 2c,d is

$$R_s^{\text{detected}} = \frac{N_s^{-1} \sum_{c \in s} \sum_{g \in Y} \mathbb{I}(n_{cg} > 0)}{N_s^{-1} \sum_{c \in s} \sum_{g \in X} \mathbb{I}(n_{cg} > 0)}. \tag{S6}$$

For each cell, genes with positive counts were counted separately for the X and Y reference sets, and the ratio of the library-level mean numbers of detected Y and X genes was the detected-gene indicator, and one library lacked a usable ratio. Female- and Male-labeled libraries respectively above and below the mean ratio of Mix libraries were flagged for source review. These flags identify disagreements for inspection and are not a classifier of metadata errors. The source-review threshold was the mean $R_s^{\text{detected}}$ among Mix libraries, equal to 0.008769985. It flagged 38 Female-labeled libraries above the threshold and 19 Male-labeled libraries below it.

The expression-based indicator for Fig. 2e and the Kanade sex covariate instead uses

$$R_s^{\text{expression}} = \frac{N_s^{-1} \sum_{c \in s} \sum_{g \in Y} x_{cg}}{N_s^{-1} \sum_{c \in s} \sum_{g \in X} x_{cg}}, \qquad x_{cg} = \log\left(1 + 10^4 \frac{n_{cg}}{\sum_h n_{ch}}\right). \tag{S7}$$

Thus the two Y/X indicators must not be interchanged. For $u_s = \log_{10}(R_s^{\text{expression}} + 10^{-5})$, reference means $\mu_M$ and $\mu_F$ were estimated from labeled Male and Female libraries separately within sc and sn. The shared variance and prior were

$$\sigma^2 = \frac{\sum_{s \in M}(u_s - \mu_M)^2 + \sum_{s \in F}(u_s - \mu_F)^2}{n_M + n_F - 2}, \qquad \pi_M = \frac{n_M}{n_M + n_F}. \tag{S8}$$

The posterior indicator was

$$P(M \mid u) = \frac{\pi_M \exp[-(u - \mu_M)^2/(2\sigma^2)]}{\pi_M \exp[-(u - \mu_M)^2/(2\sigma^2)] + (1 - \pi_M) \exp[-(u - \mu_F)^2/(2\sigma^2)]}. \tag{S9}$$

The reference curves for Fig. 2e were fitted to all eligible QC libraries with Male or Female labels. The sc fit uses 296 Male and 179 Female libraries, and the sn fit uses 401 Male and 349 Female libraries. The histogram contains 317 sc and 352 sn Unknown-sex libraries, with 30 bins for the matched expression-based histogram. One Unknown-sex library, GSM8642381, has no usable Y/X ratio and is omitted. Using the expression-based ratio consistently, 261 of 317 sc libraries and 235 of 352 sn libraries have a posterior indicator above 0.5. The remaining 56 sc and 117 sn libraries fall below 0.5. These expression-based assignments do not establish animal-sex proportions. For Kanade, the same indicator and reference model were used, but its reference parameters were fitted within the Kanade-eligible cohort, so its fitted curves need not be numerically identical.

#### scVI training and subsampling

The global analysis sampled at most 100 cells per library, dividing this allocation among the cell classes present and redistributing shortfalls when a class contained fewer cells than its allocation. Class-specific analyses sampled at most 500 cells of the target class per library. Sampling was deterministic given the global seed and library identifier, with the class identifier included for class-specific analyses. HVG selection ran Seurat v3 [3] separately within each GSE on raw counts, retained the median available within-GSE rank, and broke ties using the number of GSEs in which the gene was ranked. GSEs were skipped with fewer than 50 sampled cells or a failed variance-model fit Class-specific models used 2,000 HVGs and a 30-dimensional latent space, with initialization seed 0. The global model used 5,000 HVGs, a 30-dimensional latent space and initialization seed 0. The seven class-specific analyses contained 759,259 EN, 760,855 IN, 612,708 AST, 497,329 MG, 505,823 ODC, 342,049 OPC and 360,164 VC profiles.

Training used a 90%/10% training/validation split. All models used two hidden layers of 128 units, a zero-inflated negative-binomial likelihood, library (`sample`) as the batch key, mini-batches of 1,024 cells, a target of 40,000 optimization steps and early-stopping patience of 10 epochs. For a target of $T$ steps, the maximum number of epochs was $\min\big(\max(\lceil T/s \rceil, 20), 1{,}000\big)$, where $s$ is the number of mini-batches per training epoch, and the KL warm-up was scaled to that maximum. Early stopping monitors the validation ELBO after the warm-up and restores the best checkpoint. The cell split supports optimization rather than estimation of generalization to new studies. The global model stopped early (134 of a maximum 226 epochs), whereas all class-specific models reached the epoch limit implied by the step target (60–133 epochs). Per-model epoch counts are provided in the training metadata.

#### Local inverse Simpson's index and mixing score

This section defines the permutation-normalized local inverse Simpson's index (LISI) used to quantify how well technical and biological annotations are mixed among nearest neighbors, following the entropy-calibrated weighting used in Harmony [4]. The unit of calculation is the cell, evaluated in a given representation (observed-count PCA, scVI latent coordinates, or PCA of a Kanade expression layer). For cell $i$, let $\mathcal{N}(i)$ contain its 90 nearest neighbors by exact Euclidean distance in the evaluated representation, excluding the cell itself, and let $d_{ij}$ be the distance to neighbor $j$. The neighbor weights were

$$p_{ij} = \frac{\exp(-\beta_i d_{ij})}{\sum_{l \in \mathcal{N}(i)} \exp(-\beta_i d_{il})}, \qquad j \in \mathcal{N}(i), \tag{S10}$$

where $\beta_i > 0$ was chosen by bisection so that the perplexity $\exp[-\sum_j p_{ij} \log p_{ij}]$ equalled 30, with tolerance $10^{-5}$ and at most 50 iterations. For a categorical covariate $y$ with categories $c$, the weighted category proportion in the neighborhood was $P_i(c) = \sum_{j \in \mathcal{N}(i): y_j = c} p_{ij}$, and

$$\mathrm{LISI}_i = \left[\sum_c P_i(c)^2\right]^{-1}. \tag{S11}$$

LISI equals one when all neighbors share a label and increases as more categories are represented locally. Because its attainable range depends on the number and imbalance of categories, labels were permuted once within the evaluated stratum while keeping neighborhoods and weights fixed, and the median permuted LISI, $\mathrm{median}_j(\mathrm{LISI}_j^{\mathrm{permuted}})$, was used as a reference. The normalized LISI score (mixing score) was

$$M_i = \frac{\mathrm{LISI}_i - 1}{\mathrm{median}_j(\mathrm{LISI}_j^{\mathrm{permuted}}) - 1}. \tag{S12}$$

A score of zero indicates complete label separation and a score of one indicates the median random-label reference for that covariate and stratum. Scores can exceed one when a neighborhood is more balanced than the randomly permuted global composition. High mixing is not universally desirable, because a biological covariate can represent variation that should be retained.

Covariates were library, GSE, sc/sn protocol, mesoregion, sex, Model and operational age category, defined as embryonic ($<0$ weeks), postnatal (0–$<4$), adolescent (4–$<9$), adult (9–$<44$), middle aged (44–$<78$) and aged ($\geq 78$). For the global model, neighborhoods were evaluated within major cell classes and across the full sampled cohort. Cell Ontology and major-class labels were evaluated across labeled cells, so that low scores indicate separation of cell identities. Class-specific analyses used the complete sampled cohort of that class. For each covariate, Unknown labels were excluded before recomputing neighbors, and Mix was treated as a separate sex category. The same procedure was used for observed-count PCA, scVI latent coordinates and PCA of the Kanade expression layers.

#### Microglial gene-signature scoring

This section defines the gene-signature scores used to check the biological coherence of the microglial scVI integration. Microglia were scored using the published supplementary gene-signature compilation of Green et al. [5], including the AD1/AD2 signatures of Gerrits et al. [6] and the DAM signatures of Keren-Shaul et al. [7]. Only up-regulated marker genes were used, and duplicated symbols within a signature were counted once. For cell $c$ and signature $k$ with marker set $G_k$ available in the common-gene matrix, with raw counts $x_{gc}$, the raw score was

$$s_{ck} = \frac{1}{|G_k|} \sum_{g \in G_k} \log\left(1 + 10^4 \frac{x_{gc}}{\sum_{g'} x_{g'c}}\right), \tag{S13}$$

where the denominator included all genes in the microglial common-gene count matrix. The score was centered on the median of WT cells from the same GSE when at least 50 such cells were available, and otherwise on the median of WT cells from the same sc/sn protocol. Centered scores were divided by their standard deviation across all WT cells to obtain WT-referenced z-scores. Ten signatures had at least five available up-regulated markers and were used for the main score displays: Homeostatic, AD1.c9, AD1.c10, AD2, DAM1, DAM2, Inflammation, Enhanced_redox, Stress and Proliferating. The published marker sets and available marker counts are listed in Supplementary Table S8.

Cells were divided into ten rank-based bins of Model LISI separately within WT and AD, with random tie-breaking, from the least-mixed to the most-mixed decile, and mean signature z-scores were computed using all sampled microglia in each decile. Two different dominant-signature definitions were used and should not be numerically equated. (i) For the UMAP and accompanying stacked bars, 25,000 WT and 25,000 AD cells were sampled, and the dominant signature was the maximum z-score among the ten displayed signatures, without a threshold. (ii) For the library-level composition comparisons, the dominant signature was defined over all 13 candidate signatures, and cells whose maximum z-score was below 1 were assigned to "none". For the four displayed library-level dominant-signature comparisons, two-sided Welch's $t$-test $p$-values from R were multiplied by a Bonferroni factor of ten.

#### Continuous sex covariate for Kanade

Known Male, Female and Mix labels were assigned values of 1, 0 and 0.5, respectively. For Unknown libraries, the expression-based Y/X ratio defined above was used. The posterior probability of Male from the label-anchored Gaussian model described above (fitted within sc and sn separately) supplied the continuous covariate for Unknown libraries. Among the 1,373 analyzed libraries, 373 had Unknown sex. Their probabilities had median 0.924 (interquartile range, 0.738–0.992), and 83 fell strictly between 0.1 and 0.9. These probabilities did not establish the true sex composition of a library and were not used to relabel the metadata database. Known-label agreement was 80.5% in sc and 95.3% in sn libraries. These agreement values were evaluated on the same labeled libraries used to estimate the model parameters and therefore represent in-sample concordance rather than independent validation. The sc model was fitted to 171 Male and 106 Female libraries (component means −1.829 and −3.360, common standard deviation 0.810, Male prior 0.617) and applied to 144 Unknown libraries. The sn model was fitted to 320 Male and 316 Female libraries (means −1.706 and −3.326, standard deviation 0.521, Male prior 0.503) and applied to 229 Unknown libraries. The 87 Mix libraries (71 sc and 16 sn) did not contribute to the fit.

#### Kanade representation and expression adjustment

Kanade models were fitted with initialization seeds 0, 1 and 2. The cohort contained 939 WT and 434 AD libraries, including 698 cortex, 631 hippocampus and 44 combined-tissue libraries. The 433,128 sampled profiles comprised 192,145 sc and 240,983 sn profiles. At most 500 cells were selected within a library. Validation cells were reserved within each library, with training cells retained from every library. Cells were sampled with global seed 0 and the class-specific rule of the microglial scVI analysis, using a library seed derived from the seed, library identifier and class `mg`, and 10% of the sampled cells of each library were reserved for validation, yielding 389,817 training and 43,311 validation cells . The HVG candidate filters excluded symbols beginning with `mt-` or matching the regular expression `^Rp[sl]\d`. The latter filter does not encompass every possible ribosomal-protein symbol. Age was transformed to $\log_2$ weeks, and age, tissue and sex were standardized from training-cell values. The encoder received log1p counts, and the zero-inflated negative-binomial reconstruction objective used raw counts. Posterior means were used for reconstruction. GSE was the batch variable in all three models. The 28-dimensional latent space comprised a one-dimensional scale block, supervised blocks of 4 (Model), 4 (age), 2 (tissue), 1 (sex) and 2 (sc/sn) dimensions, a 10-dimensional unsupervised block and a 4-dimensional GSE block. Model and sc/sn were predicted as categorical covariates and age, tissue and sex as continuous covariates. The reconstruction and KL weights were 1, each predictor weight and the batch weight were 30, and the independence weight was 0. Training used Adam with learning rate $10^{-3}$, mini-batches of 256 cells and at most 80 epochs. Training stopped after eight epochs without improvement in the per-cell validation sum of reconstruction, KL and predictor losses, and the best weights were restored. The realized epochs (best epoch) were 35 (27), 28 (20) and 28 (20) for seeds 0, 1 and 2, respectively. Reconstructed layers were stored as float16.

Decoded expression was converted as follows. If $\mu_g$ and $\pi_g$ are the decoder's negative-binomial mean component and zero-inflation probability for gene $g$, the library-size-independent expected abundance was $E_g = (1 - \pi_g)\mu_g$, which was converted to $\log(1 + 10^4 E_g / \sum_h E_h)$ over the 2,000 HVGs. These values are model-derived expression estimates, not integer UMI counts.

Let $\boldsymbol{z}_c^{\mathrm{sc/sn}}$ denote the sc/sn-associated latent block of cell $c$. The reference used for every adjusted cell was

$$\boldsymbol{z}_{\mathrm{ref}}^{\mathrm{sc/sn}} = (1 - p_{\mathrm{sn}})\bar{\boldsymbol{z}}_{\mathrm{sc}}^{\mathrm{sc/sn}} + p_{\mathrm{sn}}\bar{\boldsymbol{z}}_{\mathrm{sn}}^{\mathrm{sc/sn}}, \qquad p_{\mathrm{sn}} = \frac{240{,}983}{433{,}128} = 0.5564. \tag{S14}$$

The GSE-associated block was replaced by zero and the sex block by the midpoint of the Male and Female centroids. The other latent blocks were retained. The unadjusted and adjusted layers were named `R0` and `R2`. All adjusted-expression figures and numerical summaries use this `R2` definition.

#### Covariate mixing and expression fidelity

Mixing was assessed separately for each seed using the complete 433,128-cell raw, `R0` and `R2` layers. Thirty principal components were recomputed from each layer without gene-wise scaling. Exact Euclidean neighbors, the 90-neighbor entropy-calibrated kernel with perplexity 30, and the random-label normalization followed the Supplementary Methods section "Local inverse Simpson's index and mixing score". Unknown labels were removed before rebuilding the neighborhood graph for the corresponding covariate. The sex mixing statistic used original curated labels and excluded Unknown, even though continuous Unknown-sex covariates were available for model training. The visual UMAP comparison independently sampled 20,000 seed-0 cells and fitted PCA and UMAP to that subset, so it was not the space on which full-dataset LISI was computed.

Expression fidelity was evaluated across two gene sets: the 480 genes selected by the downstream consistency/effect criterion and all 2,000 HVGs. The comparison across 480 genes is conditional on that selection and does not represent all genes equally. Within a deterministically sampled subset of cells from each library, correlations were computed for raw versus `R0`, raw versus `R2`, and `R0` versus `R2`. Average ranks were used for tied values, including zero counts, and constant expression vectors were assigned undefined correlations. The paired change $\rho(\mathrm{raw}, \mathrm{R2}) - \rho(\mathrm{raw}, \mathrm{R0})$ was computed per cell before aggregation. Per-library medians were summarized across libraries, with each library receiving equal weight, separately by seed and original sc/sn protocol. Each processing library, including libraries split from a parent GSM, was treated as one aggregation unit. The maximum number of sampled cells per library was 200.

#### Age contrasts, clustering and GO over-representation

Library-level reconstructed expression was the arithmetic mean of the cell-level log-scale expression estimates. The tissue-by-age differences used the mean of the three independently trained `R2` models, while the illustrative library-level boxplots used seed 0. The boxplots pooled all retained tissue groups and combined age bands for display only. The contrast analysis retained cortex, hippocampus and combined tissue as separate strata. Within each tissue $t$ and right-open age band $a$, and separately for each Model group (WT and AD), library means were averaged within GSE and then equally across GSEs to give $\bar{x}_{g,\mathrm{Model},t,a}$ for gene $g$. The AD–WT contrast was defined as $\Delta_{g,t,a} = \bar{x}_{g,\mathrm{AD},t,a} - \bar{x}_{g,\mathrm{WT},t,a}$, with percentile intervals from 1,000 GSE bootstrap resamples. The age bands were $[2, 4)$, $[4, 9)$, $[9, 13)$, $[13, 17)$, $[17, 26)$, $[26, 35)$, $[35, 44)$, $[44, 65)$, $[65, 78)$ and $[78, 105)$ weeks. A stratum (tissue-by-age band) was supported when it contained at least two AD GSEs and two WT GSEs. Observed-count pseudobulk was obtained by summing counts within each library before normalization and log transformation. Bootstrap weights resampled GSEs from the union of studies in a stratum, preserving the pairing of WT and AD observations from the same GSE. Resamples lacking one group were discarded. Within-study contrasts first compared WT and AD from the same GSE and sc/sn protocol within a tissue–age stratum, averaged the protocol-specific contrasts within GSE, and then averaged across GSEs. No cross-study matching or latent adjustment establishes exchangeability of AD and WT observations when genotype and study are confounded.

The consistency criterion required an R2 95% bootstrap interval excluding zero, sign agreement among all three model seeds, and sign agreement with raw pseudobulk. No genome-wide multiplicity correction was applied to these gene-by-stratum intervals, and the selection was used for exploratory pattern discovery. Of 2,000 HVGs, 480 passed this criterion with absolute difference at least 0.25 in one or more of the 14 supported cortex/hippocampus strata. Each selected gene's complete 14-element vector was divided by its maximum absolute value. PCA and Ward clustering used this same matrix, with PCA centering but not scaling the columns. The three-cluster solution was chosen after inspection of the clustering results rather than by a pre-specified criterion. Cluster identifiers were assigned in descending order of gene count (203, 178 and 99 genes). For the raw-expression comparison, the same 480 genes and 14 strata were used, raw pseudobulk difference vectors were normalized independently, and a separate Ward tree was cut at three clusters. Raw and adjusted partitions were derived independently, and raw-cluster labels do not imply identity with the adjusted clusters.

GO Biological Process annotations were obtained from the MGI GAF 2.2 file (generated 2026-05-21) and the `go-basic.obo` ontology (release 2026-07-26), both downloaded on 2026-10-19. Annotations were propagated through `is_a` and `part_of` relations, retaining electronically inferred annotations and excluding `NOT` annotations and obsolete terms. Terms with 10–500 members among the input HVGs were eligible. The union of genes covered by eligible terms gave the effective 1,886-gene background. For a tested cluster, its annotated genes formed the hit set, and terms containing fewer than two hit genes were omitted before testing and adjustment. Hypergeometric probabilities were adjusted by Benjamini–Hochberg across the terms remaining after this overlap filter within each tested set (the filter changes the tested family relative to a family-wide GO FDR), with $q < 0.05$ considered significant. The effective tested hit-set sizes were 194 of 203 genes in figure cluster 1 and 173 of 178 in figure cluster 2.

##### AD-associated gene reference provenance

The candidate AD-associated labels used for gene highlighting were obtained by combining the effector-gene nominations in Supplementary Data 2 of Uffelmann et al. [8] with the AD-associated genes text file in the LRAPA datasets GitHub repository [9], which attributes its gene compilation to the ADSP Gene Verification Committee [10]. These are gene-level annotations, not a uniform set of independently causal risk variants. A risk locus can have multiple nominated genes and a nominated gene need not be expressed by the profiled cell type. The LRAPA list is a redistributed snapshot, not a live query of the committee resource. Human genes were mapped to mouse symbols by orthology. Entries with no unambiguous mouse orthologue were not coerced to similar-looking mouse symbols.

##### Gene-wise mixed-effects model and study-block validation

This section specifies the gene-wise linear mixed-effects model used to quantify age-, Model- and tissue-associated expression within the 480-gene panel selected for the descriptive AD–WT trajectory analysis. Gaussian linear mixed-effects models were fitted separately to each of the 480 genes. The input subset comprised 1,329 processing libraries from 255 GSEs (415,579 microglial cells), restricted to cortex or hippocampus libraries with usable age, Model and sc/sn annotations. The response was either observed log-normalized expression (log counts per 10,000 over the 2,000 HVGs) or the cell-level mean of the three Kanade-adjusted `R2` seeds. For gene $g$ and cell $i$ from processing library $s$, the model was

$$\begin{aligned} y_{gsi} = \beta_{0g} &+ \beta_{Ag} a_s + \beta_{Tg} t_s + \beta_{Mg} m_s \\ &+ \beta_{AT,g} a_s t_s + \beta_{AM,g} a_s m_s + \beta_{TM,g} t_s m_s \\ &+ \beta_{ATM,g} a_s t_s m_s + \beta_{Pg} p_s + b_{gs} + \varepsilon_{gsi}, \end{aligned} \tag{S15}$$

where $a_s = \log_{10}(\text{age}_s/20\,\text{weeks})$; $t_s = -0.5$ for cortex and $+0.5$ for hippocampus; $m_s = -0.5$ for WT and $+0.5$ for AD; and $p_s = -0.5$ for sc and $+0.5$ for sn. The library-specific intercept $b_{gs}$ and the residual $\varepsilon_{gsi}$ were assumed Gaussian. The full three-way interaction of age, tissue and Model was fitted by restricted maximum likelihood (REML), with an additive sc/sn term. The centering at 20 weeks and the effect coding make lower-order terms refer to the average over the effect-coded groups at the age reference point, subject to the model's linearity assumptions. The age-by-AD interaction ($\beta_{AM,g}$) estimates the difference in log-age slopes between AD and WT, averaged across cortex and hippocampus at the coding reference, rather than the expression difference at a specific age. The eight non-intercept terms were age, tissue, Model, the three two-way interactions, the three-way interaction and sc/sn protocol.

The fit used library-specific sufficient statistics $(n, \bar{y}, \text{SSW})$, which preserve the within-library sums of squared deviations needed for the Gaussian random-intercept likelihood, and was validated against an independent mixed-model implementation (`statsmodels`).

For inferential checks, the model-based Wald $p$ value was distinguished from the GSE-cluster-robust CR1 $p$ value ($p_{\text{GSE}}$). For the latter, the covariance sandwich was constructed by GSE cluster and a $t$ reference with $G-1$ degrees of freedom was used (where $G$ counts GSE clusters). Benjamini–Hochberg adjustment was performed separately for each model term and expression layer across the 480 selected genes (16 testing families, used for Fig. 4e,f and the principal gene-count summaries), and genes with $q_{\text{BH}} < 0.05$ were marked as significant. Standard model-based Wald tests using the library random effect alone were retained as a sensitivity analysis ($p$) but were not used for the primary significance calls. Bonferroni-adjusted values ($32{,}250 \times p_{\text{GSE}}$, capped at one) were used as a stringent sensitivity comparison. Supplementary Table S1 reports BH gene counts with GSE-robust Bonferroni counts in parentheses as a stringent sensitivity comparison. The BH testing families were restricted to a data-selected panel and not pooled across terms or layers. Because the genes were preselected from the same material and the permutation audit indicated some $p$-value inflation, the adjusted values should be interpreted as exploratory, not a demonstration of unconditional FDR control.

Null calibration used 100 permutations per representation and 480 genes, shuffling age and Model within GSEs and additionally permuting study-level means across GSEs. The variance components from the full-data fit were held fixed and the Kanade encoder/decoder was not retrained. The test addresses the downstream LMM procedure conditional on a frozen representation, not the validity of the full supervised reconstruction pipeline. The corrected null results are reported in Supplementary Table S2. Null rejection proportions at $p_{\mathrm{GSE}} < 0.05$ were 0.061–0.104, compared with 0.27–0.39 for library-only tests. These proportions describe the 480-gene selected analysis, rather than an external assessment of genome-wide error rates.

For coefficient-concordance analysis, Spearman correlations between raw and R2 estimates were 0.67 for age, 0.68 for the AD main effect and approximately 0.2–0.3 for the interactions. For age-by-AD specifically, 100 genes were BH-significant in R2 and nine in raw, with six genes significant in both representations. Of the 100 R2 hits, 90 had the same coefficient sign in raw, and 49 had raw nominal $p_{\mathrm{GSE}} < 0.05$ with the same direction. Among R2 study-robust significant genes, age coefficient signs agreed with raw for 97%, and AD coefficient signs agreed for 87%. The corresponding median R2/raw absolute coefficient ratios were approximately 2.3 (age) and 5 (AD) over R2-significant genes. R2 also reduced the reported library-intercept standard deviation from about 0.35 in raw to 0.24, without establishing whether this reflects denoising rather than decoder smoothing. For leave-one-GSE-out analysis, the full-fit variance components were held fixed, so this is not a complete study-level re-fit or a re-training of Kanade. No sign reversals occurred among the R2 GSE-robust *Bonferroni*-significant AD (283), age (112) or age-by-AD (5) terms under this restricted sensitivity analysis. This leave-one-GSE-out audit was not rerun on all BH-significant genes.

A separate GO Biological Process hypergeometric analysis was performed on the stricter GSE-robust *Bonferroni*-significant LMM genes (`q_gse`<0.05), rather than the newly defined BH-significant sets, using the 480 selected genes as the GO background and BH adjustment within each tested gene set. This is different from the three-cluster enrichment analysis, whose effective background contained 1,886 annotated HVGs. Among the 283 R2 AD-associated genes, 185 had positive coefficients and 98 negative. Among the 112 age-associated genes, 93 were positive and 19 negative. The AD-positive set contained DAM-associated, phagolysosomal, antigen-presentation and interferon-response genes and enriched broad immune functions (reported adjusted GO $q$ approximately 0.002–0.004). The age-positive set included inflammation- and antigen-presentation-associated genes but did not yield significant GO enrichment after adjustment (best reported $q$ approximately 0.08). Many negative-coefficient genes were neuronal, oligodendroglial, haemoglobin or vascular-associated, and the inferred reduction in their normalized counts may arise from ambient RNA, cross-cell contamination or compositional effects rather than regulation in microglia. The single-gene assignments used to describe these pathways were manually inspected, not independently curated functional annotations.

### Supplementary Tables

**Table S4: Schema of the harmonized GSM-level metadata.** Source links and curatorial notes are retained where available. They are not a complete field-level evidence record.

| Field | Values or format | Meaning |
|---|---|---|
| `GSE` | GEO series accession | Study-level source identifier. |

Continued on next page

Table S4 continued

| Field | Values or format | Meaning |
|---|---|---|
| GSM | GEO sample accession or local placeholder | Parent sample identifier. GSM000003–GSM000014 in this curated table are local grouping identifiers rather than GEO accessions. |
| GSE_link | GEO study URL | Link to the source GEO series. |
| Study_link | Publication URL or missing marker | Publication link retained from curation. This is not a complete per-field evidence record. |
| Study_title | Study title | Human-readable study title retained from curation. |
| Region | Curated anatomical description | Retains anatomical detail used in curation, including combined-region descriptions. |
| UBERON_ID | One or more UBERON IDs | Anatomical ontology terms. Multiple IDs are separated by semicolons. |
| scsn | sc / sn | Whole-cell or isolated-nucleus protocol, based on curated source descriptions. |
| Platform | 10X | Target platform label retained for this collection. |
| Strain | Curated strain text | Mouse genetic background when available. |
| Genotype | Full curated genotype | Detailed genotype including reported disease alleles, drivers, reporters and other tool alleles. |
| Allele | Curated allele text | Additional allele details retained from curation. |
| Generalized_genotype | Generalized genotype text | Disease-relevant or control-group genotype representation before final display standardization. |
| Condition | Curated experimental context | Condition information retained from curation. Not interchangeable with Model or Treatment. |
| Treatment | Curated treatment text | Detailed administration, control or manipulation, including unknown or unreported values. |
| Model | WT / AD | Broad analysis group. WT includes relevant controls, tool genotypes and non-AD knock-ins and does not imply an unmodified or untreated animal. |
| Note | Curator note text | Available corrections, evidence descriptions and screening notes. These heterogeneous notes are not a complete field-level provenance log. |
| Mesoregion | Cortex / Hippocampus / Cortex;Hippocampus | Coarse anatomical analysis group. Combined regions do not imply equal tissue proportions. |
| Generalized_genotype_std | Standardized label | Label after selected spelling and capitalization mappings and WT tool-allele grouping. Labels are not guaranteed to define distinct biological models. Other includes unresolved genotypes. |
| Treatment_std | None / Control / Tamoxifen / Tag / Viral_tracing / Viral_reporter / Viral_Cre / Viral_WT_gene_KI / Tau | Coarse treatment category. None also includes unknown or unreported descriptions and some routine housing/preparation entries. Vehicle controls are included in Control. |
| DL_status | Target / Non-target | Availability of usable, sample-assignable counts for expression processing. Every row is biologically eligible for the metadata resource. Non-target here does not reverse that inclusion decision. |

Continued on next page

Table S4 continued

| Field | Values or format | Meaning |
|---|---|---|
| Sex | Female / Male / Mix / Unknown | Curated sex. Mix represents pooling of both sexes. Unknown individual sample sex remains Unknown. |
| Age | Numeric weeks / Unknown | Curated collection age in weeks. Range midpoints are used for bounded intervals. Embryonic ages are represented relative to E21. |

**Table S5: Metadata fields used for Gemma screening.** GSE/GSM identifier columns were omitted, but identifiers embedded in the free-text values were not systematically removed.

| Level | Input field | Description |
|---|---|---|
| Study | Study_title | GEO series title |
| Study | Overall_summary | Series summary |
| Study | Overall_design | Series experimental design |
| Sample | Characteristics | GSM characteristics |
| Sample | GSM_title | GSM title |
| Sample | GSM_source | Sample source name |
| Sample | Growth_protocol | Growth or preparation description |
| Sample | Treatment_protocol | Treatment description |
| Sample | Extract_protocol | Extraction description |

**Table S1: Gene-wise mixed-effects coefficients passing multiplicity adjustment.** Counts are genes among 480 selected genes with GSE-cluster-robust $p$-values BH-adjusted *separately* for each model term and expression layer ($q_{\mathrm{BH}} < 0.05$). Parenthetical numbers give GSE-cluster-robust Bonferroni counts (factor 32,250), not GSM-only counts. Tissue denotes the hippocampus-minus-cortex effect, AD denotes the AD-minus-WT effect and age uses $\log_{10}(\text{age}/20\,\text{weeks})$.

| Model term | Adjusted R2 | Observed raw |
|---|---|---|
| AD | 379 (283) | 92 (11) |
| Age | 259 (112) | 65 (3) |
| Hippocampus | 208 (75) | 52 (1) |
| Age × AD | 100 (5) | 9 (0) |
| Age × Hippocampus | 76 (9) | 0 (0) |
| Hippocampus × AD | 70 (14) | 0 (0) |
| Age × Hippocampus × AD | 23 (0) | 0 (0) |
| Single nucleus (sn) | 103 (0) | 405 (284) |

**Table S2: Null calibration of the LMM inference in the selected 480-gene panel.** Null rejection proportions from 100 GSE-block permutations at nominal unadjusted $p$ thresholds. Comparisons use fixed variance components and an unchanged Kanade representation.

| Layer | Term | GSM $p < 0.05$ | GSE $p < 0.05$ | GSE $p < 0.01$ |
|---|---|---|---|---|
| R2 | AD | 0.370 | 0.070 | 0.016 |
| R2 | Age | 0.380 | 0.061 | 0.018 |
| R2 | Age × AD | 0.390 | 0.104 | 0.030 |
| Raw | AD | 0.330 | 0.082 | 0.021 |
| Raw | Age | 0.370 | 0.077 | 0.020 |
| Raw | Age × AD | 0.270 | 0.086 | 0.024 |

**Table S3: Software and computational environment.**

| Stage | Software or recorded provenance |
|---|---|
| Metadata retrieval | R 4.3.2 and GEOquery 2.70.0. |
| Screening | Gemma 4 31B-it (Hugging Face snapshot `842da37`) with offline vLLM 0.25.1, Python 3.12.13, PyTorch 2.11.0 and Transformers 5.14.0, and Claude Opus 5.5 for prompt revision. |
| Read reprocessing | scRecounter (Docker image sc-recounter-star 0.1.0; STARsolo, STAR 2.7.11b) with GeneFull for all 414 reprocessed GSMs. |
| Gene harmonization | Mouse Ensembl BioMart mapping, retrieved on 2026-10-14. |
| Cell calling | STAR 2.7.11b, EmptyDrops_CR. |
| QC and annotation | Python 3.12.3, Scanpy 1.11.5 (including its Scrublet implementation), AnnData 0.12.7, leidenalg 0.11.0, python-igraph 1.0.0, scikit-image 0.26.0, ddqc_scanpy (vendored source, BSD-3-Clause) with a mouse mitochondrial/ribosomal gene list extracted from GENCODE vM38 (GRCm39), and the CMA mouse healthy-tissue database v1.0.0. |
| scVI | Python 3.12.0, scvi-tools 1.5.1, PyTorch 2.11.0, Lightning 2.6.6, Scanpy 1.11.5, AnnData 0.12.6, NumPy 2.3.5, SciPy 1.16.3, pandas 2.3.3, scikit-misc 0.5.2, umap-learn 0.5.9.post2 and pynndescent 0.5.13. |
| Kanade | Kanade commit `2addc42` with project-specific runtime adaptations, Python 3.11.17, PyTorch 2.6.0, AnnData 0.12.19 and NumPy 2.4.6. Run configuration in Supplementary Methods. |
| Original figures | R 4.3.2, ggplot2 3.4.4 and patchwork 1.1.3. |

**Table S6: Markers used for the CMA internal-consistency display.** The five markers per class were selected from the CMA resource using the library-frequency and feature-coverage rules described in the Supplementary Methods.

| Class | Selected markers |
|---|---|
| EN | *Sv2b*, *Bcl11b*, *Nell1*, *Cdh13*, *Nell2* |
| IN | *Erbb4*, *Kcnc1*, *Gad2*, *Col25a1*, *Cnr1* |
| OPC | *Pdgfra*, *Kcnip3*, *Olig1*, *Tnr*, *C1ql1* |
| AST | *Slc1a3*, *Gja1*, *Gfap*, *Acsbg1*, *Slc1a2* |
| MG | *C1qc*, *C1qb*, *Ctss*, *P2ry12*, *Cx3cr1* |
| ODC | *Mog*, *Plp1*, *Mbp*, *Aspa*, *Opalin* |
| VC | *Flt1*, *BC028528*, *Col4a1*, *Esam*, *Cd93* |

**Table S7: Mapping of candidate CMA Cell Ontology terms to major classes.** Candidate terms need not occur among final assignments. Unknown and other ungrouped annotations were summarized as Others. Tissue-specific marker availability and eligibility are retained in the source CSV.

| Major class | Cell Ontology identifier | CMA cell type |
|---|---|---|
| AST | CL:0000127 | astrocyte |
| AST | CL:0002627 | mature astrocyte |
| EN | CL:4030059 | L2/3 intratelencephalic projecting glutamatergic neuron |
| EN | CL:4030067 | L5/6 near-projecting glutamatergic neuron |
| EN | CL:4023042 | L6 corticothalamic-projecting glutamatergic cortical neuron |
| EN | CL:4030065 | L6 intratelencephalic projecting glutamatergic neuron |
| EN | CL:4023038 | L6b glutamatergic cortical neuron |
| EN | CL:2000089 | dentate gyrus granule cell |
| EN | CL:0000679 | glutamatergic neuron |
| EN | CL:0000598 | pyramidal neuron |
| IN | CL:0000617 | GABAergic neuron |
| IN | CL:4023075 | L6 th sst GABAergic interneuron (Mmus) |
| IN | CL:4023016 | VIP GABAergic cortical interneuron |
| IN | CL:0000498 | inhibitory interneuron |
| IN | CL:0000099 | interneuron |
| IN | CL:4023011 | lamp5 GABAergic cortical interneuron |
| IN | CL:4023018 | pvalb GABAergic cortical interneuron |
| IN | CL:4023015 | sncg GABAergic cortical interneuron |
| IN | CL:4023017 | sst GABAergic cortical interneuron |
| IN | CL:4023121 | sst chodl GABAergic cortical interneuron |
| Others | CL:0000235 | macrophage |
| MG | CL:0000129 | microglial cell |
| Others | CL:0000881 | perivascular macrophage |
| ODC | CL:0000128 | oligodendrocyte |
| OPC | CL:0002453 | oligodendrocyte precursor cell |
| Others | CL:0000084 | T cell |
| Others | CL:0000066 | epithelial cell |
| Others | CL:0011012 | neural crest cell |
| Others | CL:0000047 | neural stem cell |
| Others | CL:0000031 | neuroblast (sensu Vertebrata) |
| Others | CL:0000540 | neuron |
| Others | CL:0011026 | progenitor cell |
| Others | CL:0000681 | radial glial cell |
| VC | CL:0000115 | endothelial cell |
| Others | CL:1001045 | kidney cortex artery cell |
| VC | CL:0000669 | pericyte |

Continued on next page

Table S7 continued

| Major class | Cell Ontology identifier | CMA cell type |
|---|---|---|
| VC | CL:4023056 | vascular leptomeningeal cell (Mmus) |

**Table S8: Microglial gene signatures used in this analysis.** Exact mouse-symbol sets follow the published supplementary compilation of Green et al. [5]. AD1/AD2 and DAM originate from Gerrits et al. [6] and Keren-Shaul et al. [7], respectively. Input counts use unique symbols, and available counts are recorded in the analysis output. Ten sets with at least five available markers were used for the main score displays. **Bold signature names** indicate the ten signatures used for the main display, and **bold italic gene symbols** indicate the used marker genes. All 13 candidate sets entered the separate dominant-signature comparison. The repeated Hspa1b entry in the deposited Stress set was counted once.

| Signature | Input | Available | Up-regulated marker set |
|---|---|---|---|
| **AD1.c10** | 7 | 7 | ***Cpm***, ***Gpnmb***, ***Apoe***, ***P2ry12***, ***Stard13***, ***Cx3cr1***, ***Eya2*** |
| AD1.c7 | 4 | 4 | *Dscam*, *Socs6*, *Adamts17*, *Ipcef1* |
| **AD1.c9** | 7 | 5 | ***Dtna***, ***Myo1e***, ***Lpl***, ***Spats2l***, ***Gldn***, *Tprg*, *Itgax* |
| **AD2** | 6 | 5 | ***Foxp2***, ***Dpp10***, ***Grid2***, ***Ddx17***, ***Nav2***, *Adgrb3* |
| **DAM1** | 9 | 7 | ***Ctsb***, ***Fth1***, ***Apoe***, ***B2m***, ***Timp2***, ***Tyrobp***, ***Cstb***, *9530003J23Rik*, *Gm49369* |
| **DAM2** | 17 | 15 | ***Spp1***, ***Cd9***, ***Hif1a***, ***Axl***, ***Serpine2***, ***Cadm1***, ***Lpl***, ***Cd52***, ***Ank1***, ***Cd63***, ***Csf1***, ***Cst7***, ***Gusb***, ***Ctsz***, ***Ctsa***, *Clec7a*, *Itgax* |
| **Enhanced_redox** | 9 | 8 | ***Fth1***, ***Rpl19***, ***Apoe***, ***C1qa***, ***Rps6***, ***C1qb***, ***C1qc***, ***Rps19***, *Ftl1-ps1* |
| **Homeostatic** | 7 | 7 | ***Sox5***, ***P2ry12***, ***Sh3rf3***, ***Cx3cr1***, ***Tmem163***, ***Frmd4a***, ***Syndig1*** |
| **Inflammation** | 8 | 8 | ***Srgn***, ***Gpr183***, ***Cd83***, ***Gpnmb***, ***Nfkb1***, ***Ccl4***, ***Il1b***, ***Ccl3*** |
| Interferon Response | 2 | 1 | *Ifi44l*, *Ifit3* |
| **Proliferating** | 6 | 6 | ***Top2a***, ***Fanci***, ***Hmmr***, ***Brip1***, ***Mki67***, ***Arhgap11a*** |
| **Stress** | 6 | 6 | ***Hsp90aa1***, ***Fos***, ***Gpr183***, ***Srgn***, ***Hspa1b***, ***Junb*** |
| Stress_response/anti-inflammatory | 3 | 3 | *Spp1*, *Tmem163*, *Ddit4* |

### Supplementary Figures

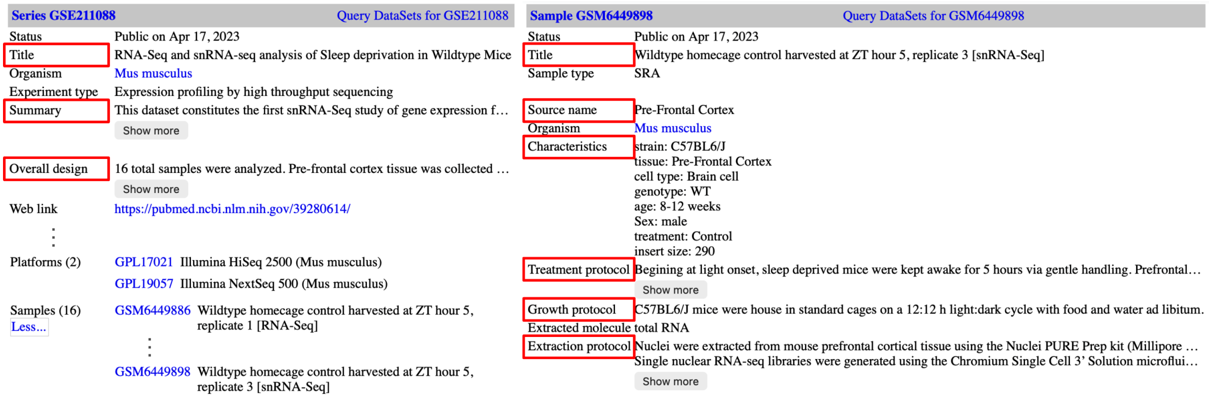


**Figure S1: Target descriptions in GSE and GSM.** Text values at the highlighted attribute keys of the example GSM were used for LLM information extraction and evaluation.

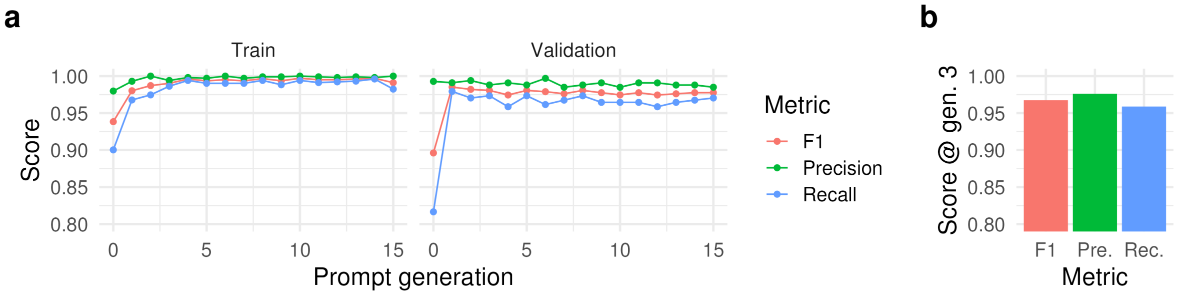


**Figure S2: Performance of LLM-assisted sample screening.** (a) Precision, recall and $F_1$ for all 16 prompt generations on the training and validation partitions. (b) The frozen generation-3 prompt on the study-separated held-out test set of 676 GSMs.

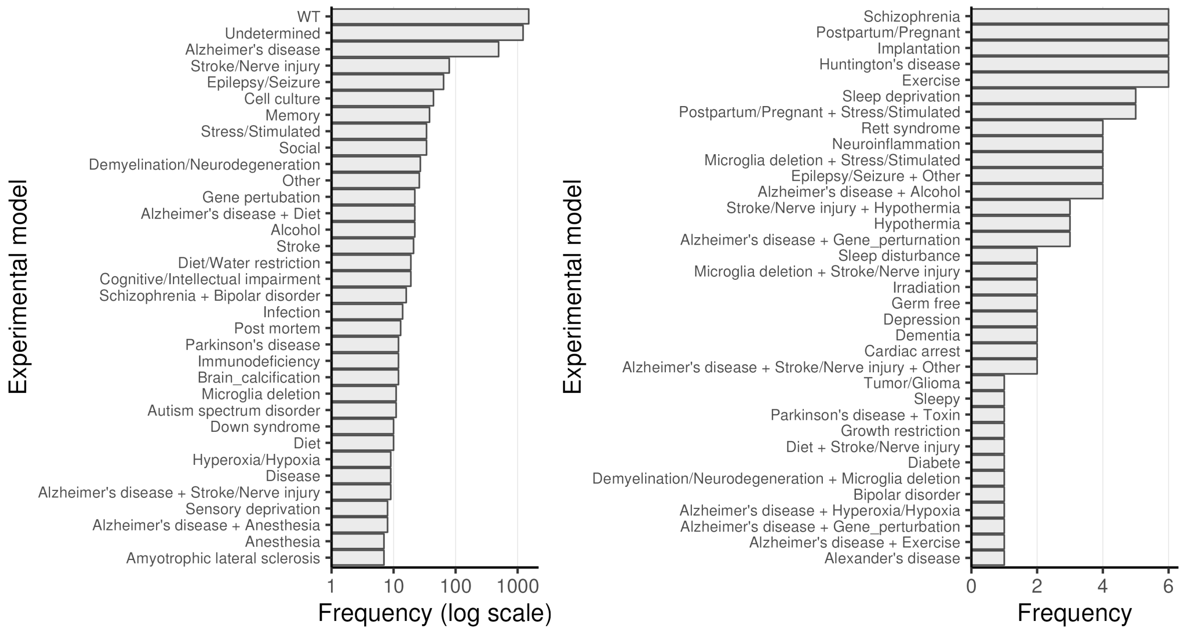


**Figure S3: Experimental-model annotations in the broader screening inventory.** Frequencies of manually annotated model categories before restriction to WT/control and AD-related models. Common categories are displayed on a logarithmic axis and less frequent categories on a linear axis. Combined labels preserve multiple experimental contexts. These frequencies describe the broader inventory and are not the composition of the final 2,005-record metadata resource.

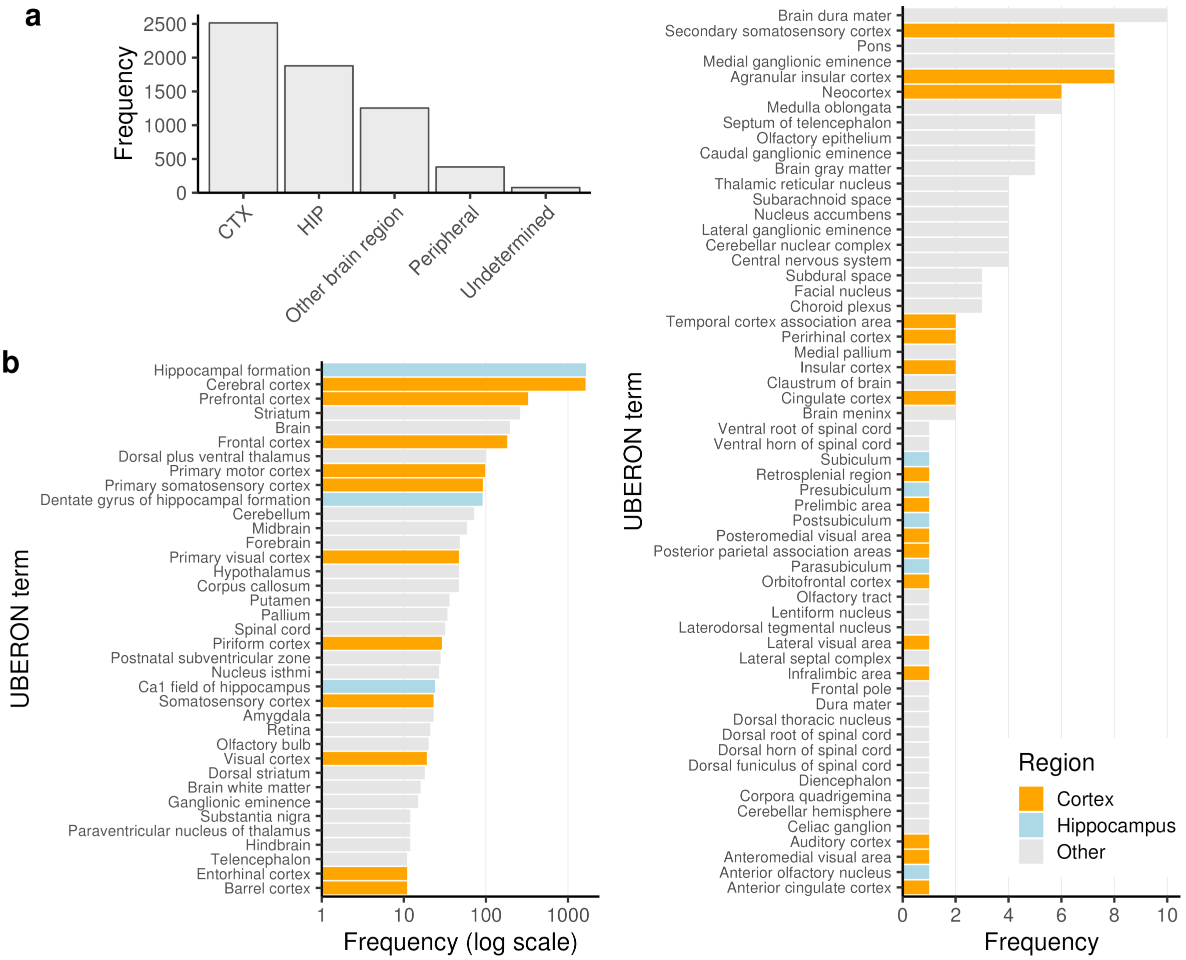


**Figure S4: Anatomical annotations in the broader manually annotated screening inventory.** (a) Frequencies of broad anatomical assignments before restriction to the final cortex/hippocampus WT/AD cohort. (b) Frequencies of detailed UBERON terms, with common terms on a logarithmic axis and less frequent terms on a linear axis. Cortex- and hippocampus-associated terms are highlighted. One record can contribute multiple anatomical terms, so term frequencies are not mutually exclusive sample counts and should not be summed to estimate the final cohort size.

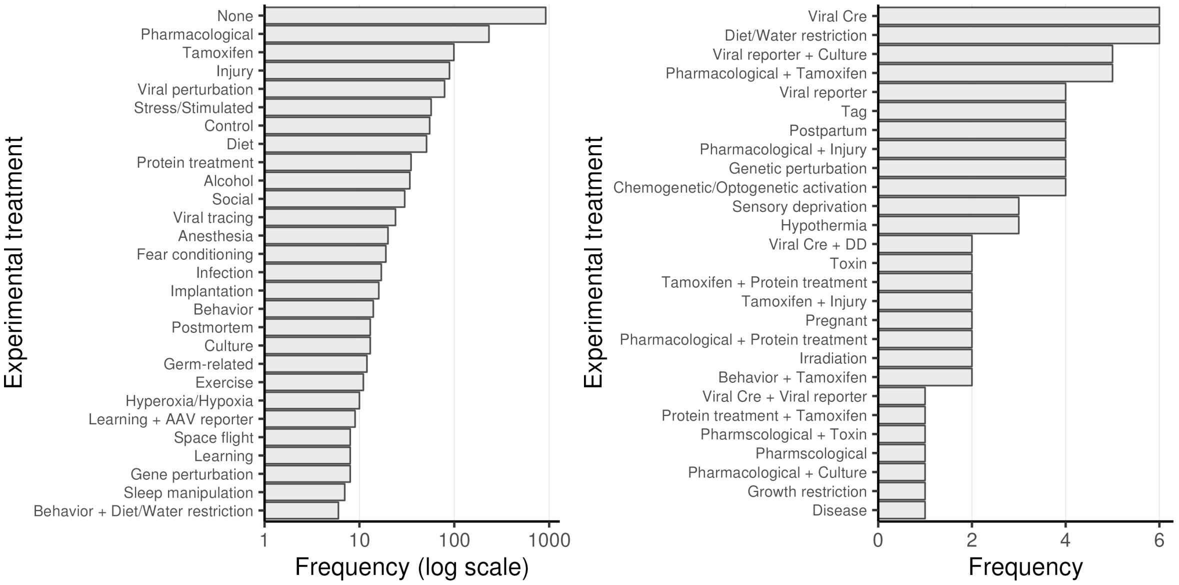


**Figure S5: Treatment annotations outside the final WT/AD grouping.** Treatment categories for manually annotated records assigned to other experimental-model groups. Common categories are displayed on a logarithmic axis and less frequent categories on a linear axis. Combined labels indicate multiple recorded interventions. None includes unspecified or unreported treatment descriptions and is not evidence that every contributing animal was untreated.

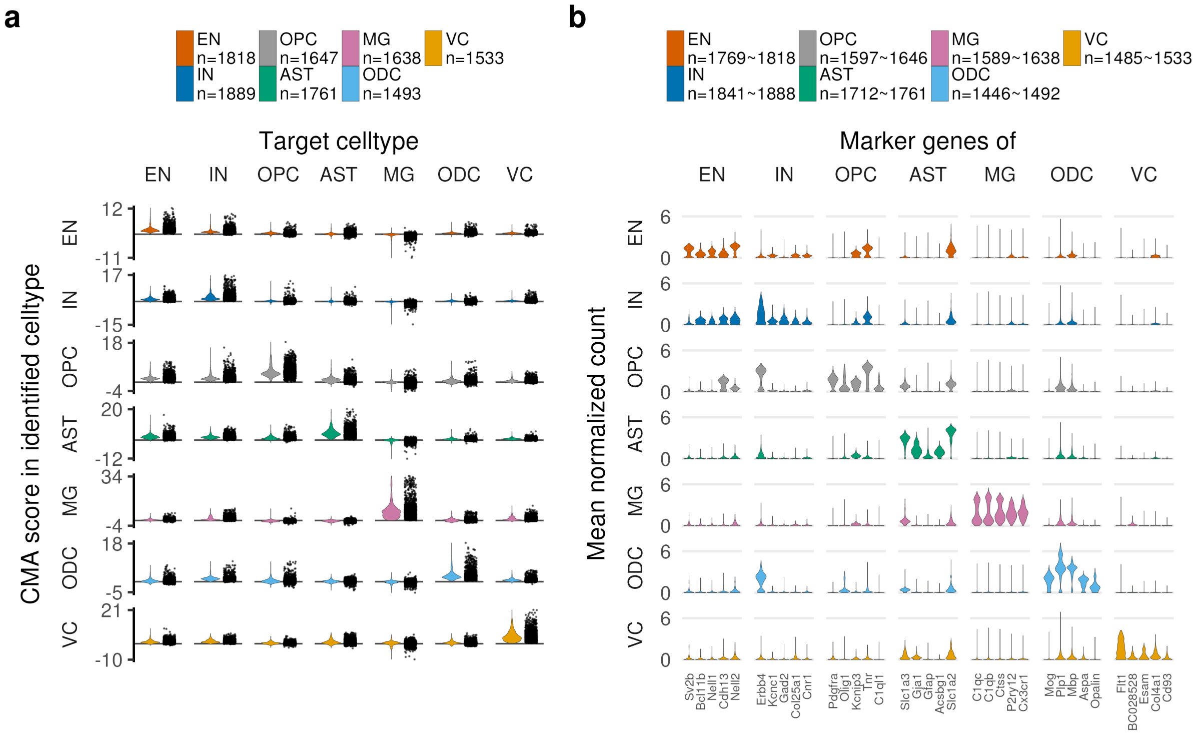


**Figure S6: Internal consistency of CMA annotations with scores and marker expression.** (a) For each identified major class and target marker class, the library-level cell-count-weighted median of the best target-class 75th-percentile CMA score in each assigned cluster. Rows indicate identified classes and columns indicate target marker classes. (b) Mean cell-level log1p-normalized expression of five selected CMA markers per target class, summarized within each library and identified class. Only library–class combinations with at least ten cells are used in the marker display, and missing genes are excluded. Marker selection uses the same CMA resource as cell assignment, so the displays evaluate internal consistency rather than independent classification accuracy.

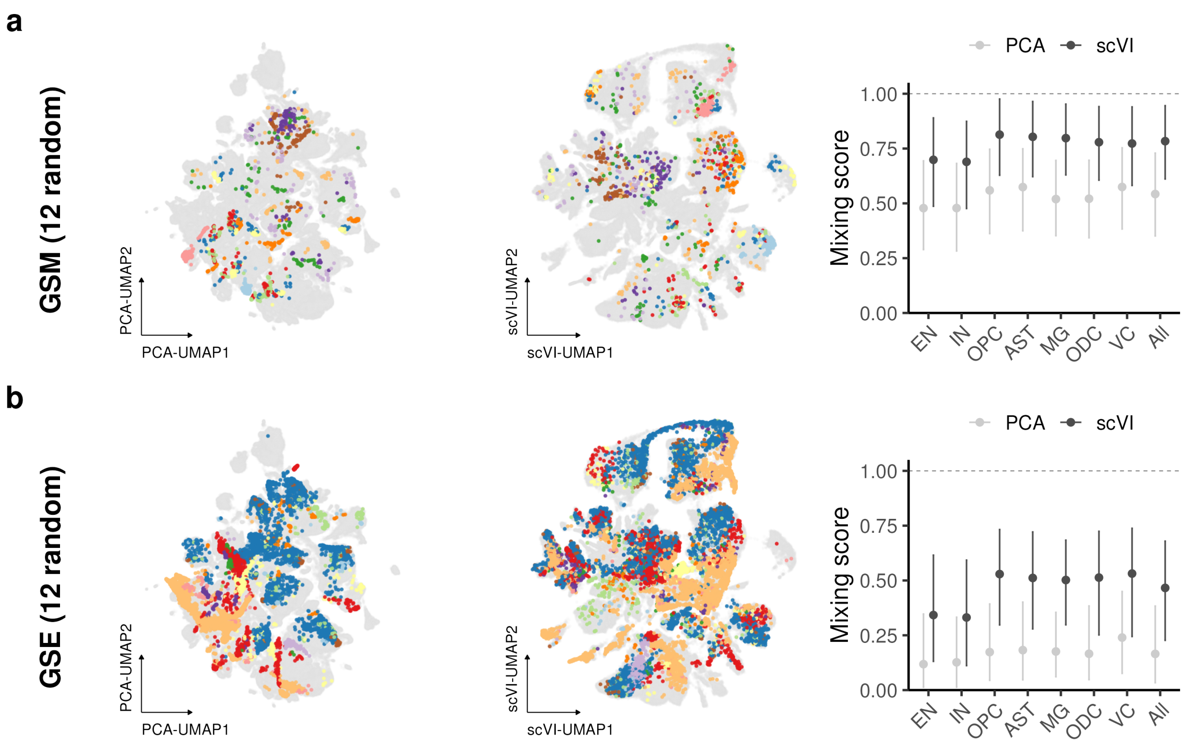


**Figure S7: Study and library organization before and after global scVI integration.** (a) PCA- and scVI-derived UMAPs of 200,425 sampled profiles, highlighting 12 randomly selected libraries, and median library mixing scores with interquartile ranges. (b) Corresponding displays for 12 randomly selected GSEs. Mixing scores were calculated using exact neighborhoods within each class and, separately, across all cells. The dashed line marks the random-label reference of one. GSM labels denote processing-library identifiers, including split GSM_k entries.

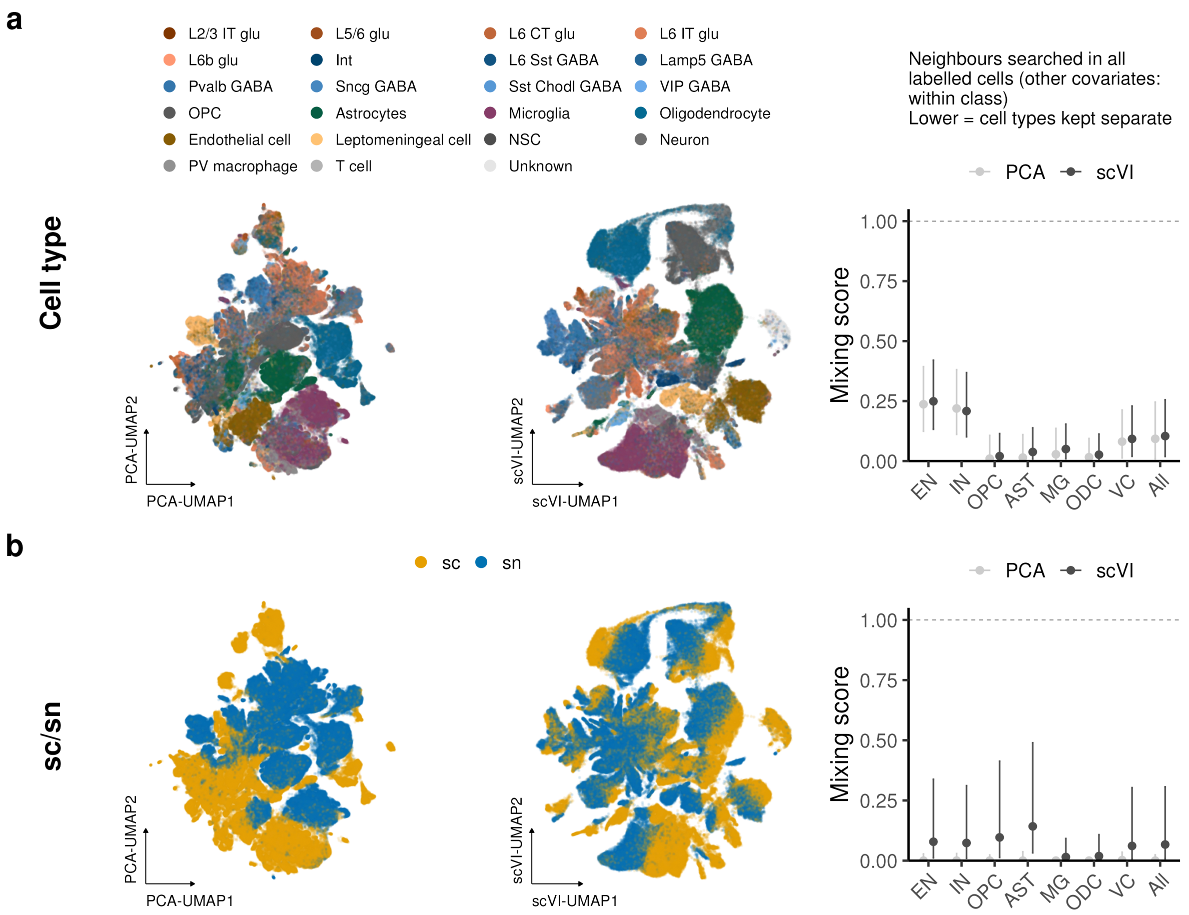


**Figure S8: Cell-type and sc/sn organization in the global PCA and scVI representations.** (a) UMAPs colored by Cell Ontology annotation and mixing of those labels. Cell-type neighborhoods are evaluated across labeled cells, and lower mixing indicates separation of the annotated types. (b) UMAPs and mixing summaries for sc/sn protocol. Points and bars indicate medians and interquartile ranges, and the dashed line denotes the random-label reference of one. The input contains 200,425 sampled profiles.

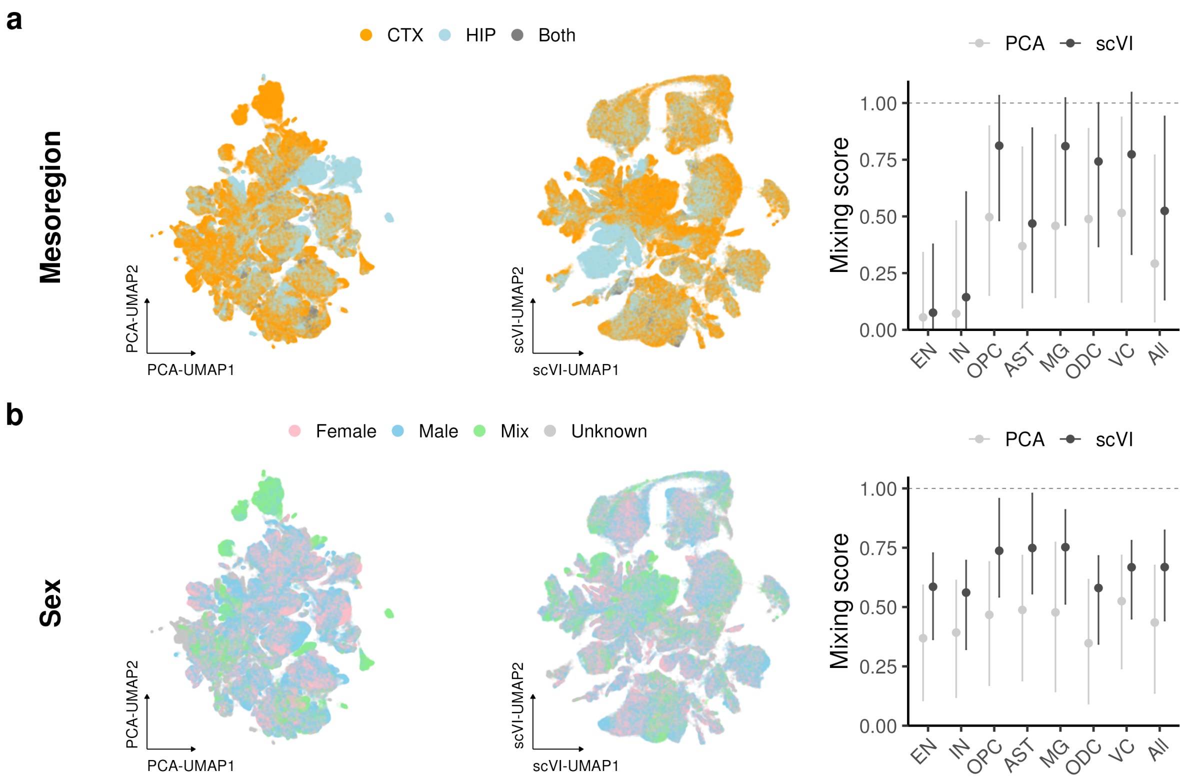


**Figure S9: Mesoregion and sex organization in the global PCA and scVI representations.** (a) UMAPs and permutation-normalized LISI summaries for cortex, hippocampus and combined-tissue labels. (b) Corresponding displays for sex, with Unknown excluded from the mixing calculation and Mix treated as a separate category. Points and bars indicate medians and interquartile ranges.

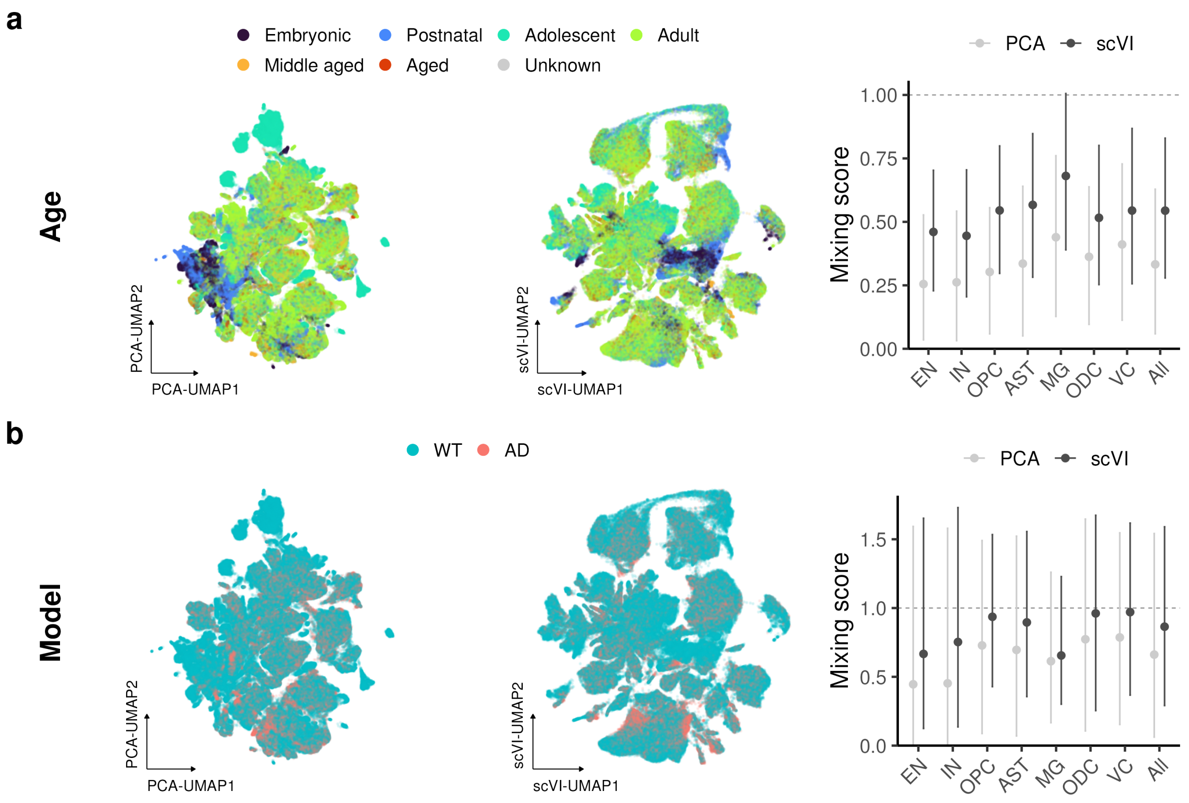


**Figure S10: Age and WT/AD organization in the global PCA and scVI representations.** (a) UMAPs and permutation-normalized LISI summaries for operational age categories. Operational age categories (defined in the Supplementary Methods section on the local inverse Simpson's index). Unknown ages are excluded. (b) Corresponding displays for WT/AD Model.

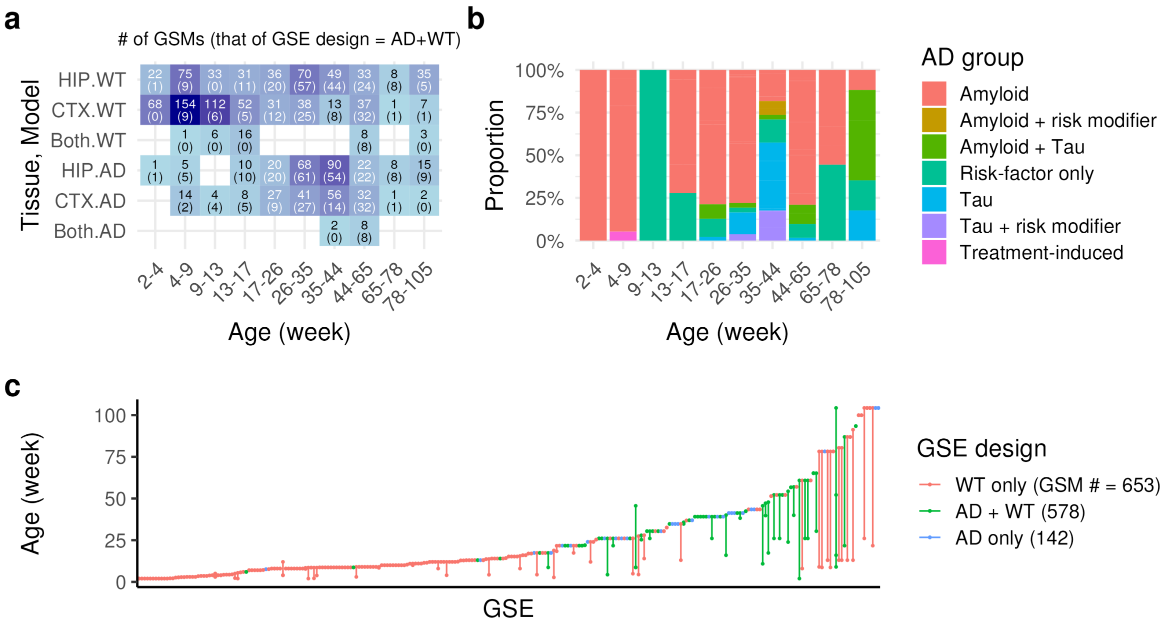


**Figure S11: Study design of the exploratory Kanade cohort.** (a) Numbers of libraries by tissue, Model and original age band. Parenthetical numbers count libraries from GSEs that contain both WT and AD somewhere in the analysis cohort. They do not count GSEs or require both groups within the particular tile. (b) Relative abundance of AD genotype categories by age band, pooling all retained tissue groups. (c) Age ranges per GSE–tissue combination, ordered by maximum age and colored by whether the GSE contains WT only, AD only or both. Lines join the observed youngest and oldest ages and do not indicate continuous coverage. Legend counts refer to libraries: 653 from WT-only GSEs, 142 from AD-only GSEs and 578 from GSEs with both groups.

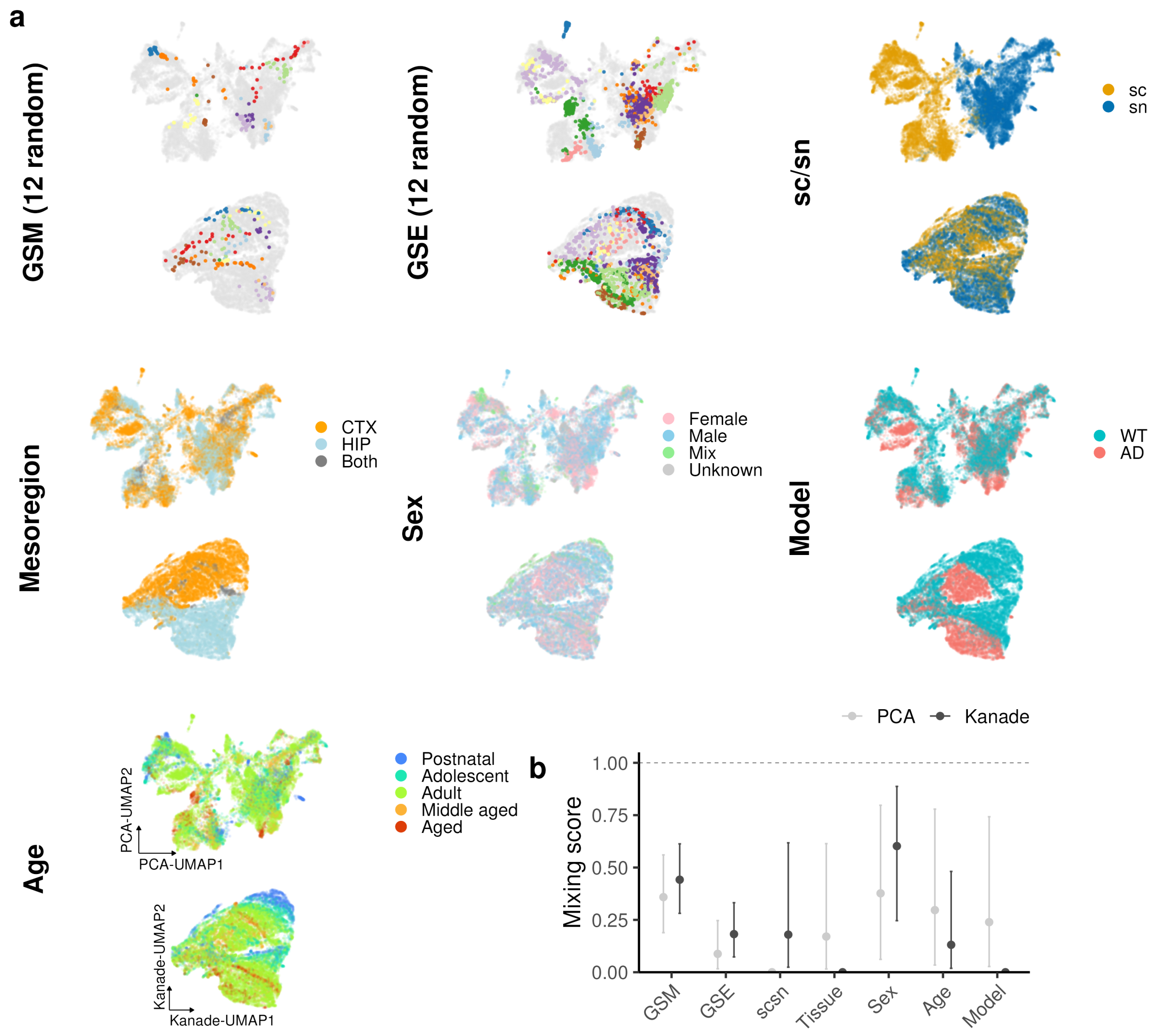


**Figure S12: Covariate organization before and after Kanade adjustment.** (a) Separate UMAP projections of raw and adjusted R2 expression for 20,000 sampled cells from seed 0, colored by the indicated metadata. Twelve randomly selected library or GSE labels are highlighted against the remaining profiles. Each UMAP was fitted independently to the displayed subset. (b) Median permutation-normalized LISI mixing scores from PCA of the full 433,128-cell raw and R2 representations, with interquartile ranges across cells. The dashed line is the random-label reference of one. Unknown sex labels were excluded from sex mixing, although a continuous sex covariate was estimated for model training. Model, age and tissue were supervised training covariates, so their separation is not independent validation of the reconstructed expression.

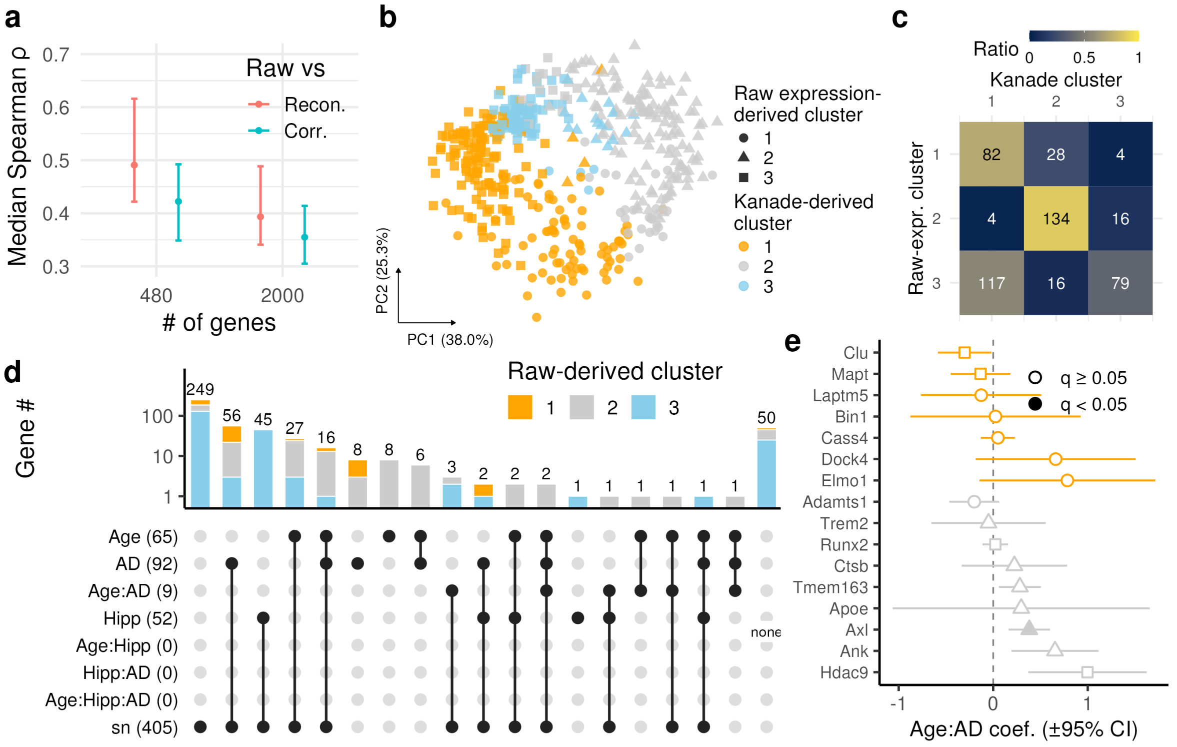


**Figure S13: Expression fidelity and dependence of gene clustering on the expression representation.** (a) Median and interquartile range across libraries of within-library median cell-level Spearman correlations between raw and reconstructed `R0` expression (Recon.) or adjusted `R2` expression (Corr.), for the 480 selected genes and all 2,000 HVGs (seed 0). (b) PCA of raw pseudobulk AD–WT profiles for the same 480 genes and 14 strata used in the adjusted-expression analysis. Colors indicate the adjusted-expression cluster and symbols the independently derived raw-expression cluster. (c) Cross-tabulation of raw- and R2-derived gene clusters. Numbers indicate gene counts, and colors represent within-row proportions. (d) UpSet display of overlaps among mixed-model terms for observed-count pseudobulk expression, using clusters derived independently from raw expression. The Y axis is logarithmic. Significance uses within-term BH adjustment of GSE-cluster-robust $p$-values over the 480-gene panel ($q_{\mathrm{BH}} < 0.05$). (e) Gene-wise log-age-by-AD interaction estimates for AD-reference-list genes in the observed-count raw layer. Point shapes indicate the raw-expression cluster, colors the R2-derived cluster and point fill indicates within-term BH-adjusted GSE-robust significance ($q_{\mathrm{BH}} < 0.05$). Error bars are approximate 95% intervals $\hat{\beta} \pm 1.96\,\mathrm{SE}_{\mathrm{GSE}}$. The corresponding adjusted-layer results appear in Fig. 4f.

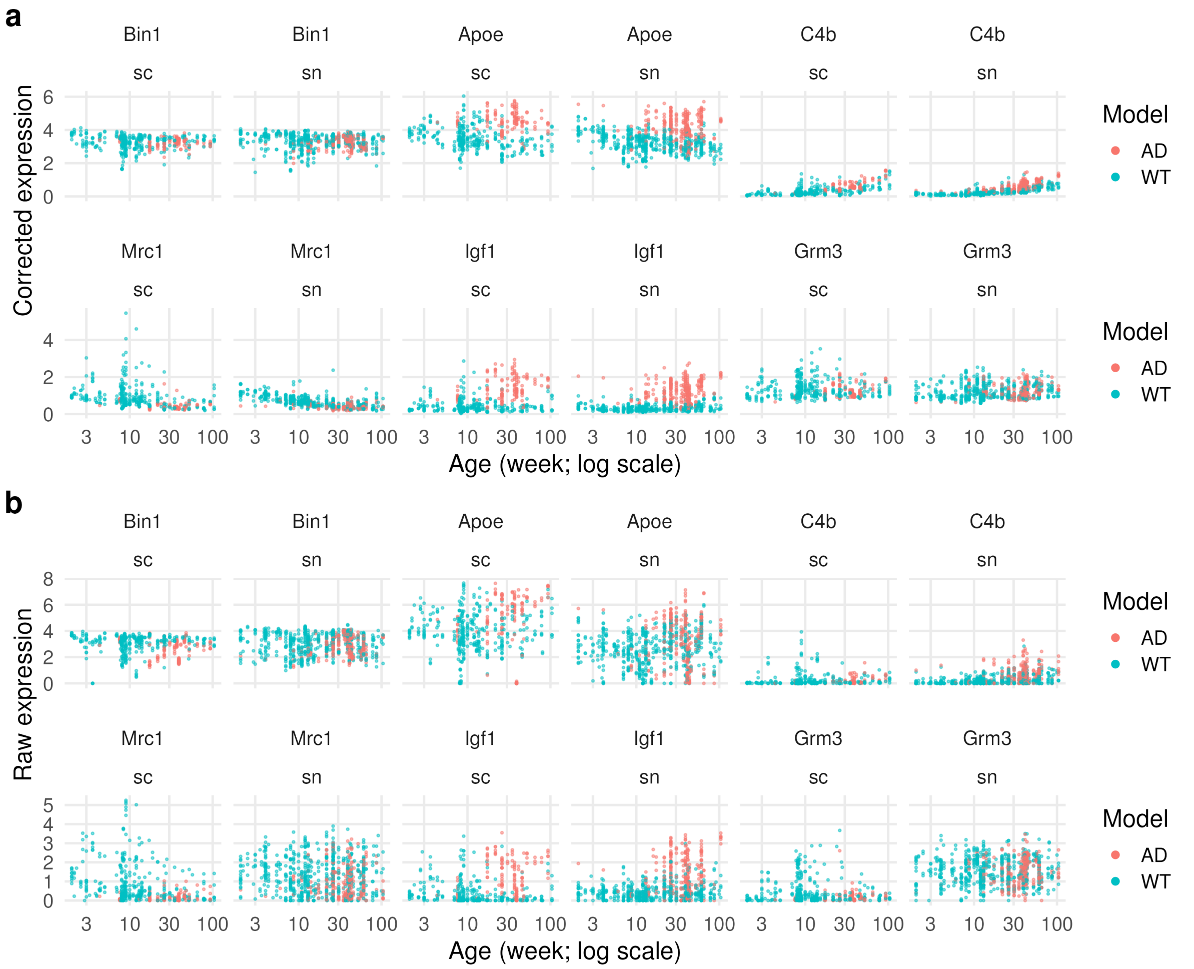


**Figure S14: Observed and adjusted expression of illustrative microglial genes across age.** Each point is a library, colored by Model and separated by gene and original sc/sn protocol. Age is shown on a log scale in weeks. (a) Library means of cell-level `R2` reconstructed expression from seed 0. (b) Observed-count pseudobulk, computed by summing raw counts within library before normalization and log transformation. All retained tissue groups are displayed. The normalization denominator is the 2,000-HVG set, but the order of averaging and log transformation differs between (a) and (b), so their absolute numerical scales should not be equated.

### Supplementary Note: exact LLM prompts

The generation-0 seed and selected generation-3 prompts are reproduced here, containing 2,379 and 15,265 characters, respectively. The `{METADATA}` placeholder was replaced by the assembled fields at inference time. The five examples in the selected prompt are synthetic training-derived records, not held-out test examples. The Claude instruction-revision and synthesis templates below contain the static instructions. The literal prompt text is retained even where its punctuation differs from the manuscript's editorial style.

#### Prompt S1: Gemma generation-0 seed

```
You are an expert curator of GEO (Gene Expression Omnibus) transcriptomics metadata.

Task: decide whether ONE GEO sample (GSM) should be labeled 1 or 0.

Label = 1 only if ALL three criteria hold for THIS sample:

(a) Assay: the sample was profiled by 10x Genomics single-cell RNA-seq or single-nucleus RNA-seq
   (e.g. Chromium droplet-based 3' or 5' gene expression, including Chromium-based multiome RNA).
   Not 10x: bulk RNA-seq, Smart-seq/Smart-seq2, Drop-seq, inDrop, SPLiT-seq, sci-RNA-seq, microarrays,
   spatial transcriptomics, ATAC-only, ChIP-seq, or other non-10x methods.
(b) Anatomy: the sample comes from hippocampus or cortex (any cortical area or hippocampal subregion,
   e.g. prefrontal, motor, somatosensory, visual, entorhinal cortex, CA1/CA3, dentate gyrus).
   Not target: whole brain without dissection, other brain regions (striatum, cerebellum, thalamus,
   hypothalamus, brainstem, spinal cord, retina, ...), non-brain tissues, cultured cell lines,
   iPSC-derived cells, or organoids.
(c) Condition: the sample itself is either a wild-type / non-transgenic / healthy control sample, or
   an Alzheimer's-disease-related sample (an AD model such as amyloid or tau transgenic or knock-in
   animals, or AD patient tissue). Not eligible: samples whose own condition is a different disease
   model or an unrelated genetic manipulation.

Rules:
- Judge the specific GSM sample. Sample-level fields (Characteristics, GSM_title, GSM_source,
 Extract_protocol, Treatment_protocol) take priority over the study-level description
 (Study_title, Overall_summary, Overall_design). A study may contain both eligible and ineligible samples.
- Use study-level information when sample-level fields are silent about a criterion.
- If a criterion is not supported by the available metadata, set that criterion to false.

Return ONLY one JSON object, with no other text, with exactly these keys:
{
  "assay_10x_sc_or_sn": true or false,
  "target_anatomy": true or false,
  "wt_or_ad_sample": true or false,
  "label": 1 or 0,
  "confidence": a number from 0.0 to 1.0,
  "failure_reasons": a list of short category labels for the criteria that failed (e.g. "non_10x", "wrong_tissue",
 ↪  "non_ad_mutant", "insufficient_evidence"); empty list if label is 1
}
"label" must be 1 only if all three criteria are true.

Metadata for this GSM sample:

{{METADATA}}
```

#### Prompt S2: selected Gemma generation-3 prompt

```
You are an expert curator of GEO (Gene Expression Omnibus) transcriptomics metadata.

Task: decide whether ONE GEO sample (GSM) should be labeled 1 or 0.

Label = 1 only if ALL three criteria hold for THIS sample.

(a) Assay: this sample's library is 10x Genomics single-cell or single-nucleus RNA-seq.
- Counts as 10x RNA:
  - Chromium droplet-based 3' or 5' gene expression;
  - the gene-expression (RNA) library of a 10x Multiome run.
- Not 10x: bulk RNA-seq, Smart-seq/Smart-seq2/v4, Drop-seq, inDrop, SPLiT-seq, sci-RNA-seq, microarrays, spatial
↪  transcriptomics, ATAC-only, ChIP-seq.
- A study-level statement that scRNA-seq/snRNA-seq was done with 10x is enough when the sample-level fields do not
↪  contradict it.
```

```
- Default when no platform is named: assume 10x (moderate confidence) if both of these hold:
  - the sample is single-cell or single-nucleus RNA-seq of dissociated cells or isolated nuclei, typically reporting thousands of cells or cell clusters;
  - no non-10x single-cell method is mentioned anywhere.
- Generic phrases such as "RNA libraries were prepared using standard Illumina protocols" are GEO boilerplate. They do NOT name a non-10x method and do not block the default.
- Studies with several library types: check which library THIS GSM is.
  - Spatial arm: if the study also has a spatial-transcriptomics arm but this GSM's title or Characteristics say snRNA-seq/scRNA-seq, judge it as a single-cell sample. The spatial arm does not make it non-10x. A GSM that is itself a spatial section is not 10x sc/sn.
  - Multiome ATAC: if the GSM is marked as the ATAC library of a multiome run, (a) is false.
  - Bulk and single-nucleus arms: use the sample's title, dissection and protocol to decide its arm. A finely dissected subregion sample with a bulk-RNA protocol is bulk.
  - 10x plus a non-10x single-cell method (e.g. Smart-seq) used on comparable samples: you need sample-specific proof that THIS GSM is 10x. Proof includes "10x", "Chromium", "droplet" or "3'-end scRNA-seq" in the title, Characteristics or a sample-specific protocol.
  - When the non-10x method was used only for bulk or low-input RNA-seq of sorted populations or cultured cells, and this GSM is a dissociated-tissue scRNA-seq sample, it is the 10x arm.
  - An Extract_protocol that merely lists both methods, or a GSM named only by a brain region or dissection code with no other cue, is NOT proof. Without proof, set (a) false and add "insufficient_evidence".

(b) Anatomy: the sample comes from hippocampus or cortex.
- Counts: any cortical area or hippocampal subregion. Examples are prefrontal, motor, somatosensory, visual, auditory, entorhinal and cerebral cortex, CA1/CA3, dentate gyrus, hippocampal formation, and embryonic or developing cortex.
- Also counts:
  - a cerebral hemisphere, cerebrum or forebrain sample that clearly includes cortex/hippocampus;
  - a generic "brain"/"CNS" tissue label in an AD or neurodegeneration study, when the protocol says regions of interest were excised or dissected.
- Not target:
  - whole brain without dissection;
  - other regions only (striatum, cerebellum, thalamus, hypothalamus, midbrain, brainstem, spinal cord, retina);
  - non-brain tissue, cell lines, iPSC-derived cells, organoids;
  - preparations enriched for blood vessels or microvessels (vascular fraction), rather than brain parenchyma cells or nuclei.

(c) Condition: the sample is WT/control or an Alzheimer's disease (AD)-related model.

Judge the condition of THIS sample, not the topic of the study. A study about stroke, epilepsy, surgery, prenatal exposure, behaviour, development or a gene of interest still contains eligible control samples.

Eligible (true):
- Wild-type, non-transgenic, healthy, sham-operated, vehicle-, saline- or untreated control animals, and +/+ or WT-allele littermate controls. This applies in any study, including non-AD disease studies.
- Animals that carry only neutral tool or labelling constructs, with no intended disease or gene-function change. Examples:
  - Cre or CreERT2 driver lines;
  - fluorescent reporter, Sun1-GFP, EGFP or BAC reporter lines;
  - Cre combined with the wild-type allele of the gene of interest;
  - viral tracing or labelling (rabies, AAV reporters);
  - reporter or plasmid libraries delivered by electroporation;
  - FACS sorting of labelled cells.
- Wild-type animals given somatic, local plasmid or gene delivery (e.g. in utero electroporation of a construct). These count when Characteristics give only a WT or outbred strain, tissue and age, and no mutant genotype or disease/drug treatment. The germline genotype stays wild type.
- Normal biological variables on WT animals: age, sex, developmental stage, brain region, cell-type sorting.
- AD-related samples:
  - amyloid, tau or APP/PS1-type transgenic or knock-in animals (e.g. 5xFAD, APP/PS1, APP23, P301S/PS19, TauPS2APP, App knock-in, App/MAPT double knock-in);
  - AD models or WT animals inoculated or seeded with tau, amyloid-β, or brain extracts from AD or tauopathy patients. Pathology seeding is an AD-related manipulation, not an experimental drug;
  - AD genetic-risk models: human APOE2/3/4 knock-in, including APOE knock-ins with Cre-mediated (conditional) APOE deletion, and TREM2 AD-risk variants. These count alone or crossed with an amyloid/tau model;
  - AD models crossed with a knockout, cKO or heterozygous deletion of a gene the study explicitly frames as part of an AD-risk pathway. An example is a microglial regulator of TREM2/DAP12 or ITIM signaling of AD GWAS genes;
  - such models given only vehicle/control treatment, or left untreated;
  - AD patient tissue.

Not eligible (false, "non_ad_mutant"):
- The experimental (non-control) arm of a non-AD disease, injury or manipulation model. Examples are ischemia/stroke, seizure or epilepsy mutants, surgery/anesthesia groups, alcohol or toxin exposure, infection or inflammation challenges, and drug-treated groups.
- Behaviourally trained experimental groups profiled as activity-tagged ("engram") cells, unless the sample is the home-cage or untrained control.
- Germline knockout, knock-in, cKO, overexpression or deletion of a non-AD gene, even when the study is framed around AD. This includes a non-AD gene knockout in otherwise WT animals within an AD-themed study.
```

```
- An AD model crossed with an additional gene knockout or cKO that the study does not connect to an AD-risk pathway.
- An AD model given an experimental drug, diet or compound intervention, for example when Characteristics name an
↪  inhibitor or agonist treatment. The added manipulation makes the sample no longer a plain AD-model sample.
↪  Tau/A-β/AD-brain seeding is not such an intervention.

How to read the sample's own arm:
- Sample-level fields take priority over study-level fields.
  - Sample-level fields: Characteristics, GSM_title, GSM_source, Extract_protocol, Treatment_protocol, Growth_protocol.
  - Study-level fields: Study_title, Overall_summary, Overall_design.
- A study may contain both eligible and ineligible samples. Read the genotype and treatment values in Characteristics
↪  carefully:
  - "sham", "saline", "vehicle", "Ctrl", "control", "+/+", "WT", a plain strain name (e.g. C57BL/6, CD1) or a
  ↪  driver/reporter-only genotype indicate a control;
  - an AD genotype with no treatment value, or with only a pathology-seeding treatment, indicates an AD-related sample.
- An explicit Characteristics treatment value overrides words in the GSM title. Titles often carry the experiment or
↪  batch name, such as a drug name, even for vehicle controls.
- Treatment_protocol and Growth_protocol text is often copied to every sample and may list all study arms. Do not
↪  assume this sample received a treatment listed there unless its title or Characteristics indicate it.

General rules:
- Use study-level information when sample-level fields are silent about a criterion. Do not reject a sample only
↪  because the study's disease is not AD.
- If a criterion is not supported by the available metadata, set it to false and add "insufficient_evidence". Do not
↪  guess eligibility from the study theme alone.
- Confidence:
  - high when sample-level fields explicitly settle all three criteria;
  - lower when you relied on study-level inference or a default.

Return ONLY one JSON object, with no other text, with exactly these keys:
{
  "assay_10x_sc_or_sn": true or false,
  "target_anatomy": true or false,
  "wt_or_ad_sample": true or false,
  "label": 1 or 0,
  "confidence": a number from 0.0 to 1.0,
  "failure_reasons": a list of short category labels for the criteria that failed (e.g. "non_10x", "wrong_tissue",
  ↪  "non_ad_mutant", "insufficient_evidence"); empty list if label is 1
}
"label" must be 1 only if all three criteria are true.

Labeled reference examples (short illustrative GSM records; use them to calibrate the rules above, do not copy their
↪  labels to the sample being classified):

### Example 1
Study_title: Activity of rapidly diverged primate regulatory sequences in embryonic mouse neocortex
Overall_summary: We compared fast-evolving noncoding regions across human, archaic hominin and ape genomes. We then
↪  tested 9 of these regions for enhancer activity in developing mouse brain using a pooled, single-cell readout of
↪  reporter transcripts. Several hominin alleles showed altered activity.
Overall_design: Reporter plasmid pools and a GFP co-transfection marker were introduced into embryonic mouse neocortex
↪  by in utero electroporation. After 20 hours, GFP+ cells were sorted and sequenced to capture transcriptomes and
↪  reporter-derived reads. Two independent batches were run, and the plasmid input was sequenced for normalization.
Characteristics: tissue: Neocortex genotype: C57BL/6J age: E13.5 electroporation; E14.5 collection treatment: in utero
↪  electroporation
GSM_title: Reporter output, batch B, E13.5 electroporation; single-cell RNA
GSM_source: Neocortex
Growth_protocol:
Treatment_protocol: Plasmid reporter pools were electroporated into embryonic cortex and the tissue was collected 20 h
↪  later.
Extract_protocol: GFP+ cortical pieces were dissociated with trypsin and DNase, triturated, filtered and stained for
↪  viability. Live GFP+ cells were FACS-sorted. Input plasmids were fragmented for DNA library prep. Single-cell
↪  libraries were prepared with the Chromium Next GEM Single Cell 3' v3.1 kit (10x Genomics) using GEM generation,
↪  barcoded RT, cDNA amplification and SPRI size selection. scRNA-seq/DNA-seq
Correct final label: 1

### Example 2
Study_title: Microglial subsets shaping memory recovery after chronic cerebral hypoperfusion
Overall_summary: We profiled hippocampal cells from mice after carotid stenosis and from sham-operated mice. About
↪  19,500 cells were assigned to 11 neural, glial and vascular populations, and condition-specific changes were
↪  examined.
Overall_design: Hippocampi from 3 sham and 3 bilateral carotid stenosis mice
Characteristics: age: 10 weeks strain: C57BL/6 treatment: sham tissue: hippocampus
GSM_title: sham operated replicate 3
GSM_source: hippocampus
Growth_protocol:
Treatment_protocol:
```

```
Extract_protocol: Hippocampal tissue was minced and digested at 37°C for 25 min in Hibernate A/B27 medium containing
↪  pronase and DNase I with gentle shaking. The digest was triturated with fire-polished tips, and debris was removed
↪  by density gradient centrifugation. The cells were washed in PBS with 1% BSA, pelleted at low speed and resuspended
↪  at the target concentration. Sequencing libraries were prepared with standard Illumina protocols.
Correct final label: 1

### Example 3
Study_title: Projection neuron identity tunes the abundance of matched cortical inhibitory subtypes
Overall_summary: Cortical inhibitory subtypes pair with specific excitatory classes. In Tbr1-deficient cortex, layer 6
↪  corticothalamic neurons are lost and other projection types expand. We find that the associated interneuron
↪  subtypes shift in proportion, either through altered survival or through a change in identity.
Overall_design: We genetically altered projection neuron fate and profiled the interneuron nuclei that were labeled
↪  through a Cre-dependent nuclear reporter.
Characteristics: tissue: Whole cortex genotype: Gad2-Cre;Tbr1 flox/flox;Sun1-GFP
GSM_title: P9 cortical interneurons Tbr1 cKO
GSM_source: Whole cortex
Growth_protocol:
Treatment_protocol:
Extract_protocol: Cortex was dissected into cold sucrose homogenization buffer and Dounce-homogenized in buffer
↪  containing detergent and RNase inhibitor. Nuclei were filtered, pelleted, resuspended in PBS/BSA and stained with
↪  DRAQ5. GFP+ nuclei were then sorted. Libraries were made with the Chromium Single Cell 3' v3.1 kit (10x Genomics)
↪  and sequenced on a NovaSeq. snRNA-seq
Correct final label: 1

### Example 4
Study_title: Multimodal atlas of amyloid-associated cell states in APP/PS1 mouse brain
Overall_summary: We combined nucleus-level and spatial transcriptomics from the same animals to map early
↪  amyloid-related changes. Plaque-adjacent astrocytes, white-matter oligodendrocytes and region-specific excitatory
↪  neurons showed altered metabolic and synaptic programs.
Overall_design: Ten wild-type and ten APP/PS1 mice were analyzed at 8 months. In four mice per group, the brain was
↪  split into hemispheres. One hemisphere was used for snRNA-seq. The other hemisphere was sectioned coronally through
↪  the hippocampus at anterior, middle and posterior levels for immunostaining and spatial transcriptomics.
Characteristics: tissue: APP/PS1 mouse brain Sex: female age: 8 months genotype: APP/PS1
GSM_title: APP/PS1 brain, snRNA-seq, replicate 2
GSM_source: APP/PS1 mouse brain
Growth_protocol: Animals were kept on a 12 h light cycle with free access to chow and water
Treatment_protocol: Transcardial PBS perfusion was performed, and the brains were then bisected.
Extract_protocol: Hemispheres for histology were fixed in 4% PFA. Hemispheres for snRNA-seq were flash-frozen in
↪  chilled isopentane. RNA libraries were prepared with standard Illumina protocols. Spatial Transcriptomics
Correct final label: 1

### Example 5
Study_title: Wnt ligand overexpression redirects cortical progenitor output [scRNA-seq]
Overall_summary: Purpose: To compare single-cell transcriptomes of control E17.5 cortex with cortex electroporated with
↪  a Wnt3a plasmid at E14.5. Methods: Cortices were dissected, digested with papain, filtered and washed to obtain
↪  single cells. Libraries were generated on the 10X Genomics Chromium droplet platform and sequenced on a HiSeq.
↪  Results: About 6,900 control cells and 7,700 electroporated cells were recovered.
Overall_design: scRNA-seq of single cells from E17.5 control and Wnt3a-IUE mouse cortices.
Characteristics: strain: CD1 tissue: Cortex developmental stage: Embryonic day 17.5
GSM_title: Wnt3a-IUE Cortex E17.5_B2
GSM_source: Wnt3a IUE E17.5 Cortex
Growth_protocol:
Treatment_protocol: The Wnt3a plasmid was electroporated in utero at E14.5. Cortices were dissected at E17.5 under
↪  fluorescence guidance, then papain-digested, triturated and filtered into a single-cell suspension.
Extract_protocol: scRNA-seq libraries were generated on the 10X Genomics Chromium droplet system following the
↪  manufacturer's instructions.
Correct final label: 1

Metadata for this GSM sample:

{{METADATA}}
```

### Prompt S3: Claude instruction-revision template

```
CLAUDE SYSTEM PROMPT (--system-prompt):
You are an expert prompt engineer optimizing a classification prompt that another language model will execute. Answer
↪  only in the requested tagged format.

INSTRUCTION-REVISION REQUEST: FIXED TEMPLATE
You are revising the INSTRUCTION part of a classification prompt for another language model (Gemma).

Do not classify the samples yourself. Revise the instruction so that Gemma makes fewer errors of

the kinds shown below. All examples and metrics below come from the TRAINING split only.
```

```
Rules:

- Analyse WHY Gemma made the errors (e.g. study-level vs sample-level confusion, 10x vs non-10x,

  missed hippocampus/cortex subregions, missed WT/control samples, AD model vs non-AD mutant,

  over-guessing from sparse metadata) and fix the rules responsible. Preserve rules that work.

  Do not simply make the instruction longer.

- Write general rules. Do NOT copy sample-specific text, study names, GSE/GSM identifiers or URLs.

- Do NOT include any labeled examples in the instruction. A separate block of short synthetic labeled

  examples (modelled on some of these train errors) is appended automatically after the instruction, followed by the metadata of the sample to classify.

- Do NOT include the placeholder {{METADATA}} or a "Metadata for this GSM sample" header.

- Keep the output contract unchanged: Gemma must return ONLY one JSON object with exactly these keys:

  "assay_10x_sc_or_sn" (bool), "target_anatomy" (bool), "wt_or_ad_sample" (bool), "label" (0/1),

  "confidence" (0.0-1.0), "failure_reasons" (list of short category labels such as "non_10x",

  "wrong_tissue", "non_ad_mutant", "insufficient_evidence").

- Metadata fields available to Gemma: Study_title, Overall_summary, Overall_design, Characteristics,

  GSM_title, GSM_source, Growth_protocol, Treatment_protocol, Extract_protocol.

- English. Keep it under about 9000 characters. No optimization commentary inside it.

Respond in exactly this format:

<change_summary>

2-5 short bullet points describing what you changed and why.

</change_summary>

<instruction>

(the complete revised instruction)

</instruction>

## Task the instruction must solve
Label = 1 only if THIS GSM sample (a) was profiled by 10x Genomics single-cell or single-nucleus RNA-seq, (b) comes from hippocampus or cortex (or a subregion), and (c) is itself WT/control or an Alzheimer-disease-related model. Sample-level metadata takes priority over the study description.

## Current prompt on the TRAIN split (n=<N>, positive class = label 1)
accuracy=<ACCURACY> F1=<TRAIN_F1> precision=<PRECISION> recall=<RECALL> TP=<TP> FP=<FP> TN=<TN> FN=<FN> parse_errors=<N_PARSE_ERRORS>

## Train-metric history (train split only)
<UP_TO_FIVE_PREVIOUS_GENERATIONS_AND_CHANGE_SUMMARIES>

## Criterion-level breakdown of train errors (Gemma's own booleans)
<COUNTS_OF_FALSE_NEGATIVES_AND_FALSE_POSITIVES_BY_EACH_OF_THREE_CRITERIA>

## Train FALSE NEGATIVES (label 1, Gemma did not output 1) [<N_SHOWN> of <N_ALL>]
<UP_TO_EIGHT_TRAIN_FALSE_NEGATIVES_WITH_METADATA_AND_GEMMA_OUTPUT>

## Train FALSE POSITIVES (label 0, Gemma output 1) [<N_SHOWN> of <N_ALL>]
<UP_TO_EIGHT_TRAIN_FALSE_POSITIVES_WITH_METADATA_AND_GEMMA_OUTPUT>

## Current instruction (to revise)
<current_instruction>
<PRECEDING_GENERATION_INSTRUCTION>
</current_instruction>
```

#### Prompt S4: Claude synthesis template

```
CLAUDE SYSTEM PROMPT (--system-prompt):
You are an expert prompt engineer optimizing a classification prompt that another language model will execute. Answer only in the requested tagged format.

FEW-SHOT SYNTHESIS REQUEST: FIXED TEMPLATE
You are writing SYNTHETIC few-shot examples for a classification prompt executed by another model.

Each source record below is a real GEO sample (GSM) metadata record that the classifier got wrong.

For EACH source record write ONE synthetic, analogous, SHORT metadata record that:

- keeps the same ground-truth label (given; do not change it) and the same classification-relevant

  evidence pattern: the same assay cues, the same anatomy cues, the same genotype/condition cues, the

  same ambiguity, and the same split between sample-level and study-level fields (if a cue is only

  in the study description, or only in one field, keep it there; if a field is empty, leave it empty);

- has the same difficulty: do NOT add cues that are absent from the source to make the answer easier,

  and do NOT remove the misleading cues that made it hard;

- changes every incidental detail: rephrase all text in your own words, invent different sample

  names, study topic details, labs, numbers, ages and replicate IDs; never copy phrases of more than

  a few words from the source;

- contains no GSE/GSM/SRA/accession identifiers, no URLs, no author names;

- is short: about 600-1200 characters in total over all fields.

Fields (all must be present as strings, possibly empty): Study_title, Overall_summary, Overall_design, Characteristics, GSM_title, GSM_source, Growth_protocol, Treatment_protocol, Extract_protocol

Respond with ONLY a JSON array, one object per source in the same order, each of the form

{"source": <source number>, "fields": {<field>: <string>, ...}}

## Task definition
Label = 1 only if THIS GSM sample (a) was profiled by 10x Genomics single-cell or single-nucleus RNA-seq, (b) comes from hippocampus or cortex (or a subregion), and (c) is itself WT/control or an Alzheimer-disease-related model. Sample-level metadata takes priority over the study description.

## Source records (training split)
### Source <NUMBER> (ground-truth label <0_OR_1>; classifier wrongly gave <PREDICTION>)
<SOURCE_TRAINING_GSM_METADATA_TRUNCATED_TO_6000_CHARACTERS>
[REPEATED FOR UP TO_FIVE_POSITIVE_AND_FIVE_NEGATIVE_TRAIN_ERRORS]
```

#### Prompt S5: preliminary GPT-4o requests

```
GPT-4o PRIMARY METADATA EXTRACTION (exact prompt strings from supplied code)
Model: gpt-4o; temperature: 0. Seven requests per GSM.

A. Region, Sex, Age, and Strain: four separate requests. Each request uses
   description1, made by concatenating nonempty strings from columns 9 and 11
   of the source dataframe (zero-based pandas iloc), separated by "\n ".

Message 1 (role=user):
Read following descriptions and answer user's questions. Here is the descriptions: {description1}
Message 2 (role=assistant):
I'm, ready
Message 3 (role=user): one of the following, sent separately:

REGION:
Which brain region the biological sample came from? Provide only the unique region, like ' cortex' including subregion if it's provided. If it came from more than one region, like cortex and hippocampus, provide like 'Cortex;Hippocampus'. If it is ambiguous, like brain, provide 'unknown'

SEX:
Extract sex (Female/Male/Mix/Unknown). Without any rationale, provide the information like 'Male'
```

```
AGE:
Extract age (with unit, like 2-months, 3-weeks, unknown). Without any rationale, provide the information like
↪  '2-months'

STRAIN:
Extract strain (like C57BL/6J, C57BL/6N, unknown). Without any rationale, provide the information like 'C57BL6J'

B. Sequencing modality and Genotype/Treatment: two separate requests.
   These use description2, made by concatenating nonempty strings from columns
   9, 11, 12, 13, and 14 of the same dataframe, separated by "\n ".

Message 1 (role=user):
Read following descriptions and answer user's questions. Here is the descriptions: {description2}
Message 2 (role=assistant):
I'm, ready
Message 3 (role=user): one of the following, sent separately:

SEQUENCING STRATEGY / SCSN / PLATFORM:
Extract the sequencing strategy (like bulk-RNA-seq, ATAC-seq, sc/snRNA-seq, unknown), single-cell or single-nucleus
↪  RNA-seq (like sc, sn, unknown) and whether the sc/snRNA-seq sample comes from 10X Genomics/Chromium platform or not
↪  (like 10X, other, unknown). Without any rationale, provide the information like 'Strategy: sc/snRNA-seq; scsn: sc;
↪  Platform: 10X'

GENOTYPE / TREATMENT:
Extract genotype (like WT, 5xFAD, unknown) and treatment (like Tamoxifen, Water restriction, unknown). Without any
↪  rationale, provide the information like 'Genotype: WT; Treatment: unknown'

C. Model: one request. The model-answer stage only gets an extracted-field
   string, not the original GEO metadata, constructed exactly as:
   description3 = "Strain: " + strain_answer + "; Genotype: " +
                  genotype_treatment_answer[0] + "; Treatment: " +
                  genotype_treatment_answer[1]

Message 1 (role=user):
What is the mouse experimental model based on this: {description3} ?. Without any rationale, provide only a label
↪  like'WT', 'Obesity', 'Alzheimer's disease', 'High calory'
Message 2 (role=assistant):
I'm, ready

NOTE: The unusual wording and assistant acknowledgments above reflect the
supplied script verbatim. The model request has the assistant message last.
The supplied primary example calls process_df(df[0:2]); it does not establish
how the entire earlier cohort was processed or what caused its failures.
```

#### References


[1] Damian Smedley, Syed Haider, Benoit Ballester, Richard Holland, Darin London, Gudmundur Thorisson, and Arek Kasprzyk. BioMart – biological queries made easy. *BMC Genomics*, 10(1):22, 2009. doi: 10.1186/1471-2164-10-22. URL https://doi.org/10.1186/1471-2164-10-22.

[2] John N. Hutchinson, Alexander W. Ensminger, Christine M. Clemson, Christopher R. Lynch, Jeanne B. Lawrence, and Andrew Chess. A screen for nuclear transcripts identifies two linked noncoding RNAs associated with SC35 splicing domains. *BMC Genomics*, 8:39, 2007. doi: 10.1186/1471-2164-8-39. URL https://doi.org/10.1186/1471-2164-8-39.

[3] Tim Stuart, Andrew Butler, Paul Hoffman, Christoph Hafemeister, Efthymia Papalexi, William M. Mauck, III, Yuhan Hao, Marlon Stoeckius, Peter Smibert, and Rahul Satija. Comprehensive Integration of Single-Cell Data. *Cell*, 177(7):1888–1902.e21, 2019. doi: 10.1016/j.cell.2019.05.031. URL https://doi.org/10.1016/j.cell.2019.05.031.

[4] Ilya Korsunsky, Nghia Millard, Jean Fan, Kamil Slowikowski, Fan Zhang, Kevin Wei, Yuriy Baglaenko, Michael Brenner, Po-ru Loh, and Soumya Raychaudhuri. Fast, sensitive and accurate integration of single-cell data with Harmony. *Nature Methods*, 16(12):1289–1296, 2019. doi: 10.1038/s41592-019-0619-0. URL https://doi.org/10.1038/s41592-019-0619-0.

[5] Gilad Sahar Green, Masashi Fujita, Hyun-Sik Yang, Mariko Taga, Anael Cain, Cristin McCabe, Natacha Comandante-Lou, Charles C. White, Anna K. Schmidtner, Lu Zeng, Alina Sigalov, Yangling Wang, Aviv Regev, Hans-Ulrich Klein, Vilas Menon, David A. Bennett, Naomi Habib, and Philip L. De Jager. Cellular communities reveal trajectories of brain ageing and Alzheimer's disease. *Nature*, 633(8030):634–645, 2024. doi: 10.1038/s41586-024-07871-6. URL https://doi.org/10.1038/s41586-024-07871-6.

[6] Emma Gerrits, Nieske Brouwer, Susanne M. Kooistra, Maya E. Woodbury, Yannick Vermeiren, Mirjam Lambourne, Jan Mulder, Markus Kummer, Thomas Möller, Knut Biber, Wilfred F. A. den Dunnen, Peter P. De Deyn, Bart J. L. Eggen, and Erik W. G. M. Boddeke. Distinct amyloid-$\beta$ and tau-associated microglia profiles in Alzheimer's disease. *Acta Neuropathologica*, 141(5):681–696, 2021. doi: 10.1007/s00401-021-02263-w. URL https://doi.org/10.1007/s00401-021-02263-w.

[7] Hadas Keren-Shaul, Amit Spinrad, Assaf Weiner, Orit Matcovitch-Natan, Raz Dvir-Szternfeld, Tyler K. Ulland, Eyal David, Kuti Baruch, David Lara-Astaiso, Beata Toth, Shalev Itzkovitz, Marco Colonna, Michal Schwartz, and Ido Amit. A Unique Microglia Type Associated with Restricting Development of Alzheimer's Disease. *Cell*, 169(7):1276–1290.e17, 2017. doi: 10.1016/j.cell.2017.05.018. URL https://doi.org/10.1016/j.cell.2017.05.018.

[8] Emil Uffelmann, Douglas P. Wightman, et al. Alzheimer's disease genetics converges on microglial and neuronal dysregulation. *Nature*, 2026. doi: 10.1038/s41586-026-11206-y. URL https://doi.org/10.1038/s41586-026-11206-y.

[9] LRAPA datasets contributors. AD-associated genes reference list. https://github.com/yalanyang/LRAPA_datasets/blob/main/reference/genelist/AD_associated_genes.txt. GitHub repository. Accessed: 2026-11-03.

[10] ADSP Gene Verification Committee. Ad/adrd top genes repository. https://topgenes.niagads.org/. Accessed: 2026-11-03.
